# Selection bias in microbial mutation accumulation studies and the impact of colony growth

**DOI:** 10.64898/2026.05.28.725302

**Authors:** Julie M. Grosse-Sommer, Jarrod D. Hadfield

## Abstract

In microbial mutation-accumulation (MA) studies, it is widely thought that natural selection is silenced by repeated single-cell bottlenecks. In support of this claim, 40 published tests for selection bias failed to detect a deficit of non-synonymous relative to synonymous mutations in wild-type microbes. Here we show that this most likely reflects selective reporting and lack of power. Our meta-analysis of 10,856 mutations from wild-type microbial MA reveals a clear signal of selection: non-synonymous mutations are observed 7.7% less often than synonymous mutations. However, our inference of the bias is hampered by a widespread failure to consider the mutation spectrum. To overcome this, we provide a multinomial-logit model that jointly estimates the mutation spectrum and selection. By applying this to a 194-line *Escherichia coli* MA experiment and five previous *E. coli* datasets (869 mutations) we reveal a deficit of non-synonymous mutations, although the reduction is not significant. While approaches do exist for correcting for selection bias, all currently assume that microbial MA lines are grown in well-mixed liquid culture rather than as surface colonies where competition is spatially structured. Although existing theory suggests selection bias should be stronger under colony growth, using agent-based simulations we show that this actually depends on the scale over which neighbouring cells compete and how unevenly they divide: it can be weaker, equivalent to, or stronger than in homogeneous growth. While our preliminary assessment is that it is considerably stronger, quantitative predictions will require the empirical details of colony growth to be better resolved.

## Introduction

Many fundamental concepts in evolutionary genetics, such as the evolution of sex (Peck et al. 1997), mutation load (Lesecque et al. 2012), adaptation (MacLean and Buckling 2009) and the maintenance of quantitative genetic variation (Walsh and Lynch 2018, Chapter 28) require information about the properties of new mutations. However, sampling new mutations without bias remains difficult as their fitness effects often influence their probability of being observed. Mutation accumulation (MA) lines address this issue by maintaining populations at very low effective population sizes (*N*_*e*_), ensuring the fixation probability of any newly arising mutation is dominated by drift rather than selection (Muller 1928; Mukai 1964). In some cases, *N*_*e*_ can be kept as low as one, for example in species that can reproduce through selfing (Keightley and Caballero 1997) or parthenogenesis (Lynch et al. 1998), or where genetic constructs can be used to passage single chromosomes (Mukai 1964). For obligate out-crossing species, however, *N*_*e*_ can only be kept as low as two by propagating lines via sib mating (Katju and Bergthorsson 2019; Fernández and López-Fanjul 1996). In microbes, MA lines are often serially bottlenecked using colony propagation (Kibota and Lynch 1996; Andersson and Hughes 1996; Zeyl and DeVisser 2001) (but see Krašovec *et al*. (2017a), Robert *et al*. (2018) and Baehr *et al*. (2025) for alternative methodologies) where the number of cells in a colony can be as large as 10^8^ between bottlenecks (Mahilkar et al. 2022; Baehr et al. 2025). If these large population sizes result in a large *N*_*e*_, selection may be effective at filtering out deleterious mutations and enriching for beneficial mutations, potentially skewing inferences about mutation rates and the distribution of fitness effects of new mutations. Given that even small increases in *N*_*e*_ have been shown to have dramatic effects in MA lines of *Caenorhabditis elegans* (Katju *et al*. 2015), it is crucial to determine *N*_*e*_ in microbial MA lines and assess the prevalence of selection.

In microbial MA, the harmonic mean census population size has traditionally been used to estimate *N*_*e*_ (Karlin 1968), assuming that the population size doubles each generation between bottlenecks. For *τ* population doublings between bottlenecks, the harmonic mean population size is approximately (*τ* + 1)/2 such that estimates of *N*_*e*_ typically range between five (Ness *et al*. 2012) and 17 (Chen *et al*. 2021) (see Table 1). Since selection is only expected to have a substantial effect on mutations with absolute selection coefficients (*s*) greater than 1/*N*_*e*_, it is widely believed that the large majority of mutations are likely fixed irrespective of their fitness effect (e.g. Dillon and Cooper 2016). While the relevance of the harmonic-mean effective population size for fixation probabilities has been queried (Wahl and Agashe 2022), simulations of microbial MA lines have shown that the decline in mean fitness is similar in lines of constant size and those of fluctuating population sizes with an equivalent harmonic mean population size, suggesting that it is a good metric for quantifying the effectiveness of selection (Gordo and Dionisio 2005). However, reservations about its accuracy remain since the model and the simulations of Gordo and Dionisio (2005) assume, among other things, a Poisson number of offspring which may be unrealistic in populations that grow by binary fission with little cell death (Wahl and Agashe 2022).

**Table 1.**
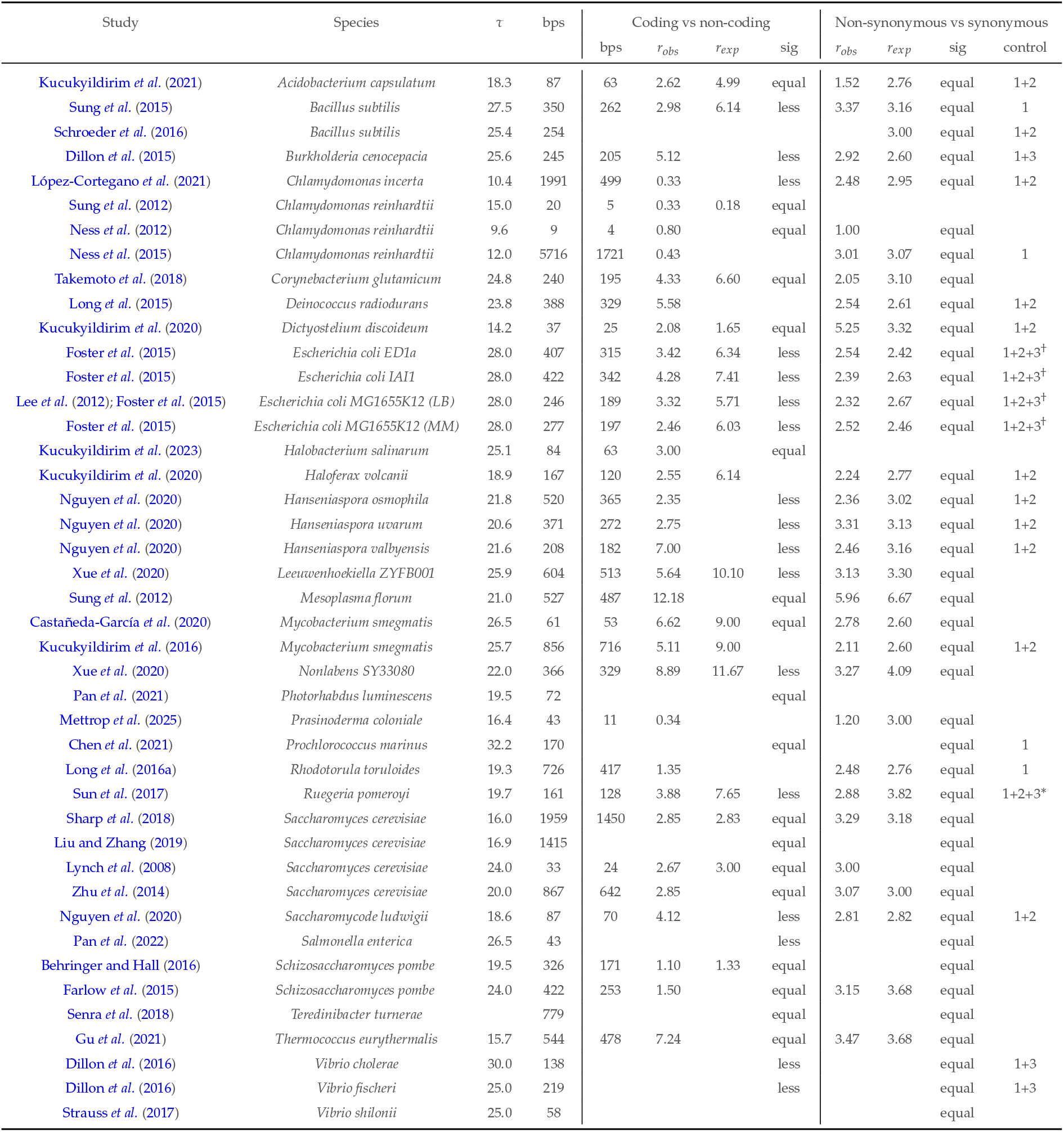
Signatures of selection in microbial MA line studies that have propagated lines using colony growth and used wild-type strains. *τ* is the number of population doublings between bottlenecks. ‘bps’ are the number of base-pair substitutions either in total or in coding regions. *r*_*exp*_ and *r*_*obs*_ are the expected and observed ratios for coding versus non-coding or non-synonymous versus synonymous (NS/S) mutations. ‘sig’ indicates whether the observed ratio was significantly different from expected (less or greater) or not (equal). When testing NS/S, the control column indicates what was controlled for: codon usage (1), transition/transversion ratio (2) and GC bias (3). ^†^ indicates simulation was used to assess expectations and ^*\**^ indicates the method of Yang and Nielsen (2000) was used. In many cases the method was not explicitly stated. For studies included in the meta-analysis, see Methods for discrepancies between published data and those reported in the table. See Results for discrepancies in significance tests. In addition, in Chen *et al*. (2021), 79 of the 170 bps fell into a single gene, all of which were non-synonymous - tests of neutrality either excluded mutations in this gene, or lines containing mutations in that gene. Pan *et al*. (2022) used 15 wild-type lines and tested for neutrality on each line separately - the bps in the table is the average number of bps per line. Strauss *et al*. (2017) performed MA in four environments and tested for neutrality in each environment.

While the concept of *N*_*e*_ has been used to give heuristic insight into how effective selection might be in microbial MA, quantitative assessments of selection bias in microbial MA have typically utilised deterministic models of selection under exponential population growth (Kibota and Lynch 1996; Joseph and Hall 2004; Wahl and Agashe 2022). Kibota and Lynch (1996) derived the odds of sampling a mutation with selection coefficient *s* versus an unmutated, ancestral cell, and used this result to revise their estimate of the lower-bound on the average selection coefficient in *Escherichia coli* from −0.012 to −0.013. Using a very similar model derived by Otto and Orive (1995) for exploring somatic selection, Joseph and Hall (2004) used a maximum-likelihood framework for correcting their estimate of the distribution of fitness effects. The bias correction reduced the estimate of the proportion of mutations that are beneficial in their *Saccharomyces cerevisiae* MA study from 19% to 5.75%, although uncertainty was high. Wahl and Agashe (2022) approximated the model of Kibota and Lynch (1996) by assuming the mutation rate is negligible compared to the growth rate, and derived the odds of sampling a mutation with selection coefficient *s* versus a neutral mutation (*s* = 0). The odds only depends on *s* and *τ* and was incorporated into a heuristic procedure for reweighting the distribution of fitness effects of sampled mutations. Both Wahl and Agashe (2022) and Otto and Orive (1995) complemented their deterministic models with stochastic simulations and found them to be accurate. Mahilkar *et al*. (2022) also used simulations to study the amount of bias in microbial MA, and although a method for correcting the bias was not developed, it was noted that the bias may be large, with beneficial mutations being over-represented by more than a factor of two, a sentiment echoed by Wahl and Agashe (2022).

To our knowledge, there is no experimental verification of whether the predicted selection bias is quantitatively accurate. However, Mahilkar *et al*. (2022) tested for the presence of selection bias in colony-propagated *E. coli* MA by setting up three sets of MA lines that were bottlenecked after 8, 16 or 24 hours of growth. Contrary to prediction, the lines that experienced the most growth between bottlenecks had faster fitness declines per generation despite the expectation that selection would be more efficient at removing deleterious mutations1^1^. While this result indicates that selection bias may b minimal, MA lines propagated from cells taken from the colony periphery had a higher growth rate than lines propagated from cells taken from the colony centre, implying the presence of selection (Mahilkar et al. 2022).

Indirect tests of selection bias in microbial MA are more widespread. Given that non-synonymous sites are under the strongest purifying selection, followed by non-coding and synonymous coding sites (Thorpe et al. 2017), a number of studies have explored whether there is a deficit of non-synonymous mutations compared to expectation, indicative of selection. In a review of the literature for naturally occurring mutations from colony-based microbial MA lines (Table 1), we could not find a single study that reported a significant difference between the expected and observed ratio of non-synonymous versus synonymous mutations, consistent with weak or no selection bias. Similarly, tests of whether the observed number of non-sense mutations is less than expected have failed to find a difference (Lee *et al*. 2012). While some studies also find no difference between the observed and expected ratio of coding versus non-coding mutations (Lynch *et al*. 2008; Sung *et al*. 2012; Ness *et al*. 2012; Zhu *et al*. 2014; Farlow *et al*. 2015), other studies have found a deficit of observed coding mutations, consistent with purifying selection (Foster *et al*. 2015; Sung *et al*. 2015; Sane *et al*. 2018; López-Cortegano *et al*. 2021).

To summarise, while theory often predicts appreciable selection bias in microbial MA, the evidence for it is mixed. However, a major assumption of the theory, and the subsequent methods developed from it to correct for selection bias, is that growth between bottlenecks is exponential, mimicking well-mixed, homogeneous growth in liquid culture (Kibota and Lynch 1996; Joseph and Hall 2004; Mahilkar *et al*. 2022; Wahl and Agashe 2022). In contrast, microbial MA nearly always involves colony propagation, where population growth predominantly occurs at the colony periphery as nutrients are depleted centrally (Hallatschek *et al*. 2007; Farrell *et al*. 2017; Nadell *et al*. 2016). There are several reasons why the efficiency of selection may be different under colony growth compared to homogeneous growth. First, only the subset of cells on or close to the colony periphery divide, meaning *N*_*e*_ may be much smaller than the total number of cells in the colony, reducing selection efficacy compared to homogeneous growth (Hallatschek *et al*. 2007). Second, the number of generations required to achieve the same final population size is higher under colony growth than for homogeneous growth, allowing more time for selection to alter mutation frequencies, potentially increasing selection bias (Gralka *et al*. 2016). Finally, neighbouring cells are more likely to carry the same mutation which will reduce the efficiency of selection under kin-competition or increase it under kin-cooperation (Nadell *et al*. 2016). Empirical observations show that *S. cerevisiae* mutants with a selective advantage of 0.15, seeded at an initial frequency of 2%, achieve a frequency of 50% under colony growth compared to 6% under homogeneous growth, strongly suggesting increased selection efficiency in colony growth (Gralka *et al*. 2016). Baehr *et al*. (2025) compared liquid-based and colony-based MA procedures in *E. coli* and found that mutations accumulated through colony-based MA were 7.6% less deleterious and had a 3% reduction in the odds of being non-synonymous versus synonymous. Although consistent with selection being more efficient in colony growth, the differences were far from significant. In contrast, Lavrentovich *et al*. (2016) showed that selection is less efficient under colony growth in *S. cerevisiae* when studying the outcome of mutation-selection balance: for a given mutation rate and *s*, new deleterious alleles have higher equilibrium frequency under colony growth. Similarly, Bosshard *et al*. (2017) showed that in mutator strains of *E. coli* subject to spontaneous mutations, fitness declined under colony growth yet increased under homogeneous growth. These striking differences to homogeneous growth suggest that existing correction methods for selection bias in MA using colony growth may be quantitatively inaccurate.

Another explanation for discrepancies between theory and empirical evidence is that existing models for quantifying selection bias are accurate, but attempts to test for it by looking for signatures in mutational patterns are underpowered or bi-ased. Many studies may lack sufficient statistical power if the number of mutations accumulated is small and/or the difference between the selection coefficients of non-synonymous and synonymous mutations is small. In addition, the rate and type of mutation are often dependent on the mutational spectra, and disregarding this dependence may mask signatures of selection or lead them to be falsely identified. For example, an excess of INDELs in non-coding versus coding regions of *Bacillus subtilis* might have been mistakenly ascribed to purifying selection during the MA process, rather than because non-coding sequences contain more homopolymer runs than coding sequences (Schroeder et al. 2016), which tend to be susceptible to INDELs (Schroeder et al. 2016; Lee et al. 2012; Foster et al. 2015; Shewaramani et al. 2017). Although some studies (see Table 1) control for certain aspects of the mutational spectra, such as codon usage bias (Sung et al. 2015), transition/transversion rates (Long *et al*. 2015) and GC bias (Dillon et al. 2015), very few control for the full mutational-spectra (Lee et al. 2012).

In this paper we use simulations to assess whether the predicted selection bias from homogeneous-growth models (Wahl and Agashe 2022; Joseph and Hall 2004; Kibota and Lynch 1996) is accurate under colony growth. We find that the amount of selection bias in colonies can be much stronger, or much weaker, than is observed under homogeneous growth, depending on the spatial scale of competition and the degree of asymmetry in cell division rates. Meta-analysing the expected and observed non-synonymous to synonymous ratios from 29 previously published MA studies shows a clear deficit of non-synonymous mutations, consistent with purifying selection. We also conducted an MA experiment in *E. coli*, and pooled our mutation data with published data from other *E. coli* MA studies, making this one of the most powerful microbial dataset for looking at mutational signatures of selection (869 mutations and *τ* = 28). Using a new multinomial model that corrects for mutational spectra, we detect a comparable deficit of non-synonymous mutations, but not significantly so. The deficit is most probably larger than we would predict from homogeneous-growth models of selection bias given current knowledge on the selection coefficients of non-synonymous and synonymous mutations. However, we tentatively suggest that homogeneous-growth models should be used as an approximate correction for selection bias until colony-growth models are more realistic and better parameterised.

## Materials and Methods

### Homogeneous versus colony-based models of growth

Theoretical studies have examined the amount of selection bias in MA lines when cell division probability is spatially uniform between bottlenecks (Kibota and Lynch 1996; Wahl and Agashe 2022; Joseph and Hall 2004). We refer to these as homogeneous-growth models. However, MA lines in microbes are typically propagated as surface colonies (Baehr et al. 2025) and so here we study the amount of selection bias that occurs when colony growth is explicitly modelled with spatial dynamics.

The Eden model (Eden 1961) serves as a basis for modelling colony growth where cells live on a 2-D lattice and can produce ‘daughters’ in empty adjacent sites (Figure 1). In the absence of cell death, cells are only added at the colony boundary. A perimeter site is defined as a site that is empty but is adjacent to an occupied site. If cell death is not allowed, an occupied site can have up to three perimeter sites, rather than four, since one site is occupied by the cell’s ‘mother’. Any occupied site with a free perimeter site is considered an edge site. When choosing which cell reproduces and where its daughter is placed, either a cell-based or a site-based rule can be implemented (Colyer et al. 2024). In a cell-based rule system with selection, an edge cell is sampled from all edge cells proportional to their fitnesses minus one (*w−* 1), and a daughter cell is placed in a random adjacent perimeter site to the selected cell. Under a cell-based rule, competition and selection are global as might be seen in the presence of antibiotics. In a site-based rule system with selection, a random perimeter site is sampled, a daughter is placed, after which a mother is sampled from the pool of adjacent cells, relative to their *w −* 1. Under a site-based rule, competition and selection are local, with cells competing with their neighbours for resources and space instead of competing with cells of the entire colony. Both methods use single time-steps and terminate when the population reaches a predetermined size, *N*.

**Figure 1.**
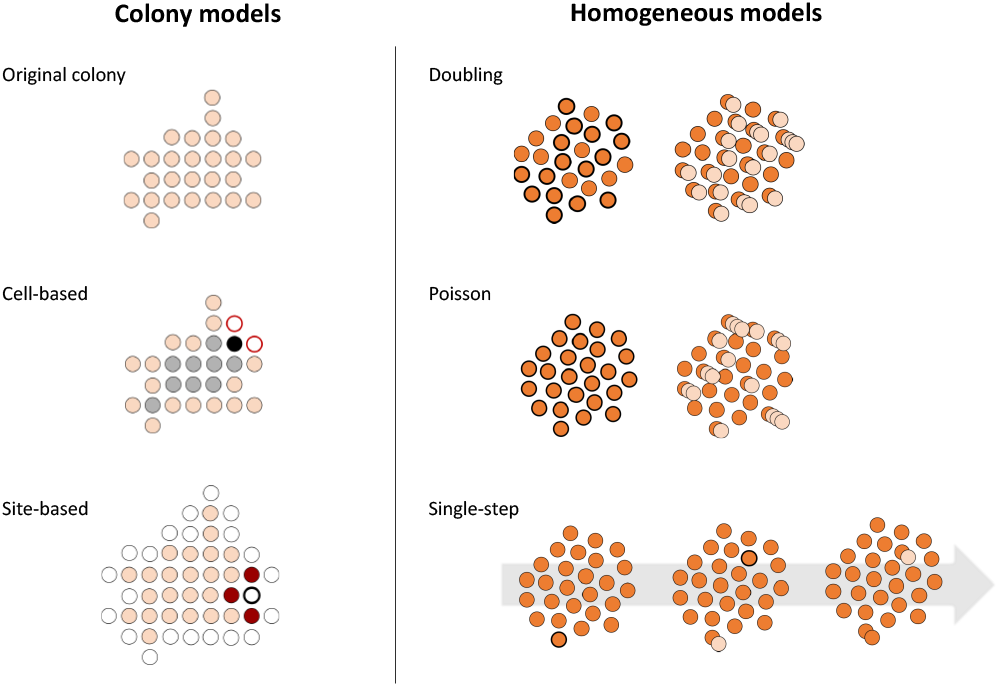
Growth models used in simulations. *Colony models* (left): The colony is made up of individual cells, represented as filled circles. Colony growth occurs when a new cell is placed adjacent to at least one pre-existing cell. This occurs once per time step. In the cell-based model, an edge cell (i.e. a cell that has at least one adjacent free space, pink circles) is chosen from all edge cells with a probability proportional to its fitness. Once a parental cell has been selected (black circle), the daughter cell is placed into a random adjacent free space (marked with thick red borders). In the site-based model, all free spaces that are adjacent to at least one cell (perimeter sites) are identified (empty circles) and one is chosen at random (empty circle with a thick black edge). A parental cell is chosen from all cells that are adjacent to the chosen perimeter site (dark red circles) with a probability proportional to its fitness. *Homogeneous growth models* (right): No spatial structure exists in the cell population. Cells chosen to be parents are always marked with a thick border, daughter cells are marked in a lighter colour. In the Doubling scenario, the population doubles each generation. Parental cells are sampled with replacement from the existing population proportional to their fitness. In the Poisson model, the number of daughters produced by each parent is drawn from a Poisson distribution with the mean equal to the fitness of the parental cell. The Single-Step model allows population growth to occur one cell at a time, as in colony based models. During each time step, one parental cell is chosen from the overall population to produce a daughter, where a cell is sampled proportional to its fitness.

**Figure 2.**
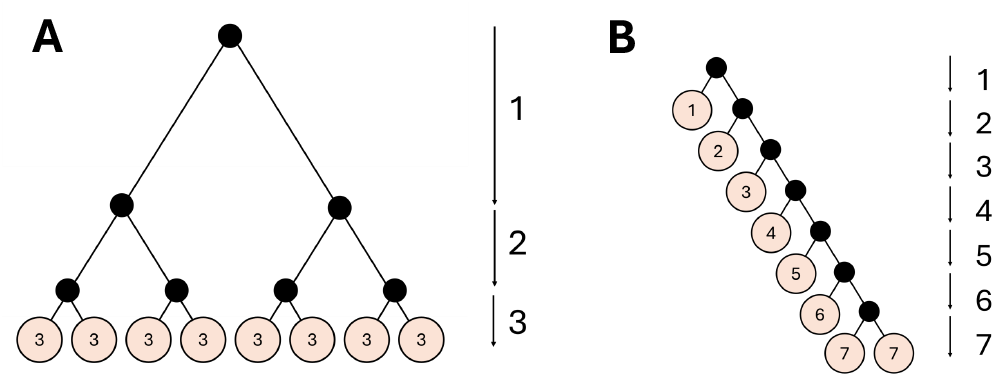
Homogeneous and colony growth can share the same number of doublings and final population size, yet differ in the number of generations. Cell lineage trees are shown for homogeneous growth (A) and an extreme example of colony growth (B). In both scenarios there are seven divisions and hence three doublings (*τ* = 3), yielding a final population of eight cells. However, because cell divisions occur synchronously in A but sequentially in B, the number of generations experienced by the final cells differs between the two models: in A every final cell has undergone three generations, whereas in B the final cells have undergone between one and seven generations, with a mean of 4.375. Small black circles represent fission events, the final cells in the population are shown as light pink circles. The number of generations is shown to the right of each tree and inside each cell.

We also implemented three homogeneous growth models for comparison (Figure 1). The model simulated in Wahl and Agashe (2022) we refer to as the Poisson model and assumes discrete generations, where each parent in a given generation produces, on average, *w−* 1 daughters. The simulation terminates after *log*_2_(*N*) generations, resulting in a stochastic final population size that has expectation *N* if mutations are neutral.

To make sure any differences we see when comparing colony versus homogeneous growth models are not attributable to the stochastic final population size and discrete generations, we implemented two more homogeneous growth models. In the Doubling simulation, the population doubles each generation, with parents for the 2^*t*^ offspring in generation *t* sampled with replacement from the 2^*t−*1^ individuals in generation *t −* 1 with probabilities proportional to *w−* 1. This model uses discrete generations but with a predetermined final population size. Similarly, the Single-Step model also terminates at a predetermined final population size, but growth is simulated using single time-steps rather than discrete generations. Like the Doubling scenario, parents of the single offspring at time *t* are sampled from the *t−* 1 parents proportional to *w −*1. In the Single-Step model there are *N* time steps, as in the colony growth models, rather than *log*_2_(*N*).

Except for the Poisson model, all models terminate once population size has reached N=65,536 cells, corresponding to 16 doublings (*τ* = 16). For the Poisson model, *τ* is the expected number of doublings since the total number of daughter cells per generation is stochastic. Although *τ* is at the lower end of what is typical in MA studies (and substantially smaller than in a typical *E. coli* MA experiment - see Table 1) – downscaled simulations (see below) show that the qualitative behaviour of the model remains consistent above *τ ≈* 12.

#### Mutations and the distribution of fitness effects (DFE)

In our simulations, an initial cell is assigned a fitness of two (*w* = 2). For each cell division, the mother and daughter cells have the chance of acquiring a new mutation with probability *µ* = 10^*−*3^ (approximately the genomic mutation rate of *E. coli* (Lee *et al*. 2012)). Note that we are assuming a constant mutation rate per cell division, rather than per time, and so most mutations are occurring in the dividing cells on the colony periphery. Following Wahl and Agashe (2022), selection coefficients *s* are drawn from a mean-shifted reflected gamma DFE, and the fitness of the cell is updated to *w*(1 + *s*) where *w* is the fitness of the cell prior to mutation. We set fitness to one in the rare cases when *s* was sufficiently negative that fitness would become less than one and incompatible with an absence of cell death. The DFE in the main simulations had a mean of −0.2, a shape of 10, and a scale of 0.05, as used by Wahl and Agashe (2022), which we refer to as the ‘wide-DFE’. We also ran simulations with a ‘narrow-DFE’, with a mean of −0.02, a shape of 5, and a scale of 0.01, more consistent with previous estimates of the DFE (Kibota and Lynch 1996; Imhof and Schlötterer 2001; Robert et al. 2018; Couce *et al*. 2024).

In a real wild-type MA experiment, the founder cell for each new colony is chosen blindly. Most founder cells therefore carry no new mutations, meaning many growth cycles are needed before a single-mutant cell is picked by chance. In our simulation, the procedure can be made more efficient by stipulating that the founder cell must contain exactly one mutation. Therefore, after simulating one round of colony growth (for *τ* = 16), a founder cell is chosen from the subset of colony cells which acquired one mutation, mimicking the moment in a real MA line when a mutant founder is randomly chosen. Given the simulation conditioning on sampling a single-mutant founder, one simulated growth cycle was sufficient in all replicates. For each scenario, 2,000 MA lines (and hence mutations) were simulated and we refer to the sampled selection coefficients using this procedure as 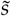 to distinguish them from the selection coefficients that would be observed in the absence of selection (*s*).

#### Assessing magnitude and impact of selection bias

Under the assumption of homogeneous growth, Kibota and Lynch (1996) derived an expression for (one minus) the odds that a mutation with selection coefficient *s* is sampled versus not. Using the same model, but assuming that the mutation rate is negligible compared to the growth rate, Wahl and Agashe (2022) obtained an expression for the odds that a mutation with selection coefficient *s* is sampled compared to a new neutral mutation. By using the odds relative to a new neutral mutation, rather than an unmutated cell, the odds do not depend on the mutation rate (Equation 2 in Wahl and Agashe 2022):

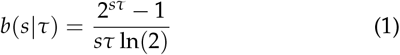

Note that the underlying model is deterministic, meaning no drift is modelled, and so increasing the number of population doublings (*τ*, where the total number of cells N=2^*τ*^) results in stronger selection bias because selection is able to operate over a longer period of time, not because the resulting population is larger. Since *s* and *τ* always appear as a product, changing *τ* simply adjusts the strength of selection.

We assess the accuracy of homogeneous-growth models of selection bias by visually inspecting how well they predict the distribution of *sampled* selection coefficients 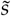 (i.e. those filtered by selection during MA), but also how well they predict the distribution of *true* selection coefficients *s* (i.e. the DFE) after bias correction has been applied. In our simulations described above we effectively draw sampled selection coefficients from their distribution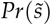, which we will refer to as the distribution of sampled fitness effects (DSFE). The DSFE is proportional to *Pr*(*s θ*)*b*(*s*) where *Pr*(*s* | *θ*) is the true DFE with parameters *θ* (in our case the mean, shape and scale of the reflected mean-shifted gamma) and *b*(*s*) is the selection bias, which in general has an unknown form. We refer to the DSFE predicted by homogeneous-growth models (Wahl and Agashe 2022) as the DSFE_*τ*_, where *b*(*s* |τ = 16) from Equation 1 is used for *b*(*s*). We refer to the DFE estimated using the bias correction derived under homogeneous-growth as the DFE_τ_, where we obtain estimates of the parameters of the DFE 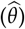 via maximum likelihood given the distribution of sampled selection coefficients and the assumed bias function. Note that our approach for obtaining the DFE_*τ*_ is similar to the maximum likelihood approach used by Joseph and Hall (2004) rather than the non-parametric approach used by Wahl and Agashe (2022). Like Wahl and Agashe (2022) our method can only be applied to single mutant lines, and extending it to multi-mutant lines would require inferring selection coefficients of individual mutations from the aggregate effect of all mutations in a line, something that would be particularly challenging in the presence of epistasis.

The degree to which the DFE_τ_ and DSFE_τ_ were good representations of the DFE and DSFE, respectively, differed across the growth models. In order to quantify the discrepancies, we assumed that the selection bias has the same form as assumed by Equation 1 but differed in strength. The adjusted strength most consistent with the DSFE given the true DFE was obtained by estimating the *effective* number of doublings (*τ*_*e*_) using maximum likelihood, rather than assuming it to be 16. Note this can only be done because we use simulated data and have access to both the DFE and the DFSE. If *τ*_*e*_ is smaller than 16, the degree of bias predicted by homogeneous-growth models is too strong (i.e. selection is *weaker* than predicted), and if *τ*_*e*_ is larger than 16, the degree of bias is too weak (i.e. selection is *stronger* than predicted). We refer to the DSFE predicted using *b*(*s* |*τ* = *τ*_*e*_), rather than *b*(*s* |*τ* = 16), as the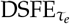, and tested whether this *adjusted* distribution is significantly better than the assumed distribution (DSFE_*τ*_) using likelihood-ratio tests. See Eden Model SI for details of the simulation and the maximum-likelihood procedures introduced above.

Related work on cell-based colony growth has identified that the rate of drift per generation is higher than under homogeneous growth (Hallatschek et al. 2007), but the number of generations over which selection operates accelerates with *τ* rather than being equal to *τ* (Gralka et al. 2016). Consequently, we might expect selection to be less efficient under colony growth when *τ* is small but as it increases the pattern may be reversed. To address this we performed the cell-based and site-based colony simulations described above with *τ* ranging from 8 (N=256) to 17 (N=131,072). Mutations were generated using the wide-DFE and 1,000 collected mutations were used to estimate *τ*_*e*_.

#### Number of generations

In MA studies, the number of generations is taken to be *τ* = *log*_2_(*N*), which constitutes a lower bound on the number of generations as it requires all cells to divide synchronously (Figure 6). In the absence of cell-death, a genealogy can be constructed for all cells, with internal nodes representing binary fission events. The average number of generations is equal to the number of internal nodes averaged over each cell, which is Sackin’s index (Shao and Sokal 1990) normalised by the number of cells. Under asynchronous growth where the next cell to divide is chosen at random (i.e. the Single-Step model under neutrality), the expectation of Sackin’s index is a harmonic series (Kirkpatrick and Slatkin 1993) such that the average number of generations is approximately 2*ln*(2)*τ* ™ 1 + *γ*, where *γ* is Euler’s constant. For colony growth, Gralka *et al*. (2016) noted that the number of generations is approximately proportional to the radius of the colony, *r*, yet the total number of cells is approximately proportional to its area, *πr*^2^ (see Fusco *et al*. (2016) also). Since *τ* ∝ *log*_2_(*πr*^2^), the expected number of generations is proportional to *e*^*τln*(2)/2^ and *τ* will be a poor measure of the number of generations, particularly in large colonies. In order to assess whether these relationships hold in our simulations, we also recorded the average number of cell divisions experienced by cells in populations ranging from a *τ* of two (N=4) to 16 (N=65,536) for both colony models and the Single-Step growth model. Each combination of model and *τ* was simulated ten times in the absence of mutations and the average of the average number of generations recorded.

### Meta-analysis of mutation accumulation experiments

Papers cited in Wang and Obbard (2023) and Lynch *et al*. (2023) reporting mutation rate estimates from wild-type strains (i.e. not hypermutators) of unicellular organisms were checked and retained if they had performed colony-based MA and reported a test of whether coding/non-coding and/or non-synonymous/synonymous (NS/S) ratios were different from expectation. The number of non-coding, synonymous coding and non-synonymous coding base-pair substitutions were recorded together with the expected ratios and the number of population doublings between bottlenecks, *τ* (Table 1). Whether codon-usage bias, transition/transversion ratios or GC bias were controlled for when obtaining the expected NS/S ratio was also recorded in addition to the significance of any tests performed. 29 studies provided complete information on the number of non-synonymous and synonymous mutations and their expected ratio. In some cases the values recorded are different from those reported: a) Kucukyildirim *et al*. (2021) report an observed NS/S ratio of 2.12 yet also state 25 out of 63 coding mutations were synonymous - after checking the supplementary information we used an observed NS/S ratio of 38/63=1.5. b) Foster *et al*. (2015) only reported *τ* for *E. coli* MG1655K12 grown in Lysogeny Broth, and we use the same value for Minimal Media. c) The numbers reported for Long *et al*. (2016a) were calculated from the supplementary data and differ slightly from those reported in the main text. d) López-Cortegano *et al*. (2021) reported a *K*_*a*_ /*K*_*s*_ ratio of 0.84 which is the observed/expected ratio. The authors were contacted for the observed ratio and the expected ratio was calculated by dividing the observed ratio by 0.84. e) For Ness *et al*. (2015), we use the number of mutations at zero-fold and four-fold sites, instead of the number of non-synonymous and synonymous mutations respectively, since only these numbers are reported. f) Sun *et al*. (2017) were contacted for *τ*. For discrepancies regarding statistical significance, see the results section on statistical power. For discrepancies in studies reported in Table 1 but not forming part of the meta-analysis, see the caption to Table 1.

To test whether the NS/S ratio deviated from expectation, a binomial generalised linear mixed model (GLMM) with logit link was fitted to the number of non-synonymous and synonymous mutations using the lme4 library (Bates *et al*. 2015) in R (R Core Team 2025). The logit link models *ln*(*E*[*r*_*obs*_]), where *r*_*obs*_ is the observed NS/S ratio, and so the log of the expected ratio (*r*_*exp*_) was fitted as an offset. The intercept of this model is an estimate of the average deviation of the observed ratio from the expected ratio on the log scale. Observation-level random effects were fitted to deal with any overdispersion. We also tested whether the deviation differed between studies that controlled for differences in transition/transversion rate and those that did not, since our multinomial model of *E. coli* mutations suggests a failure to control for a difference may result in an apparent deficit of non-synonymous mutations (see below). To place the deficit of non-synonymous mutations in the context of selection we also fitted a binomial GLMM with logit link, but with the expectation given by *ln*(*r*_*exp*_*b*(*s*| *τ*)) with *s* estimated using maximum likelihood. Under deterministic homogeneous growth, this model gives the *s* most compatible with the observed deficits if all synonymous mutations are neutral and all non-synonymous mutations have the same selection coefficient, *s*.

### *E. coli* mutation accumulation and analysis

#### New mutation accumulation data

MA lines were carried out using a MG1655-K12 *E. coli* isolate, from now on the ‘ancestor’, which was previously obtained from the *E. coli* Genetic Stock Center. The ancestral strain was streaked out onto 110 Lysogeny broth (LB) agar plates. Each plate was divided into two halves, each half carrying one line. Every 24 hours colonies from the two halves were re-streaked, without vortexing, onto new LB agar plates. To reduce biased colony propagation, an arbitrary colony was picked from a pre-determined area of a plate. This was done for 36 days, which corresponds to around 1,008 population doublings, the average time we predicted it would take for a line accumulate one mutation, based on mutation rates and doubling rates from Lee *et al*. (2012). Eight out of the 220 lines did not complete the full mutation accumulation cycle due to human error (six lines), extinction (one line) and possible contamination (one line). Lines were subsequently frozen at −80°C for future fitness measurements.

In addition to the ancestral line, which was sequenced twice, 194 out of the 212 remaining MA lines were selected for sequencing, determined by the quality of their DNA extractions. DNA extractions were performed with Qiagen DNeasy Blood and Tissue Extraction Kit (96-wells method) and stored at *−* 20°C. DNA was sequenced at the Centre for Genomic Research, University of Liverpool, with NovaSeq SP (2x150), generating an average coverage of 97.1X per line. A full description of the bioinformatic pipeline used for mutation calling can be found in (Grosse-Sommer et al. 2026) and here we summarise the main points. Read integrity and quality was assessed with FastQC and Multi-FastQC, where all criteria were passed. Reads were subsequently aligned to the MG1655-K12 *E. coli* reference genome (RefSeq accession NC_000913.3, assembly GCF_000005845.2 (ASM584v2)) using the bwa-mem package (v0.7.19) (Li 2013) with default parameters. PCR duplicates, secondary reads and non-mapped reads were excluded using samtools (v1.3.1) (Danecek et al. 2021). Base-pair substitutions (bps) and short insertion/deletions (INDELs) were called with bcftools (v1.23.1) (Danecek et al. 2021). Default parameters were used except extended BAQ was used to reduce false-positive INDEL calls and reads with mapping quality *<* 30 and bases with base-quality *<* 20 were excluded. Variants at sites with a Phred-scaled score (QUAL) *<* 20 were excluded, in addition to two ancestral mutations that were present in all lines. 255 possible mutations were identified, of which 61 had fewer than 30 reads, or *≤* 95% of reads, supporting the alternate allele. These ‘provisional’ mutations were visualised in the Integrative Genomics Viewer (IGV) (Robinson et al. 2011). 41 of these were considered real mutations, with the remainder mainly falling in rRNA genes, insertion sequences or the *icd* gene of a single line. An additional *icd* mutation in the same line, which did not fall on the provisional list, was also excluded, as was one duplicate mutation that was likely due to contamination (the mutation was shared by two lines propagated on the same plate). One line had 22 mutations passing the initial filter, one of which was in the mismatch-repair gene, *mutS*. These mutations (including the 5 on the provisional list which probably occured after mutation accumulation during the growth stage prior to sequencing) were removed from further downstream analyses. Large structural variants (LSVs) were called using Parliament2 (Zarate *et al*. 2021), where LSVs were called by compiling the results of the programs Lumpy, Breakdancer and CNVNATOR. Three large deletions were detected and manually checked in IGV. One large deletion was caused by the excision of a cryptic prophage integrated into *icd*. Mutations were annotated using the Bioconductor (Huber *et al*. 2015) libraries VariantAnnotation (v1.56) (Obenchain *et al*. 2014), GenomicFeatures (v1.62) (Lawrence *et al*. 2013) and rtracklayer (v1.70.1) (Lawrence *et al*. 2009) for R (v4.5.1) (R Core Team 2025). GTP-5.5 was used to troubleshoot the bioinformatic pipeline.

A total of 203 mutations were detected of which 176 (86.7% of all mutations) were bps, 24 (11.8%) were INDELs and 3 (1.5%) were LSVs. Of all bps, 133 (75.6%) were coding, and of these, 101 (75.9%) were non-synonymous, including two premature stop mutations and one stop to non-stop mutation. Given Lee *et al*.’s 2012 generation estimate of 28 generations per MA transfer, the per genome per generation mutation rate was found to be (after excluding the hypermutator) 1.049[0.904 – 1.208] *×* 10^*−*3^. For bps the mutation rate was 0.909[0.774 – 1.059] *×* 10^*−*3^. The mutation counts across non-excluded lines had a variance/mean ratio of 1.111, close to the Poisson expectation of one.

#### Additional mutation accumulation data

In order to increase the power to detect mutational signatures of selection, we combined our MA data with comparable data from previous *E. coli* MA studies. Studies were selected based on the following criteria: mutations must have occurred in a wild-type (WT) MG1655-K12 strain (i.e. not a hypermutator), not exposed to potential mutagens and grown on LB agar using colony propagation as the MA method. Hypermutator strains were excluded for two reasons. First, hypermutators may experience a DFE distinct from that of wild type (Sane *et al*. 2023), which could amplify or dampen selection bias. Second, their mutational spectrum can differ from those of WT strains (Lee *et al*. 2012), making reliable spectrum correction difficult. Similarly, lines grown on media other than LB were excluded since different media types can influence the mutation spectrum (Foster *et al*. 2015). In addition we also excluded lines that had gone through experimental evolution prior to the MA process. Five studies met all criteria: Foster *et al*. (2015); Sane *et al*. (2018); Long *et al*. (2016b); Tincher *et al*. (2017) and Wei *et al*. (2022) (ancestral clones only). Data were sourced from the supplements of these papers, except for Sane *et al*. (2018) who kindly provided us with their data upon request. Note that the subsequent analysis was restricted to bps only. Foster *et al*.’s 2015 mutations were originally aligned to an earlier version of the MG1655-K12 genome (NC_000913.2). We used minimap2 (v2.31) (Li 2018) to align the two assemblies and then a paftools.js liftover to convert positions. In each study there were duplicate mutations, likely representing contamination (Foster *et al*. (2015): one duplicate pair, Sane *et al*. (2018): 61 duplicate pairs, Wei *et al*. (2022): seven duplicate pairs) and in each case only one of the pair was retained. After collating mutations from all studies, three sites had mutated twice across studies, and at one of those sites the alternate allele was different. Since our model does not allow multiple mutations at a site (see below), we randomly chose a mutation for inclusion (position 4171941, kept C to G mutation, removed C to A mutation). The final number of bps per study were 243 (Foster *et al*. 2015) (61 lines for 4,230 generations), 79 (Long et al. 2016b) (48 lines for 1,682 generations), 81 (Tincher et al. 2017) (32 lines for 2,427 generations) 176 (Sane et al. 2018) (38 lines lines for 8,400 generations), 114 (Wei et al. 2022) (100 lines for 1,680 generations) and 176 (this study - 192 lines for 1,008 generations), resulting in a total of 869 mutations, of which 639 were coding.

#### Annotation

We determined whether a site was in a coding or non-coding region using the RefSeq annotation file GCF_000005845.2. Sites labelled as ‘CDS’, but excluding ‘pseudogenes’, were classified as coding resulting in 84.92% of the genome being considered coding. The ratio of coding to non-coding sites (5.63) was comparable to that in earlier annotations which did not exclude pseudogenes (5.71 - Foster *et al*. (2015)). For our mutational model (see below) we also labelled the reference base as being on the leading strand if it was located clock-wise between the oriC location (designated as the nucleotide 3,923,883) and the termination point where replication direction changes, which we took as the genomic midpoint from oriC (position 1,603,057). We also recorded the identity of the bases immediately 5’ and 3’ of the focal site on the leading strand. Sites appearing in the Highly Expressed Genes Database (HEG-DB) (Puigbò et al. 2008) were designated as highly expressed. Our mutational model not only allows site-level predictors, but also predictors associated with the type of mutation that may occur. Consequently, for each site, we determined whether the four possible bases following a mutation would involve a transition or a transversion, and if in coding sequence, would cause a synonymous or non-synonymous change. The expected non-synonymous to synonymous ratio given (only) the observed codon usage was calculated to be 3.24, almost identical to the odds from a previous annotation (3.25 - Foster *et al*. (2015)).

#### Multinomial model of mutation

In order to model mutational patterns, we aggregated the mutational data into a single leading-strand aggregate MA sequence where each site has the original reference base if no mutation was observed in any MA line, and the mutated base if a mutation was observed in one of the lines. We model the four possible bases at each site using a multinomial logit model with a single trial, implemented in the R package mlogit (v1.1-3) (Croissant 2020) for R (R Core Team 2025). Note this is an approximation to the ideal model where each site would have as many trials as there are MA lines. However, since the probability of more than one mutation at a site is low the approximation should be very accurate.

The model calculates the (log) odds of observing a mutation in a specific context while taking into account the mutational spectrum, and also codon usage bias in the case of non-synonymous or synonymous mutations. This enables the direct effects of different genomic factors altering mutation probability to be estimated. The probability of observing base *M* at site *i* in the leading-strand aggregate MA sequence, conditional on the original base being *O*, is proportional to

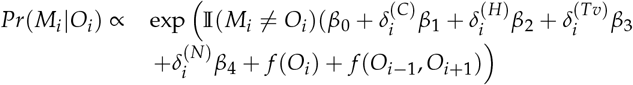

*β*_0_ represents the log-odds ratio that a site in the reference category will be mutated rather than not. In this case, the reference category is the middle base of the original non-coding sequence AAT which mutates into ATT (i.e. a non-coding transition). The reference log-odds ratio is incremented by *β*_1_ for coding sequence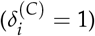, and further incremented by *β*_2_ if the gene is highly expressed 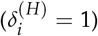. The reference log-odds ratio is also incremented by *β*_3_ for mutations that are transversions 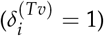 and by *β*_4_ if the mutation causes a non-synonymous change in a coding sequence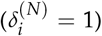. Note that these odds are after controlling for mutational opportunity (i.e. by chance mutations are more likely to be non-synonymous transversions).

The function *f* (*O*_*i*_) allows the mutation rate at site *i* to depend on the focal base in the original sequence:

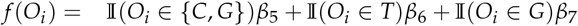

where *β*_5_ is the effect of having a leading or lagging C, *β*_6_ is the effect of having a leading T and *β*_7_ is the effect of having a leading G. The function *f* (*O*_*i −* 1_, *O*_*i*+1_) allows the mutation rate at site *i* to depend on the neighbouring bases in the original sequence:

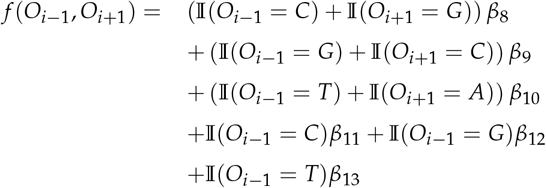

where *β*_8_, *β*_9_ and *β*_10_ are the effects of having a C, G or T, respectively, immediately 5’ of the focal site on either the leading or lagging strand. *β*_11_, *β*_12_ and *β*_13_ are the effects of having a C, G or T, respectively, immediately 5’ of the focal site on the leading strand only. If mutational patterns are independent of strand identity, then *β*_6_ = *β*_7_ = *β*_11_ = *β*_12_ = *β*_13_ = 0. A more in depth explanation of the model can be found in Multinomial Model SI. While our model only considers the effect of the direct neighbours on the focal site, long-range sequence context effects have been shown to affect mutation rate (Jago *et al*. 2026) and could be modelled.

## Results

### Homogeneous versus colony-based models of growth

To compare selection bias under colony growth and homogeneous growth, we simulated five growth models under both a ‘wide-DFE’ dominated by large-effect mutations and a ‘narrow-DFE’ dominated by weakly deleterious mutations. Under the wide-DFE, for all homogeneous growth models, the estimate of the effective number of population doublings (*τ*_*e*_) was significantly less than 16, implying significantly less selection than predicted by the deterministic homogeneous growth model (Wahl and Agashe 2022) (see Table 2 for all simulation results). *τ*_*e*_ was largest for the Single-Step (*τ*_*e*_=14.109*±*0.743, P=0.012), followed by the Poisson (*τ*_*e*_=9.696 *±* 0.670, P*<*0.001) and the Doubling model (*τ*_*e*_=9.550*±*0.661, P*<*0.001). Despite selection bias being significantly weaker than predicted, graphical comparisons of the predicted distribution of sampled fitness effects under *τ* = 16 (DSFE_*τ*_, red line) and that predicted using the estimate of *τ*_*e*_ (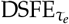blue line) showed very close correspondence, particularly for the Single-Step model (Figure 3). Similarly, the predicted distribution of fitness effects (DFE_*τ*_, dotted line) showed good correspondence with the true DFE (black line). In contrast, the colony-based simulations, showed large deviations between *τ*_*e*_ and *τ*, and the predicted distributions (both DSFE_*τ*_ and DFE_*τ*_) showed poor correspondence with their true respective distributions. Under the cell-based colony model, where competition is global, the observed strength of selection was considerably larger than predicted under homogeneous growth (*τ*_*e*_=36.877 *±* 0.630, P*<*0.001) and the predicted DSFE_*τ*_ failed to account for the removal of strongly deleterious mutations and the enrichment for beneficial mutations (Figure 3). Consequently, the DFE_*τ*_ was biased towards more positive selection coefficients. In contrast, *τ*_*e*_ was much smaller than *τ* in the site-based colony model where competition is local (*τ*_*e*_=5.328 *±* 0.544, *P<*0.001), indicating that the strength of selection is considerably less than predicted by the deterministic homogeneous growth models and their stochastic counterparts. Consequently, the DSFE_*τ*_ predicts greater removal of deleterious mutations and enrichment of beneficial mutations than is observed and the DFE_*τ*_ is biased towards more deleterious selection coefficients.

**Table 2.** Estimates of the effective strength of selection as estimated by *τ*_*e*_, for colony growth (cell-based and site-based) and homogeneous growth (Poisson, Doubling and Single-Step) models. The DFE was a mean-shifted reflected gamma with either a mean of −0.2, a shape of 10 and a scale of 0.05 (wide-DFE) or a mean of −0.02, a shape of 5 and a scale of 0.01 (narrow-DFE). In all simulations, the number of population doublings was *τ* = 16 and the null-hypothesis being tested is that *τ*_*e*_ = *τ* = 16.

| Model | DFE | $\tau_e$ | Standard Error | p-value |
| --- | --- | --- | --- | --- |
| Cell-based | wide | 36.877 | 0.630 | $< 10^{-16}$ |
| Site-based | wide | 5.328 | 0.544 | $< 10^{-16}$ |
| Poisson | wide | 9.696 | 0.670 | $< 10^{-16}$ |
| Doubling | wide | 9.550 | 0.661 | $< 10^{-16}$ |
| Single-Step | wide | 14.109 | 0.743 | 0.012 |
| Cell-based | narrow | 74.903 | 4.551 | $< 10^{-16}$ |
| Site-based | narrow | 6.523 | 3.071 | 0.003 |
| Poisson | narrow | 8.782 | 3.175 | 0.027 |
| Doubling | narrow | 14.179 | 3.254 | 0.578 |
| Single-Step | narrow | 11.021 | 3.201 | 0.127 |

**Figure 3.**
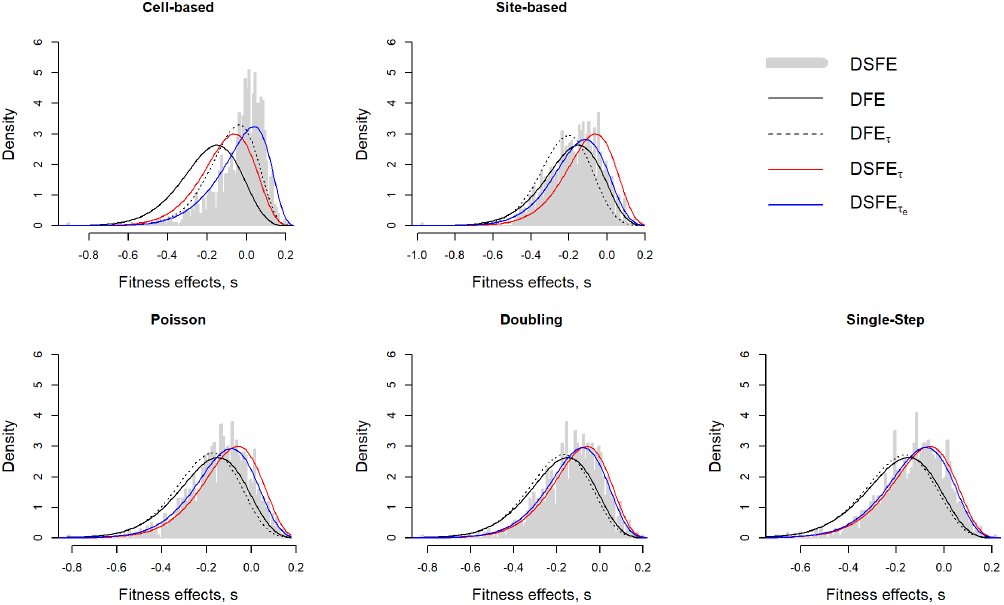
Histograms of sampled fitness effects (DSFE) from MA lines simulated under colony or homogeneous growth with *τ* = 16 population doublings with a wide-DFE. The true distribution of fitness effects (DFE) is plotted as a solid black line and is a mean-shifted reflected gamma distribution with a mean of −0.2, shape of 10 and scale of 0.05 (wide-DFE). The estimates of the DFE from the simulated data after applying the homogeneous-growth correction (DFE_*τ*_) are plotted as dashed black lines. The predicted distributions of sampled fitness effects given the homogeneous model of selection bias are plotted as red lines when *τ* is set to its true value of 16 (DSFE_*τ*_) or as blue lines when an effective *τ* is estimated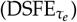.

Since the derivative of *b*(*s* | *τ*) with respect to *τ* decreases as *s* becomes small, the power to distinguish between different values of *τ*_*e*_ is reduced when using a narrower DFE, resulting in larger standard errors. Nevertheless, the same general pattern observed for the wide-DFE held for the narrow-DFE, although the average amount of selection bias experienced by a mutation was considerably weaker since selection coefficients are smaller. Indeed, under the homogeneous growth models, graphical comparisons of the true DFE with the DSFE, DSFE_*τ*_ and 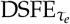 revealed little discrepancy (Figure 4). However, for cell-based colony growth the DSFE diverged much more from the DFE and both the DSFE_*τ*_ and DFE_*τ*_ were poor representations of their true counterparts. As with the wide-DFE, the actual amount of selection bias was stronger than predicted by deterministic homogeneous growth models (*τ*_*e*_=74.903 *±* 4.551,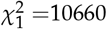, *P <*0.001). For the site-based colony model, the amount of selection bias was once again weaker than predicted by homogeneous growth models (*τ*_*e*_=6.523*±*3.071, P=0.003). However, given that selection bias was already weak under homogeneous growth for the narrow DFE, a further reduction in the amount of selection bias in the site-based model meant there was little discrepancy between the DSFE_*τ*_ and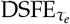.

**Figure 4.**
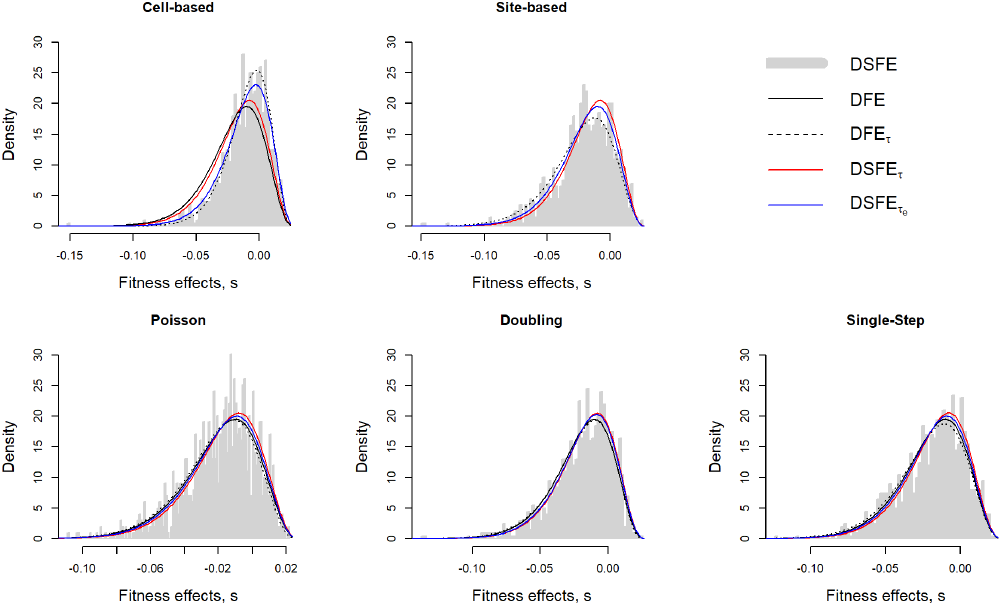
Histograms of sampled fitness effects (DSFE) from MA lines simulated under colony or homogeneous growth with *τ* = 16 population doublings with a narrow-DFE. The true distribution of fitness effects (DFE) is plotted as a solid black line and is a mean-shifted reflected gamma distribution with a mean of −0.02, shape of 5 and scale of 0.01 (narrow-DFE). The estimates of the DFE from the simulated data after applying the homogeneous-growth correction (DFE_*τ*_) are plotted as dashed black lines. The predicted distributions of sampled fitness effects given the homogeneous model of selection bias are plotted as red lines when *τ* is set to its true value of 16 (DSFE_*τ*_) or as blue lines when an effective *τ* is estimated 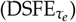.

Smaller scale simulations using the wide-DFE over a range of *τ* showed that under cell-based colony growth the increase in selection bias relative to that expected under homogeneous growth increases with *τ* (i.e *τ*_*e*_ /*τ* becomes larger with increasing *τ*). When *τ* is less than 12 the selection bias under cell-based colony growth is less than that seen under homogeneous growth, but for larger *τ*, this is reversed. Selection bias under site-based colony growth is consistently less than that expected under homogeneous growth at all *τ* (Figure 5).

**Figure 5.**
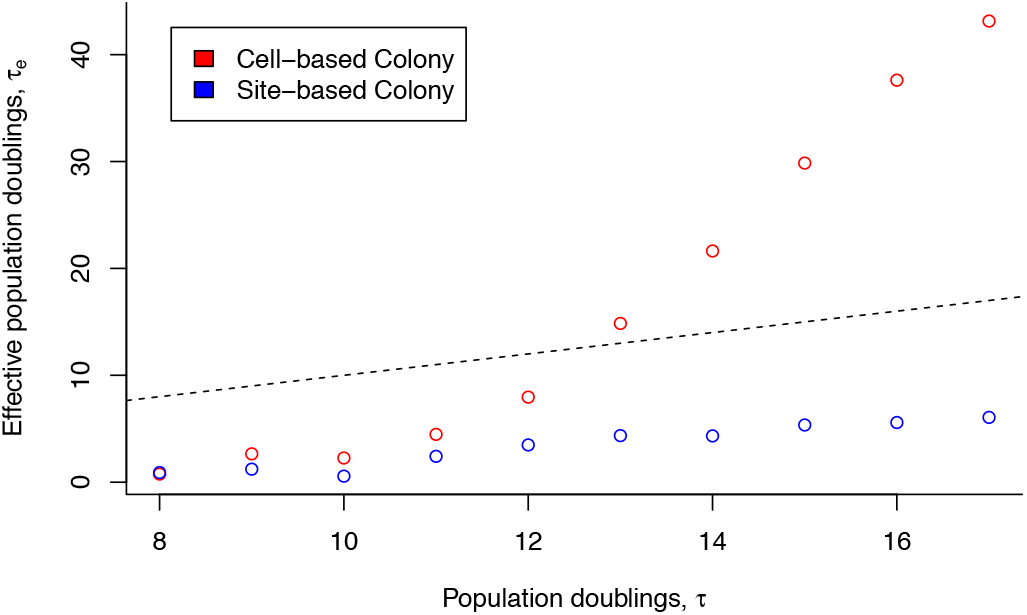
Effective versus actual *τ* under colony growth. Each point is the average of 10 simulations either under cell-based colony growth (red), or site-based colony growth (blue) using the wide-DFE. The dashed black line is the 1:1 line, where the amount of selection bias is equal to that under a deterministic homogeneous growth model. Values above this indicate selection bias is stronger than predicted.

#### Number of generation

When *τ* = 16, the Single-Step homogeneous-growth simulations produced an average of 21.3 generations, whereas the number of generations under colony growth was considerably higher (cell-based: 179, site-based: 184). Across a range of *τ*, the number of generations in the Single-Step homogeneous-growth simulations closely matched the theoretical expectation based on Sackin’s Index (Kirkpatrick and Slatkin 1993) where the number of generations is approximately 2*ln*(2)=1.39 times larger than *τ*. For the colony based models, the average number of generations grew slightly slower with *τ* than predicted assuming circular colonies (Gralka *et al*. 2016), although there was still a rapid exponential increase in the number of generations (Figure 6).

**Figure 6.**
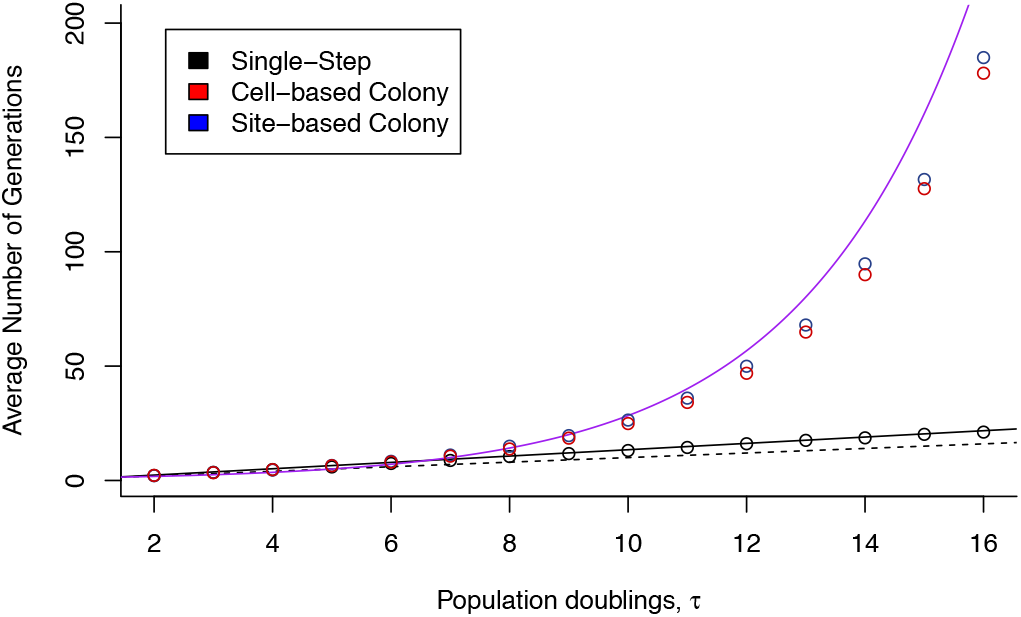
Average number of generations as a function of the number of population doublings, *τ*. Each point is the average of 10 simulations either under Single-Step homogeneous growth (black), cell-based colony growth (red), or site-based colony growth (blue). The dashed black line is the 1:1 line, indicating that *τ* is equal to the average number of generations. The solid black line is the prediction using results from Sackin’s Index (Kirkpatrick and Slatkin 1993) which should apply to Single-Step homogeneous growth. The solid purple line is the prediction based on the assumption that a colony is circular, the number of generations is proportional to the radius of the colony and the final population size is proportional to its area (Gralka et al. 2016).

### Meta-analysis of mutation accumulation experiments

The intercept of the standard meta-analysis was less than zero (−0.080 *±* 0.026, P=0.002) indicating a deficit of non-synonymous mutations compared to expectation: exponentiating the intercept gives 0.923 (i.e. a 7.7% deficit). Studies that controlled for differences in transition/transversion rate reported larger deficits, with a difference in log-odds of −0.092 *±* 0.045, P=0.043. A meta-analysis employing the deterministic homogeneous growth model to predict the deficit, estimated *s* to be −0.011*±*0.003 un-der the assumptions that all synonymous mutations are neutral and all non-synonymous mutations have the same selection coefficient (Figure 7).

**Figure 7.**
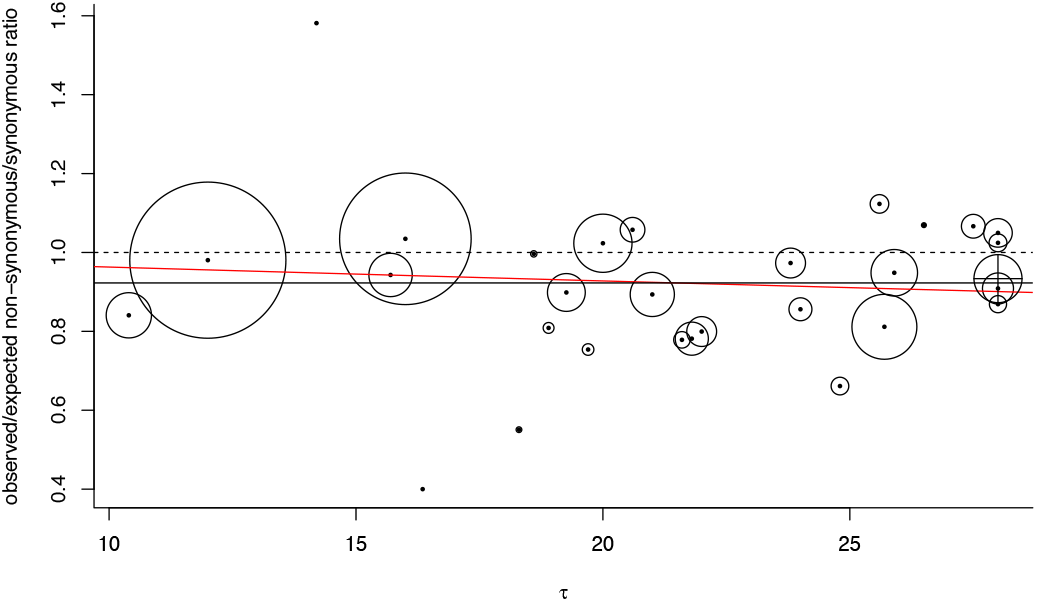
The observed non-synonymous/synonymous ratio divided by its expected value in the absence of selection as a function of number of population doublings, *τ*. Points are scaled by sample size. The dashed line indicates that observed and expected values are equal. The solid black line is the predicted bias from the standard meta-analysis. The solid red line is the predicted bias from a meta-analysis parameterised in terms the selection bias expected from a deterministic homogeneous growth model with the selection coefficient of non-synonymous mutations estimated to be −0.011. The point with the cross is the estimate made from our multinomial logit model of mutations in *E. coli*.

Despite strong evidence for a deficit of non-synonymous mutations, no individual study has reported a significant deficit (Table 1). While the power to detect a deficit equal to the observed meta-analytic deficit of 7.7% is low on average (9.7% over the 30 studies for which it could be calculated, using a two-tailed test) we should still expect approximately 3.9 of the 40 published tests to have been significant. Indeed, when performing simple two-tailed binomial tests on each study individually, 4 of the 29 studies had significant deficits. While it is possible that the methods used in the original tests also took into account uncertainty in the null expectation (as in our multinomial model below), resulting in reduced power but a better calibrated significance threshold, most discrepancies seemed to be reporting errors. Takemoto *et al*. (2018) reported 131 non-synonymous and 64 synonymous mutations and reported a *χ*^2^ value of 3.4 and a p-value of 0.07 against the null hypothesis that the ratio is 3.1. In fact, *χ*^2^ =7.5 and the p-value is 0.006. Kucukyildirim *et al*. (2020) reported an observed non-synonymous ratio of 2.12 but the correct value is 1.52 which gives a p-value of 0.022 rather than 0.29. Kucukyildirim *et al*. (2016) used a significance threshold of 0.01 (rather than the typical 0.05) to declare the deficit non-significant: a two-tailed binomial test gives a p-value of 0.011. Finally, Nguyen *et al*. (2020) reported a p-value of 0.48 for *Hanseniaspora osmophila* yet a two-tailed binomial test gives a p-value of 0.029 - the reason for the discrepancy is not obvious.

### *E. coli* mutation accumulation and analysis

The multinomial logit model is parametrised in terms of log-odds ratios, and we report the estimates, standard errors and p-values for all model coefficients on this scale in Table 3. However, for ease of interpretation, we report odds ratios below. We also give the p-value against equal odds, but the standard errors are not available on this scale. Note that the odds ratios are referring to the odds of a mutation at a synonymous site relative to a non-coding site, holding all other aspects of the mutational spectrum constant.

**Table 3.** Estimates, standard errors and p-values from a multinomial logit model of mutational signatures in *E. coli*.

| Coefficient | Effect | Estimate | SE | p-value |
| --- | --- | --- | --- | --- |
| $\beta_0$ | Intercept | -9.033 | 0.130 | $< 10^{-16}$ |
| $\beta_1$ | coding/non-coding | -0.716 | 0.102 | $0.245 \times 10^{-11}$ |
| $\beta_2$ | highly expressed gene/non-highly expressed gene | 0.061 | 0.153 | 0.692 |
| $\beta_3$ | transversion/transition | -0.845 | 0.069 | $< 10^{-16}$ |
| $\beta_4$ | non-synonymous/synonymous | -0.069 | 0.091 | 0.448 |
| $\beta_5$ | original sequence CG/original sequence AT | 0.223 | 0.091 | 0.015 |
| $\beta_6$ | original sequence leading T/original sequence leading A | -0.113 | 0.099 | 0.254 |
| $\beta_7$ | original sequence leading G/original sequence leading C | -0.573 | 0.098 | $0.443 \times 10^{-8}$ |
| $\beta_8$ | 5'C leading or lagging strand/5'A leading or lagging strand | 0.073 | 0.103 | 0.476 |
| $\beta_9$ | 5'G leading or lagging strand/5'A leading or lagging strand | 0.560 | 0.095 | $0.326 \times 10^{-8}$ |
| $\beta_{10}$ | 5'T leading or lagging strand/5'A leading or lagging strand | 0.015 | 0.104 | 0.888 |
| $\beta_{11}$ | 5'C leading strand/5'C leading or lagging strand | 0.163 | 0.151 | 0.280 |
| $\beta_{12}$ | 5'G leading strand/5'G leading or lagging strand | 0.071 | 0.132 | 0.593 |
| $\beta_{13}$ | 5'T leading strand/5'T leading or lagging strand | 0.167 | 0.149 | 0.265 |

The odds of a mutation given a site is synonymous versus non-coding is much less than one (*exp*(*β*_1_)=0.489, P*<*0.001) indicating the mutation rate in synonymous coding regions is less than half that in non-coding regions. Without controlling for the mutational spectrum, the odds ratio is greater (0.569, P*<*0.001) and a smaller deficit of mutations in coding regions would be inferred. The odds a mutation is non-synonymous versus synonymous, after controlling for the mutational spectrum, is less than one (*exp*(*β*_4_)=0.934, P=0.448) indicating the chance of observing a non-synonymous mutation is lower than for synonymous mutations, although the effect was not significantly different than equal odds. Without controlling for the mutational spectrum, the odds ratio drops considerably (0.826,

P=0.031) indicating that only controlling for codon usage bias would result in a larger deficit of non-synonymous mutations being inferred. This seems to be largely driven by differences in transition/transversion rates, and once this is controlled for, the deficit is considerably smaller (0.989, P=0.906).

Table 3 details the effects of the mutational spectrum but here we highlight the major patterns, all of which are consistent with results reported in Foster *et al*. (2015). C:G sites have higher mutation rate than A:T sites (*exp*(*β*_5_)=1.249, P=0.015) but this was reduced by a factor of *exp*(*β*_7_)=0.564 (P*<*0.001) if the G was on the leading strand. Having a T on the leading strand reduced the mutation rate at A:T sites, but not significantly so: *exp*(*β*_6_)=0.893, P=0.254. Neighbouring bases had little effect on mutation at a focal site except when there was a 5’ G on the leading or lagging strand, increasing mutation rate by a factor of *exp*(*β*_9_) = 1.751 (P*<*0.001). Having the 5’ G located on the leading strand further incremented the odds by a small, and non-significant, factor: *exp*(*β*_12_) = 1.073 (P=0.593). Sites in highly expressed genes had higher mutation rates by a factor *exp*(*β*_2_) = 1.062, although this was not significant (P=0.692). Finally, the transversion rate was considerably less than the transition rate (*exp*(*β*_3_)=0.430, P*<*0.001).

## Discussion

Deterministic homogeneous-growth models of microbial MA experiments suggest that selection bias should be weak when the underlying DFE is dominated by weakly deleterious mutations (Kibota and Lynch 1996), which in many cases will be more realistic than assuming a DFE dominated by strong (beneficial) mutations (Wahl and Agashe 2022). Using stochastic simulations of colony growth, we show that selection bias can be considerably stronger if the scale of competition is global, or weaker if it is local. Using a meta-analysis of colony-based MA studies, we show a clear deficit of non-synonymous mutations, although several of the included studies did not fully control for the mutational spectrum, which may bias inferences. Using 869 wild-type *E. coli* MG1655-K12 mutations, we apply a new multinomial model that fully accounts for the mutational spectrum, and also show a comparable deficit of non-synonymous mutations, albeit non-significant. Given corroborating evidence from experiments designed to test whether selection dynamics are different in colony and homogeneous growth (see below), we tentatively suggest that selection bias is stronger than standard homogeneous-growth models predict.

The amount of selection bias observed in our stochastic homogeneous-growth simulations was slightly less than deterministic theory predicts, presumably because of a finite *N*_*e*_ and the action of drift (Kibota and Lynch 1996). Under colony growth, reproduction is restricted to cells at the periphery of the colony resulting in a dramatic drop in *N*_*e*_ (Hallatschek et al. 2007). Intuitively, we may therefore assume that selection bias should be further reduced under colony growth. However, Gralka *et al*. (2016) noted that to reach the same final population size more generations need to elapse under colony growth than predicted by the number of doublings *τ* = *log*_2_(*N*). This gives selection more time to alter mutation frequencies, which likely explains the increase in selection bias in the cell-based model. While this same effect is manifest in the site-based model, competition acts on a local scale causing cells with the same mutation to compete with each other. The increase in kin-competition, or equivalently the reduction in *N*_*e*_, increases the per generation amount of drift which must outweigh the effect the increased number of generations. Consequently, understanding the scale at which competition happens within colonies is paramount for understanding the amount of selection bias that is likely to occur.

Four empirical studies have compared selection dynamics under colony growth with those from homogeneous growth: Gralka *et al*. (2016) reported increased selection efficiency under colony growth in *S. cerevisiae* whereas Lavrentovich *et al*. (2016) showed reduced selection efficiency. In *E. coli*, Bosshard *et al*. (2017) showed reduced selection efficiency under colony growth, whereas Baehr *et al*. (2025) found no detectable difference. Superficially, these empirical results seem to contradict each other, with the results from Gralka *et al*. (2016) suggesting a cell-based model of colony growth and the results of Lavrentovich *et al*. (2016) and Bosshard *et al*. (2017) suggesting a site-based model. However, these contradictions partly disappear once it is understood that they are relevant to different aspects of selection. Lavrentovich *et al*. (2016) studied a system with unidirectional conversion from wild-type to a deleterious mutation and found the frequency of the deleterious mutation at mutation-selection equilibrium was higher under colony growth. However, in MA, the low mutation rate and the short time-scale between bottle-necks (often 24 hours) will mean frequencies will be far from their equilibrium (as pointed out by Gordo and Dionisio (2005) when criticising previous theory that had assumed mutation-selection equilibrium). During the pre-equilibrium phase, the change in allele frequencies will be more rapid in colony growth than homogeneous growth, as selection has more generations to act on any arising mutations. Hence, by focusing on equilibrium conditions, the results of Lavrentovich *et al*. (2016) are not directly relevant to MA studies. Similarly, although Bosshard *et al*. (2017) showed that fitness declines due to spontaneous mutations were stronger under colony growth than under homogeneous growth, the total number of generations under the two scenarios was experimentally matched. Consequently, the larger per generation rate of drift under colony growth will be present without the compensating effect of increasing the number of generations over which selection can act. Nevertheless, while these arguments strongly favour the experiment of Gralka *et al*. (2016) as a model of MA, three important differences with MA remain. First, selective pressures were applied by the addition of antibiotics to the growth media. Consequently, the *S. cerevisiae* strains had a slower growth rate, with roughly 12 doublings over 120 hours compared to typical *S. cerevisiae* MA experiments, where colonies go through 16-24 doublings over 24-72 hours (Zhu et al. 2014; Lynch et al. 2008; Liu and Zhang 2019). Reducing the speed at which colonies grow has been shown to reduce the amount of drift in *E. coli* experiments (Gralka *et al*. 2016), which would increase the amount of selection bias than is typically expected in a conventional microbial MA experiment. Second, selection occurred on standing genetic variants present at an initial frequency of 0.02, rather than the spontaneous mutations that arise in either MA or the experiments of Lavrentovich *et al*. (2016) and Bosshard *et al*. (2017). Although new beneficial mutations are initially rare and subject to high rates of stochastic loss in small populations, mutations that escape initial extinction go to fixation faster in populations with a smaller effective size (Charlesworth 2020). Consequently, the soft-sweep set-up in Gralka *et al*. (2016), where the initial frequency of mutants may be high enough that at least one is destined to fix, may be a poor indication of what happens in the hard-sweep context of MA lines and our simulations. However, simulations suggest that the increased selection efficiency observed in Gralka *et al*. (2016) is not specific to soft-sweep scenarios (see Eden Model SI and also SI of Gralka *et al*. (2016)). Third, the wild-type and mutant used in Gralka *et al*. (2016) were strains resistant and susceptible to cycloheximide, an antibiotic that inhibits protein synthesis by binding to ribosomes (Hardesty *et al*. 1973). It is possible that the fitness effects of the antibiotic are largely independent of the local competitive environment and so the selection dynamics are more similar to what would be observed under a cell-based model compared to the selection dynamics of new spontaneous mutations. Indeed, Baehr *et al*. (2025) compared liquid-based and colony-based MA procedures in *E. coli* and found no significant differences in the mutation rate, fitness decline or ratio of non-synonymous to synonymous mutations. However, point estimates were consistent with increased selection under colony growth, and the extremely small fitness effects of the assayed mutations would reduce the difference in selection bias between homogeneous and colony growth and therefore reduce power. Consequently, while we believe the experiments of Gralka *et al*. (2016) are currently the best model of colony-based microbial MA, similar experiments that better mimic the conditions of conventional microbial MA would be required to make firm conclusions.

Our meta-analysis of colony-based microbial MA experiments provides strong evidence for a deficit of non-synonymous mutations. While this conclusion appears to be at odds with the complete lack of statistically supported deficits of non-synonymous mutations in the literature (Table 1), we note that at least four studies erroneously reported non-significant results and there may well be a file drawer of significant findings never published - a rare reversal of the usual bias towards publishing significant findings (Rosenthal 1979). The observed deficit is best explained by a selection coefficient of *s* =-0.011 for non-synonymous mutations under the assumptions of the deterministic, homogeneous model of Wahl and Agashe (2022) and assuming synonymous mutations are neutral. The *s* required to produce the observed deficit is likely even more negative than this estimate suggests: synonymous sites are also likely to be subject to purifying selection (Ikemura 1981) and deterministic models, which ignore drift, overpredicted selection efficiency by up to 50% (*τ*/*τ*_*e*_ *>* 1.5) in our simulations, depending on the details of the homogeneous growth model. Direct estimates of mean *s* from multi-mutant lines range from *−*0.0021 (Robert et al. 2018) to *−*0.00055 (Baehr et al. 2025) (see Grosse-Sommer *et al*. (2026) for a review of estimates) which implies that the lower bound for the mean *s* of non-synonymous mutations lies between *−*0.0038 and *−*0.0009 (assuming 56% of mutations are non-synonymous and all synonymous mutations are neutral). Although these estimates have not corrected for selection bias, they are considerably smaller than the *s* required to generate the observed deficit of non-synonymous mutations under homogeneous growth, and it seems likely that increased selection efficiency due to colony growth is operating.

A remaining caveat is that many MA studies that test for a deficit of non-synonymous mutations either do not state the procedure used for determining the expected ratio in the absence of selection, or only control for a subset of possible confounders (Table 1). The impact of this is hard to assess. Because transversions are rarer than transitions in most species (after accounting for mutational opportunity - Lynch *et al*. (2023)) yet more likely to be non-synonymous, ignoring the difference between transition and transversion rates should overstate the deficit of non-synonymous mutations (Ina 1995) as seen in our models of *E. coli* mutations. However, those studies that reported controlling for differences in transition and transversion rates actually showed larger deficits in our meta-analysis. This maybe due to those studies more fully accounting for other aspects of the mutational spectrum, which if not accounted for are likely to result in the deficit of non-synonymous mutations being under-estimated since less constrained synonymous sites are likely to be closer to mutation-drift balance where the least mutable bases have higher equilibrium frequencies.

A deficit of coding mutations has been observed previously (see Table 1) and is recapitulated in our study of *E. coli*. At face value this mutational signature is consistent with purifying selection. However, because purifying selection is generally stronger on non-coding sites than on synonymous sites (Thorpe et al. 2017), selection-driven bias should be greater for non-coding mutations than for synonymous mutations, opposite to what we see (*β*_1_ is negative). A more likely explanation is that mechanisms have evolved to suppress mutations in coding regions. Transcription-coupled repair has been proposed as one such mechanism (Hanawalt and Spivak 2008; Ganesan *et al*. 2012) although in *E. coli*, inactivation of transcription-coupled repair through the deletion of *uvrA* did not cause a change in the coding/non-coding mutational spectrum (Foster et al. 2015). However, the bias towards non-coding mutations disappeared in lines in which the mismatch repair gene *MutL* was deleted (Lee et al. 2012) indicating that mismatch repair systems predominantly do target coding sequences.

While we believe that our simulations demonstrate qualitatively how selection operates in colony-based MA, developing a theoretical framework for accurately quantifying the degree of selection bias and controlling for it would be more challenging. Until this challenge is met, we suggest using existing homogeneous growth theory to account for selection bias, even though this theory is likely to underestimate the strength of bias. Developing more realistic theory is currently impeded by several factors. First, the cell-based and site-based models are extremes of a continuum and reality is likely somewhere in-between. Second, the increased efficacy of selection under cell-based colony growth is due to the increased number of generations that elapse per population doubling. Under the Eden model, the number of generations increases exponentially (Gralka *et al*. 2016; Fusco *et al*. 2016), and in our simulations, after 16 doublings the mean number of generations was 181. For *E*.*coli*, 16 doublings takes around nine hours under colony growth (Mahilkar *et al*. 2022), implying that in our simulations the average division time of a cell line is 2.98 minutes. The lack of realism probably arises from model constraints, where each cell can only divide into three adjacent lattice sites, producing highly asymmetric genealogies. A number of strategies could be employed that generate more symmetric genealogies and reduce the number of generations: allowing three-dimensional or diagonal growth (Colyer *et al*. 2024), allowing growth within an extended growth layer (Schreck *et al*. 2023) or imposing a time delay between subsequent reproduction events of the same cell. Introducing these components to the model is likely to reduce the differences between colony and homogeneous growth and increase the threshold number of doublings at which selection efficiency is greater under cell-based colony growth. However, such models will need to be informed by empirical measurements of the number of generations under colony based growth. These could be made directly by using cell tracking software that monitors individual cell lineages over time (Balomenos *et al*. 2015; Lamprecht *et al*. 2017), or alternatively, it may be possible to characterise the clone size distribution from sequencing subsamples of a colony (Schreck *et al*. 2023) and from this indirectly infer the number of generations.

Empirical measurements of the number of generations that elapse under colony growth would not only help with assessing, and ultimately correcting for selection bias, but also provide better estimates of the mutation rate. Most MA studies equate the number of population doublings with the number of generations. However, *τ* is a lower bound on the number of generations and is only achieved in a population where every cell divides synchronously. A consequence of this is that current estimates of the ‘per generation’ mutation rate will be downwardly biased, particularly in colony-based MA where the number of generations is expected to exceed *τ* by some margin. Similar issues have been noted when estimating mutation rates from fluctuation tests: the classical clone size distribution is described by the Luria–Delbrück distribution (Luria and Delbrück 1943; Lea and Coulson 1949), but asymmetric growth within colonies can cause deviations from this distribution (Fusco *et al*. 2016; Schreck *et al*. 2023). That being said, Baehr *et al*. (2025) conducted MA in *E. coli* using both liquid culture and colony propagation with very similar *τ* (29.9 and 26.9 respectively). The base-pair substitution rate per site per doubling was (non-significantly) 11% higher under colony growth compared to homogeneous growth suggesting that the increased number of generations under colony growth may only be marginally higher than under homogeneous growth. However, this conclusion rests on two assumptions. First, it is assumed that the downward bias in the mutation rate caused by the removal of deleterious mutations is the same under homogenous and colony growth. Since our results suggest selection efficiency may be greater under colony growth, the actual number of mutations may be more than 11% higher under colony growth compared to homogenous growth. Second, when inferring the number of generations from the number of mutations we have assumed the mutation rate *per generation* is the same under colony and homogeneous growth. If (some) mutations accumulate at a constant rate per unit time (both MA procedures ran for 24 hours between bottlenecks) this would tend to reduce the mutation rate per generation under colony growth. Similarly, cells will be at higher density in colonies than in liquid culture and Krašovec *et al*. (2017b) showed that *E. coli* has a lower mutation rate in higher density cultures. If the mutation rate per generation is lower under colony growth, then an 11% increase in the number of mutations per doubling implies that the number of generations per population doubling must exceed 11%.

In conclusion, we show that microbial MA consistently produces a deficit of non-synonymous mutations. The available evidence suggests this is because deleterious non-synonymous mutations are being removed by selection. The failure of many studies to control for mutational spectra when assessing non-synonymous deficits puts some doubt on the magnitude of the deficit caused by selection, but we develop a flexible multinomial model that could be used to address this in future studies. In order to measure the distribution of fitness effects of new mutations accurately it is imperative to correct for this selection bias. Although methods exist, they are built under the assumption that MA lines grow in a well-mixed culture rather than as colonies, as is typical. We show that selection bias under colony growth may be considerably different to that under well-mixed growth, but in order to develop methods that account for this selection bias, and indeed even being able to report mutation rates ‘per generation’, empirical experiments will need to be conducted to understand patterns of growth and selection in colonies better.

## Supporting information

Ecoli Data

Eden Model SI

Multinomial SI

## Data availability

The raw sequence data from our E. coli MA experiment has been uploaded to the NCBI Sequence Read Archive under BioProject ID PRJNA1466858. The code for mapping the reads and then calling and validating the mutations can be found at https://github.com/JGrosseSommer/Ecoli_MA. All other code can be found in the repository https://github.com/JGrosseSommer/Selection_MA.

## Acknowledgements

We thank Mrudula Sane and Deepa Agashe for sharing data and useful discussion, Rob Ness, Darren Obbard, Jobran Chebib and Helen Alexander for help and insight, and Mike Lynch and two anonymous reviewers for feedback on the manuscript. This work was supported by funding from NERC through an E4 DTP studentship.

1 Note that Mahilkar *et al*. (2022) did not interpret their results in this way. However, in their Figure 4 D the fitness decline after 150 transfers is ~0.01 ~0.04 and ~ 0.1, for colonies grown for 8, 16 and 24 hours, respectively. The greater fitness decline in larger colonies will mainly be due to more time/generations having elapsed and so more (deleterious) mutations accumulating. In Figure S18, Mahilkar *et al*. (2022) reports the number of cells in colonies grown for 8 hours (~10^5^), 16 hours (~5 *×*10^7^) and 24 hours (~8 *×* 10^8^) resulting in 17, 26 and 30 population doublings, respectively. Consequently, the fitness declines per population doubling are 0.0006, 0.0016 and 0.0034, respectively. Under the assumption that the number of doublings is equal to the number of generations, these numbers imply per-generation fitness declines in the opposite direction to that expected

## Notes

### Competing Interest Statement

The authors have declared no competing interest.

### Summary of Updates

New data added to the analysis of E. coli mutation patterns and some textual changes.

## Literature cited

Andersson, D. I., and D. Hughes, 1996 Muller’s ratchet decreases fitness of a DNA-based microbe. Proceedings of the National Academy of Sciences of the United States of America 93: 906–907.

Baehr, S., W. C. Ho, S. Perez, A. Cenzano, K. Hancock et al., 2025 Consideration of a liquid mutation-accumulation experiment to measure mutation rates by successive serial dilution. Genome Biology and Evolution 17: evaf049.

Balomenos, A. D., P. Tsakanikas, and E. S. Manolakos, 2015 Tracking single-cells in overcrowded bacterial colonies. In 37th annual international conference of the IEEE engineering in medicine and biology society (EMBC). Institute of Electrical and Electronics Engineers Inc., 6473–6476.

Bates, D., M. Mächler, B. M. Bolker, and S. C. Walker, 2015 Fitting linear mixed-effects models using lme4. Journal of Statistical Software 67: 1–48.

Behringer, M. G., and D. W. Hall, 2016 Genome-wide estimates of mutation rates and spectrum in Schizosaccharomyces pombe indicate CpG sites are highly mutagenic despite the absence of DNA methylation. G3: Genes, Genomes, Genetics 6: 149–160.

Bosshard, L., I. Dupanloup, O. Tenaillon, R. Bruggmann, M. Ackermann et al., 2017 Accumulation of deleterious mutations during bacterial range expansions. Genetics 207: 669–684.

Castañeda-García, A., I. Martín-Blecua, E. Cebrián-Sastre, A. Chiner-Oms, M. Torres-Puente et al., 2020 Specificity and mutagenesis bias of the mycobacterial alternative mismatch repair analyzed by mutation accumulation studies. Science Advances 6: 4453–4465.

Charlesworth, B., 2020 How long does it take to fix a favorable mutation, and why should we care? American Naturalist 195: 753–771.

Chen, Z., X. Wang, Y. Song, Q. Zeng, Y. Zhang et al., 2021 Prochlorococcus have low global mutation rate and small effective population size. Nature Ecology & Evolution 6: 183–194.

Colyer, B., M. Bak, D. Basanta, and R. Noble, 2024 A seven-step guide to spatial, agent-based modelling of tumour evolution. Evolutionary Applications 17: e13687.

Couce, A., A. Limdi, M. Magnan, S. V. Owen, C. M. Herren et al., 2024 Changing fitness effects of mutations through long-term bacterial evolution. Science 383.

Croissant, Y., 2020 Estimation of random utility models in R: the mlogit package. Journal of Statistical Software 95: 1–41.

Danecek, P., J. K. Bonfield, J. Liddle, J. Marshall, V. Ohan et al., 2021 Twelve years of SAMtools and BCFtools. GigaScience 10: giab008.

Dillon, M. M., and V. S. Cooper, 2016 The fitness effects of spontaneous mutations nearly unseen by selection in a bacterium with multiple chromosomes. Genetics 204: 1225–1238.

Dillon, M. M., W. Sung, M. Lynch, and V. S. Cooper, 2015 The rate and molecular spectrum of spontaneous mutations in the GC-rich multichromosome genome of Burkholderia cenocepacia. Genetics 200: 935–946.

Dillon, M. M., W. Sung, R. Sebra, M. Lynch, and V. S. Cooper, 2016 Genome-wide biases in the rate and molecular spectrum of spontaneous mutations in Vibrio cholerae and Vibrio fischeri. Molecular Biology and Evolution 34: 93–109.

Eden, M., 1961 A two-dimensional growth process. Proc. 4th Berkeley Sympos. Math. Statist. and Prob 4.4: 223–239.

Farlow, A., H. Long, S. Arnoux, W. Sung, T. G. Doak et al., 2015 The spontaneous mutation rate in the fission yeast Schizosaccharomyces pombe. Genetics 201: 737–744.

Farrell, F. D., M. Gralka, O. Hallatschek, and B. Waclaw, 2017 Mechanical interactions in bacterial colonies and the surfing probability of beneficial mutations. Journal of The Royal Society Interface 14: 20170073.

Fernández, J., and C. López-Fanjul, 1996 Spontaneous mutational variances and covariances for fitness-related traits in Drosophila melanogaster. Genetics 143: 829–837.

Foster, P. L., H. Lee, E. Popodi, J. P. Townes, and H. Tang, 2015 Determinants of spontaneous mutation in the bacterium Es-cherichia coli as revealed by whole-genome sequencing. Proceedings of the National Academy of Sciences of the United States of America 112: E5990–E5999.

Fusco, D., M. Gralka, J. Kayser, A. Anderson, and O. Hallatschek, 2016 Excess of mutational jackpot events in expanding populations revealed by spatial luria–delbrück experiments. Nature communications 7: 12760.

Ganesan, A., G. Spivak, and P. C. Hanawalt, 2012 Transcription-coupled DNA repair in prokaryotes. Progress in molecular biology and translational science 110: 25–40.

Gordo, I., and F. Dionisio, 2005 Nonequilibrium model for estimating parameters of deleterious mutations. Physical review E - Statistical, nonlinear, and soft matter physics 71: 031907.

Gralka, M., F. Stiewe, F. Farrell, W. Möbius, B. Waclaw et al., 2016 Allele surfing promotes microbial adaptation from standing variation. Ecology Letters 19: 889–898.

Grosse-Sommer, J. M., D. Newman, and J. D. Hadfield, 2026 Fitness effects of single spontaneous mutants are heavily confounded with non-genetic sources of variation in Escherichia coli. in prep.

Gu, J., X. Wang, X. Ma, Y. Sun, X. Xiao et al., 2021 Unexpectedly high mutation rate of a deep-sea hyperthermophilic anaerobic archaeon. The ISME Journal 15: 1862–1869.

Hallatschek, O., P. Hersen, S. Ramanathan, and D. R. Nelson, 2007 Genetic drift at expanding frontiers promotes gene segregation. Proceedings of the National Academy of Sciences of the United States of America 104: 19926–19930.

Hanawalt, P. C., and G. Spivak, 2008 Transcription-coupled DNA repair: two decades of progress and surprises. Nature Reviews Molecular Cell Biology 9: 958–970.

Hardesty, B., T. Obrig, J. Irvin, and W. Culp, 1973 The effect of sodium fluoride, edeine, and cycloheximide on peptide synthesis with reticulocyte ribosomes. Basic life sciences 1: 377–392.

Huber, W., V. J. Carey, R. Gentleman, S. Anders, M. Carlson et al., 2015 Orchestrating high-throughput genomic analysis with Bioconductor. Nature Methods 12: 115–121.

Ikemura, T., 1981 Correlation between the abundance of Escherichia coli transfer RNAs and the occurrence of the respective codons in its protein genes: A proposal for a synonymous codon choice that is optimal for the E. coli translational system. Journal of Molecular Biology 151: 389–409.

Imhof, M., and C. Schlötterer, 2001 Fitness effects of advantageous mutations in evolving Escherichia coli populations. Proceedings of the National Academy of Sciences of the United States of America 98: 1113–1117.

Ina, Y., 1995 New methods for estimating the numbers of synonymous and nonsynonymous substitutions. Journal of molecular evolution 40: 190–226.

Jago, M. J., R. Green, M. R. Czernuszka, S. Denisov, R. Krašovec et al., 2026 Extended sequence context shapes mutational bias in Escherichia coli. Proceedings of the National Academy of Sciences 123: e2601345123.

Joseph, S. B., and D. W. Hall, 2004 Spontaneous mutations in diploid Saccharomyces cerevisiae: More beneficial than expected. Genetics 168: 1817–1825.

Karlin, S., 1968 Rates of approach to homozygosity for finite stochastic models with variable population size. The American Naturalist 102: 443–455.

Katju, V., and U. Bergthorsson, 2019 Old trade, new tricks: Insights into the spontaneous mutation process from the partnering of classical mutation accumulation experiments with Julie Grosse-Sommer and Jarrod Hadfield 15 high-throughput genomic approaches. Genome Biology and Evolution 11: 136–165.

Katju, V., L. B. Packard, L. Bu, P. D. Keightley, and U. Bergthorsson, 2015 Fitness decline in spontaneous mutation accumulation lines of Caenorhabditis elegans with varying effective population sizes. Evolution 69: 104–116.

Keightley, P. D., and A. Caballero, 1997 Genomic mutation rates for lifetime reproductive output and lifespan in Caenorhabditis elegans. Proceedings of the National Academy of Sciences of the United States of America 94: 3823–3827.

Kibota, T. T., and M. Lynch, 1996 Estimate of the genomic mutation rate deleterious to overall fitness in E. coli. Nature 381: 694–696.

Kirkpatrick, M., and M. Slatkin, 1993 Searching for evolutionary patterns in the shape of a phylogenetic tree. Evolution 47: 1171–1181.

Krašovec, M., A. Eyre-Walker, S. Sanchez-Ferandin, and G. Piganeau, 2017a Spontaneous mutation rate in the smallest photosynthetic eukaryotes. Molecular Biology and Evolution 34: 1770–1779.

Krašovec, R., H. Richards, D. R. Gifford, C. Hatcher, K. J. Faulkner et al., 2017b Spontaneous mutation rate is a plastic trait associated with population density across domains of life. PLoS Biology 15: e2002731.

Kucukyildirim, S., M. Behringer, W. Sung, D. A. Brock, T. G. Doak et al., 2020 Low base-substitution mutation rate but high rate of slippage mutations in the sequence repeat-rich genome of Dictyostelium discoideum. G3: Genes, Genomes, Genetics 10: 3445–3452.

Kucukyildirim, S., H. Long, W. Sung, S. F. Miller, T. G. Doak et al., 2016 The rate and spectrum of spontaneous mutations in Mycobacterium smegmatis, a bacterium naturally devoid of the postreplicative mismatch repair pathway. G3: Genes, Genomes, Genetics 6: 2157–2163.

Kucukyildirim, S., S. F. Miller, and M. Lynch, 2021 Low base-substitution mutation rate and predominance of insertion-deletion events in the acidophilic bacterium Acidobacterium capsulatum. Ecology and Evolution 11: 17609–17614.

Kucukyildirim, S., H. O. Ozdemirel, and M. Lynch, 2023 Similar mutation rates but different mutation spectra in moderate and extremely halophilic archaea. G3: Genes, Genomes, Genetics 13: jkac303.

Lamprecht, S., E. M. Schmidt, C. Blaj, H. Hermeking, A. Jung et al., 2017 Multicolor lineage tracing reveals clonal architecture and dynamics in colon cancer. Nature communications 8: 1406.

Lavrentovich, M. O., M. E. Wahl, D. R. Nelson, and A. W. Murray, 2016 Spatially constrained growth enhances conversional meltdown. Biophysical Journal 110: 2800–2808.

Lawrence, M., R. Gentleman, and V. Carey, 2009 rtracklayer: an r package for interfacing with genome browsers. Bioinformatics 25: 1841–1842.

Lawrence, M., W. Huber, H. Pagès, P. Aboyoun, M. Carlson et al., 2013 Software for computing and annotating genomic ranges. PLoS Computational Biology 9.

Lea, D. E., and C. A. Coulson, 1949 The distribution of the numbers of mutants in bacterial populations. Journal of genetics 49: 264–285.

Lee, H., E. Popodi, H. Tang, and P. L. Foster, 2012 Rate and molecular spectrum of spontaneous mutations in the bacterium Escherichia coli as determined by whole-genome sequencing. Proceedings of the National Academy of Sciences of the United States of America 109: E2774–E2783.

Lesecque, Y., P. D. Keightley, and A. Eyre-Walker, 2012 A resolution of the mutation load paradox in humans. Genetics 191: 1321–1330.

Li, H., 2013 Aligning sequence reads, clone sequences and assembly contigs with BWA-MEM. arXiv : 1303.3997.

Li, H., 2018 Minimap2: pairwise alignment for nucleotide sequences. Bioinformatics 34: 3094–3100.

Liu, H., and J. Zhang, 2019 Yeast spontaneous mutation rate and spectrum are environment-dependent. Curr Biol 29: 1584–1591.

Long, H., M. G. Behringer, E. Williams, R. Te, and M. Lynch, 2016a Similar mutation rates but highly diverse mutation spectra in Ascomycete and Basidiomycete yeasts. Genome biology and evolution 8: 3815–3821.

Long, H., S. Kucukyildirim, W. Sung, E. Williams, H. Lee et al., 2015 Background mutational features of the radiation-resistant bacterium Deinococcus radiodurans. Molecular Biology and Evolution 32: 2383–2392.

Long, H., S. F. Miller, C. Strauss, C. Zhao, L. Cheng et al.,2016b Antibiotic treatment enhances the genome-wide mutation rate of target cells. Proceedings of the National Academy of Sciences 113: E2498–E2505.

López-Cortegano, E., R. J. Craig, J. Chebib, T. Samuels, A. D. Morgan et al., 2021 De novo mutation rate variation and its determinants in Chlamydomonas. Molecular Biology and Evolution 38: 3709–3723.

Luria, S. E., and M. Delbrück, 1943 Mutations of bacteria from virus sensitivity to virus resistance. Genetics 28: 491.

Lynch, M., F. Ali, T. Lin, Y. Wang, J. Ni et al., 2023 The divergence of mutation rates and spectra across the Tree of Life. EMBO reports 24: e57561.

Lynch, M., L. Latta, J. Hicks, and M. Giorgianni, 1998 Mutation, selection and the maintenance of life-history variation in a natural population. Evolution 52: 727–733.

Lynch, M., W. Sung, K. Morris, N. Coffey, C. R. Landry et al., 2008 A genome-wide view of the spectrum of spontaneous mutations in yeast. Proceedings of the National Academy of Sciences of the United States of America 105: 9272–9277.

MacLean, R. C., and A. Buckling, 2009 The distribution of fitness effects of beneficial mutations in Pseudomonas aeruginosa. PLoS genetics 5: e1000406.

Mahilkar, A., N. Raj, S. Kemkar, and S. Saini, 2022 Selection in a growing colony biases results of mutation accumulation experiments. Scientific Reports 12: 1–12.

Mettrop, L., A. Lipzen, C. Vandecasteele, C. Eché, A. Labécot et al., 2025 Low mutation rate and atypical mutation spectrum in Prasinoderma coloniale: insights from an early diverging green lineage. Genome Biology and Evolution 17: evaf026.

Mukai, T., 1964 The genetic structure of natural populations of Drosophila melanogaster. I. Spontaneous mutation rate of polygenes controlling viability. Genetics 50: 1–19.

Muller, H. J., 1928 The measurement of gene mutation rate in Drosophila, its high variability, and its dependence upon temperature. Genetics 13: 279–357.

Nadell, C. D., K. Drescher, and K. R. Foster, 2016 Spatial strucre, cooperation and competition in biofilms. Nature Reviews Microbiology 14: 589–600.

Ness, R. W., A. D. Morgan, N. Colegrave, and P. D. Keightley, 2012 Estimate of the spontaneous mutation rate in Chlamydomonas reinhardtii. Genetics 192: 1447–1454.

Ness, R. W., A. D. Morgan, R. B. Vasanthakrishnan, N. Colegrave, and P. D. Keightley, 2015 Extensive de novo mutation rate variation between individuals and across the genome of Chlamydomonas reinhardtii. Genome Research 25: 1739–1749.

Nguyen, D. T., B. Wu, H. Long, N. Zhang, C. Patterson et al., 2020 Variable spontaneous mutation and loss of heterozygosity among heterozygous genomes in yeast. Molecular Biology and Evolution 37: 3118–3130.

Obenchain, V., M. Lawrence, V. Carey, S. Gogarten, P. Shannon et al., 2014 VariantAnnotation: a Bioconductor package for exploration and annotation of genetic variants. Bioinformatics 30: 2076–2078.

Otto, S. P., and M. E. Orive, 1995 Evolutionary consequences of mutation and selection within an individual. Genetics 141: 1173–1187.

Pan, J., W. Li, J. Ni, K. Wu, I. Konigsberg et al., 2022 Rates of mutations and transcript errors in the foodborne pathogen Salmonella enterica subsp. enterica. Molecular Biology and Evolution 39: msac081.

Pan, J., E. Williams, W. Sung, M. Lynch, and H. Long, 2021 The insect-killing bacterium Photorhabdus luminescens has the lowest mutation rate among bacteria. Marine Life Science and Technology 3: 20–27.

Peck, J. R., G. Barreau, and S. C. Heath, 1997 Imperfect genes, fisherian mutation and the evolution of sex. Genetics 145: 1171–1199.

Puigbò, P., A. Romeu, and S. Garcia-Vallvé, 2008 HEG-DB: A database of predicted highly expressed genes in prokaryotic complete genomes under translational selection. Nucleic Acids Research 36: D524–D527.

R Core Team, 2025 R: A Language and Environment for Statistical Computing. R Foundation for Statistical Computing, Vienna, Austria.

Robert, L., J. Ollion, J. Robert, X. Song, I. Matic et al., 2018 Mutation dynamics and fitness effects followed in single cells. Science 359: 1283–1286.

Robinson, J. T., H. Thorvaldsdóttir, W. Winckler, M. Guttman, E. S. Lander et al., 2011 Integrative genomics viewer. Nature Biotechnology 29: 24–26.

Rosenthal, R., 1979 The file drawer problem and tolerance for null results. Psychological bulletin 86: 638.

Sane, M., G. D. Diwan, B. A. Bhat, L. M. Wahl, and D. Agashe, 2023 Shifts in mutation spectra enhance access to beneficial mutations. Proceedings of the National Academy of Sciences 120: e2207355120.

Sane, M., J. J. Miranda, and D. Agashe, 2018 Antagonistic pleiotropy for carbon use is rare in new mutations. Evolution 72: 2202–2213.

Schreck, C. F., D. Fusco, Y. Karita, S. Martis, J. Kayser et al., 2023 Impact of crowding on the diversity of expanding populations. Proceedings of the National Academy of Sciences 120: e2208361120.

Schroeder, J. W., W. G. Hirst, G. A. Szewczyk, and L. A. Simmons, 2016 The effect of local sequence context on mutational bias of genes encoded on the leading and lagging strands. Current Biology 26: 692–697.

Senra, M. V., W. Sung, M. Ackerman, S. F. Miller, M. Lynch et al., 2018 An unbiased genome-wide view of the mutation rate and spectrum of the endosymbiotic bacterium Teredinibacter turnerae. Genome Biology and Evolution 10: 723–730.

Shao, K. T., and R. R. Sokal, 1990 Tree balance. Systematic Biology 39: 266–276.

Sharp, N. P., L. Sandell, C. G. James, and S. P. Otto, 2018 The genome-wide rate and spectrum of spontaneous mutations differ between haploid and diploid yeast. Proceedings of the National Academy of Sciences of the United States of America 115: E5046–E5055.

Shewaramani, S., T. J. Finn, S. C. Leahy, R. Kassen, P. B. Rainey et al., 2017 Anaerobically grown Escherichia coli has an enhanced mutation rate and distinct mutational spectra. PLOS Genetics 13: e1006570.

Strauss, C., H. Long, C. E. Patterson, R. Te, and M. Lynch, 2017 Genome-wide mutation rate response to pH change in the coral reef pathogen Vibrio shilonii AK1. mBio 8: 10–1128.

Sun, Y., K. E. Powell, W. Sung, M. Lynch, M. A. Moran et al., 2017 Spontaneous mutations of a model heterotrophic marine bacterium. The ISME Journal 11: 1713–1718.

Sung, W., M. S. Ackerman, J. F. Gout, S. F. Miller, E. Williams et al., 2015 Asymmetric context-dependent mutation patterns revealed through mutation-accumulation experiments. Molecular Biology and Evolution 32: 1672–1683.

Sung, W., M. S. Ackerman, S. F. Miller, T. G. Doak, and M. Lynch, 2012 Drift-barrier hypothesis and mutation-rate evolution. Proceedings of the National Academy of Sciences of the United States of America 109: 18488–18492.

Takemoto, N., I. Numata, M. Su’Etsugu, and T. Miyoshi-Akiyama, 2018 Bacterial EndoMS/NucS acts as a clampmediated mismatch endonuclease to prevent asymmetric accumulation of replication errors. Nucleic Acids Research 46: 6152–6165.

Thorpe, H. A., S. C. Bayliss, L. D. Hurst, and E. J. Feil, 2017 Comparative analyses of selection operating on nontranslated intergenic regions of diverse bacterial species. Genetics 206: 363–376.

Tincher, C., H. Long, M. Behringer, N. Walker, and M. Lynch, 2017 The glyphosate-based herbicide roundup does not elevate genome-wide mutagenesis of Escherichia coli. G3: Genes, Genomes, Genetics 7: 3331–3335.

Wahl, L. M., and D. Agashe, 2022 Selection bias in mutation accumulation. Evolution 76: 528–540.

Walsh, B., and M. Lynch, 2018 Evolution and selection of quantitative traits. Oxford University Press.

Wang, Y., and D. J. Obbard, 2023 Experimental estimates of germline mutation rate in eukaryotes: a phylogenetic metaanalysis. Evolution Letters 7: 216–226.

Wei, W., W. C. Ho, M. G. Behringer, S. F. Miller, G. Bcharah et al., 2022 Rapid evolution of mutation rate and spectrum in response to environmental and population-genetic challenges. Nature Communications 13: 1–10.

Xue, C. X., H. Zhang, H. Y. Lin, Y. Sun, D. Luo et al., 2020 Ancestral niche separation and evolutionary rate differentiation between sister marine flavobacteria lineages. Environmental Microbiology 22: 3234–3247.

Yang, Z., and R. Nielsen, 2000 Estimating synonymous and nonsynonymous substitution rates under realistic evolutionary models. Molecular biology and evolution 17: 32–43.

Zarate, S., A. Carroll, M. Mahmoud, O. Krasheninina, G. Jun et al., 2021 Parliament2: Accurate structural variant calling at scale. GigaScience 9: giaa145.

Zeyl, C., and J. A. G. M. DeVisser, 2001 Estimates of the rate and distribution of fitness effects of spontaneous mutation in Saccharomyces cerevisiae. Genetics 157: 53–61.

Zhu, Y. O., M. L. Siegal, D. W. Hall, and D. A. Petrov, 2014 Precise estimates of mutation rate and spectrum in yeast. Proceedings of the National Academy of Sciences of the United States of America 111: E2310–E2318.

