## Supplementary material for "Selection bias in microbial mutation accumulation studies and the impact of colony growth": Ecoli Data: Ecoli_Data.html

Aggregating Escherichia coli Mutation Accumulation Data


### Aggregating *Escherichia coli* Mutation Accumulation Data

- 1 Co-ordinate conversion
- 2 Aggregate mutations from all studies
- 3 Initialise data-frame
- 4 Annotate sites/variants
- 5 Add mutations

In this workbook we take base-pair mutations accumulated in our own mutation accumulation experiment, and combine them with base-pair mutations from five different mutation accumulation experiments in *Escherichia coli* (Foster et al. (2015), Long et al. (2016), Tincher et al. (2017), Sane, Miranda, and Agashe (2018), Wei et al. (2022)). We then create a data frame which allows us to analyse patterns mutation using a multinomial model (see Multinomial\_Model\_SI).

### 1 Co-ordinate conversion

All previous studies used the same reference genome as us (NC\_000913.3), except Foster et al. (2015) who used an earlier reference genome (NC\_000913.2). In order to map their positions to those in the new reference we used minimap2 (v2.31) Li (2018) to align the two assemblies and then a paftools.js liftover - this is implemented in bash script convert\_foster.sh.

```
ref=~/Work/Selection_MA/Data/Raw
out=~/Work/Selection_MA/Data/Intermediate 

oldChr=NC_000913.2
# contig name

CSV=BPS_WT_Foster_et_al_2015.csv
OUT=lifted_coords.bed

minimap2 -x asm5 -c --cs $ref/GCF_000005845.2_ASM584v2_genomic.fna $ref/NC_000913.2.fna  > "$out/913.2_vs_913.3.paf"
# aligns the two genomes

awk -F, 'NR==1{next} {start=$2-1; end=$2; print "'$oldChr'", start, end, $0}' OFS="\t" "$ref/$CSV" > "$out/old_coords.bed"
# takes old coordinates from a csv file (column 2) and converts them to a bed file (counts from zero, tab-delimited)

paftools.js liftover -l 0 "$out/913.2_vs_913.3.paf" "$out/old_coords.bed" > "$out/$OUT"

rm "$out/913.2_vs_913.3.paf"
rm "$out/old_coords.bed"
```

The second and third columns of `lifted_coords.bed` give the old and new co-ordinates. A conda environment can be set up to allow paftools.js to run:

```
name: co-ord
channels:
  - bioconda
  - conda-forge
  - defaults
dependencies:
  - k8=1.2=hda5e58c_6
  - libcxx=22.1.6=h55c6f16_0
  - libzlib=1.3.2=h8088a28_2
  - minimap2=2.31=h6bd33b9_0
prefix: /opt/homebrew/Caskroom/miniconda/base/envs/co-ord
```

and the bash script can executed

```
system("conda activate co-ord"))
system(paste0("cd ", file.path(root, "R"))
system("./convert_foster.sh")
```

### 2 Aggregate mutations from all studies

The following code takes information from the supplementary materials of the source publications and organises them into a single data frame. Since we can’t host the supplementary material of the source publications on GitHub, this code will not run, but can be viewed. All mutation data are aggregated into the data frame `all_mutations.csv`, which can be found in `Data/Raw/`. Note that duplicate mutations due to contamination are retained in this file and filtered out later.

```
library(Biostrings)
gff_file <- file.path(root, "Data/Raw/GCF_000005845.2_ASM584v2_genomic.gff")
fna_file <- file.path(root, "Data/Raw/GCF_000005845.2_ASM584v2_genomic.fna")    

# path to gff genome annotation file and FASTA nucleotide sequence for the reference

bases<-c("A", "C", "T", "G")

complement<-function(x){
  bases<-c("A", "C", "T", "G")
  c("T", "G", "A", "C")[match(x, bases)]
}

genome <- readDNAStringSet(fna_file)

sequence <- strsplit(as.character(genome[[1]]), "")[[1]]
complement_sequence<-complement(sequence)


GrosseSommer<-read.csv("~/Work/Ecoli_MA/meta/real_mutations.csv")
GrosseSommer<-subset(GrosseSommer, nchar(REF)==nchar(ALT) & line!=95 & !duplicated(POS)) # bps only
# our mutations

Foster<-read.csv(file.path(root, "Data/Raw/BPS_WT_Foster_et_al_2015.csv"))

foster_map<-read.table(file.path(root, "Data/Intermediate/lifted_coords.bed"))

if(any(foster_map[,6]!="+")){stop("foster_map has a strand change")}

Foster$POS<-foster_map[,3]
# Turns Foster's co-ordinates into NC_000913.3 co-ordinate

colnames(Foster)[colnames(Foster)=="ConsensusBase"]<-"REF"
colnames(Foster)[colnames(Foster)=="ObservedBase"]<-"ALT"
# Give POS/REF/ALT same names

# table(sequence[Foster$POS], Foster$REF)
# make sure reference base reported is correct

Sane<-read.csv(file.path(root, "Data/Raw/Sane_et_al_2018.csv"))
Sane<-subset(Sane, nchar(orig_base)==nchar(mut_base)) # bps only

colnames(Sane)[colnames(Sane)=="genome_position"]<-"POS"
colnames(Sane)[colnames(Sane)=="orig_base"]<-"REF"
colnames(Sane)[colnames(Sane)=="mut_base"]<-"ALT"
# Give POS/REF/ALT same names

# table(sequence[Sane$POS], Sane$REF)
# make sure reference base reported is correct

Wei<-read.csv(file.path(root, "Data/Raw/BPS_WT_Wei_et_al_2022.csv"))
colnames(Wei)[colnames(Wei)=="Position"]<-"POS"

Wei$REF<-unlist(lapply(strsplit(Wei$Base.mutation, ":"), function(x)x[1]))
Wei$ALT<-unlist(lapply(strsplit(Wei$Base.mutation, ">|:"), function(x)x[3]))

Wei$ALT[which(Wei$REF=="G" & sequence[Wei$POS]=="C")]<-complement(Wei$ALT[which(Wei$REF=="G" & sequence[Wei$POS]=="C")])
Wei$REF[which(Wei$REF=="G" & sequence[Wei$POS]=="C")]<-"C"

Wei$ALT[which(Wei$REF=="A" & sequence[Wei$POS]=="T")]<-complement(Wei$ALT[which(Wei$REF=="A" & sequence[Wei$POS]=="T")])
Wei$REF[which(Wei$REF=="A" & sequence[Wei$POS]=="T")]<-"T"

# table(sequence[Wei$POS], Wei$REF)
# make sure reference base reported in each study is correct

Long<-read.csv(file.path(root, "Data/Raw/Long_et_al_2016.csv"), skip=1)
Long<-subset(Long, Norfloxacin.concentration..ng.ml.==0)
# subset for control samples

Long$POS<-as.numeric(unlist(lapply(strsplit(Long$Position, ":"), function(x)x[2])))
Long$REF<-unlist(lapply(strsplit(Long$Mutation, ">"), function(x)x[1]))
Long$ALT<-unlist(lapply(strsplit(Long$Mutation, ">"), function(x)x[2]))

# table(sequence[Long$POS], Long$REF)
# make sure reference base reported is correct

Tincher<-read.csv(file.path(root, "Data/Raw/Tincher_et_al_2017.csv"), skip=1)
Tincher<-subset(Tincher, grepl("_A", Samples))

Tincher$POS<-as.numeric(unlist(lapply(strsplit(Tincher$Genomic.position, ":"), function(x)x[2])))
Tincher$REF<-unlist(lapply(strsplit(Tincher$Base.substitution, ">"), function(x)x[1]))
Tincher$ALT<-unlist(lapply(strsplit(Tincher$Base.substitution, ">"), function(x)x[2]))

# table(sequence[Tincher$POS], Tincher$REF)
# make sure reference base reported is correct

all_mutations<-rbind(Foster[,c("POS", "REF", "ALT")], GrosseSommer[,c("POS", "REF", "ALT")], Sane[,c("POS", "REF", "ALT")], Wei[,c("POS", "REF", "ALT")], Long[,c("POS", "REF", "ALT")], Tincher[,c("POS", "REF", "ALT")])

all_mutations$source<-c(rep("Foster_2015", nrow(Foster)), rep("GrosseSommer_2026", nrow(GrosseSommer)), rep("Sane_2018", nrow(Sane)), rep("Wei_2022", nrow(Wei)), rep("Long_2016", nrow(Long)), rep("Tincher_2017", nrow(Tincher)))

write.csv(all_mutations, file.path(root, "Data/Raw/all_mutations.csv"), row.names=FALSE, quote=FALSE)
```

### 3 Initialise data-frame

We can start by creating a data frame of each of the possible bases at each site. To this data frame we will add those variables that can be obtained from the sequence without annotation.

```
library(Biostrings)
library(data.table)
library(VariantAnnotation)  
library(GenomicFeatures) 
library(rtracklayer)      
library(txdbmaker)
library(GenomeInfoDb)

gff_file <- file.path(root, "Data/Raw/GCF_000005845.2_ASM584v2_genomic.gff")

fna_file <- file.path(root, "Data/Raw/GCF_000005845.2_ASM584v2_genomic.fna")    

# path to gff genome annotation file and FASTA nucleotide sequence for the reference

bases<-c("A", "C", "T", "G")

complement<-function(x){
  bases<-c("A", "C", "T", "G")
  c("T", "G", "A", "C")[match(x, bases)]
}

genome <- readDNAStringSet(fna_file)

sequence <- strsplit(as.character(genome[[1]]), "")[[1]]
complement_sequence<-complement(sequence)

chrom <- "NC_000913.3"
len <- length(genome[[1]])

pos <- seq_along(sequence)

long_dat<-data.frame(base=rep(bases, len), REF_NUC=rep(sequence, each=4), site=rep(pos, each=4))

direction<-c("CW", "CCW")[(pos>1603057 & pos<(1603057+len/2))+1]
# specify whether reference is leading (CW) or lagging (CCW)

leading_strand<-sequence
leading_strand[which(direction=="CCW")]<-complement_sequence[which(direction=="CCW")]
# if reference is lagging, leading base needs to be complement of reference

leading_minus_neighbor<-leading_plus_neighbor<-rep(NA, len)
# vectors for 5' and 3' neighbours on leading strand

leading_minus_neighbor[which(direction=="CW")]<-c(NA, sequence[pos[which(direction=="CW")]-1])
# for leading reference use preceding reference base (not sequence[0] is empty so add NA)

leading_minus_neighbor[1]<-leading_strand[len]
# since genome circular minus one neighbour for first base is the final base

leading_minus_neighbor[which(direction=="CCW")]<-complement_sequence[pos[which(direction=="CCW")]+1]
# for lagging reference use preceding complementary base

leading_plus_neighbor[which(direction=="CW")]<-sequence[pos[which(direction=="CW")]+1]
# for leading reference use next reference base (not sequence[0] is empty so add NA)

leading_plus_neighbor[len]<-leading_strand[1]
# since genome circular plus one neighbour of last base is the first base

leading_plus_neighbor[which(direction=="CCW")]<-complement_sequence[pos[which(direction=="CCW")]-1]
# for lagging reference use next complementary base

long_dat$direction<-rep(direction, each=4)
long_dat$leading_strand<-rep(leading_strand, each=4)
long_dat$leading_T<-as.numeric(long_dat$leading_strand=="T")
long_dat$leading_G<-as.numeric(long_dat$leading_strand=="G")
long_dat$leading_minus_neighbor<-rep(leading_minus_neighbor, each=4)
long_dat$leading_plus_neighbor<-rep(leading_plus_neighbor, each=4)

# set up variables for multinomial model (see below)

long_dat$AT_nb<-rep((leading_minus_neighbor=="A")+(leading_plus_neighbor=="T"), each=4)
long_dat$TA_nb<-rep((leading_minus_neighbor=="T")+(leading_plus_neighbor=="A"), each=4)
long_dat$GC_nb<-rep((leading_minus_neighbor=="G")+(leading_plus_neighbor=="C"), each=4)
long_dat$CG_nb<-rep((leading_minus_neighbor=="C")+(leading_plus_neighbor=="G"), each=4)

long_dat$trans<-c("transversion", "transition")[with(long_dat, (REF_NUC=="A" & base=="G") | (REF_NUC=="G" & base=="A") | (REF_NUC=="C" & base=="T") | (REF_NUC=="T" & base=="C") )+1]
# assign mutations to transition or transversion (note reference==base is classed as a transversion but this is ignored in the analysis - JARROD once everything agrees with Julie's script add REF_NUC==base to logical statement)

long_dat$comb_REF<-rep(c("AT", "CG")[(sequence=="C" | sequence=="G")+1], each=4)
# variables for multinomial model (see below)

long_dat$mutate<-with(long_dat, as.numeric(base!=REF_NUC))
# indicate whether the base would be a mutation
```

### 4 Annotate sites/variants

Next we can add information about whether mutations are in coding regions, and if they are, the whether they are synonymous or non-synonymous and in highly expressed genes or not. To do this we create four vcf files, each of which has one of the four bases as the variant at every site in the genome.

```
txdb <- makeTxDbFromGFF(gff_file, format = "gff")
# create database of genomic features

gff <- import(gff_file)
# import the gff file

cds_gr <- cds(txdb,  columns = c("gene_id", "tx_name"))
# extract coding sequence regions

pseudo_gr <- gff[gff$type=="pseudogene"]
rrna_gr <- gff[gff$type=="rRNA"]
is_gr <- gff[grepl("insertion sequence", gff$mobile_element_type)]
gene_gr <- gff[gff$type=="gene"]

# filter gff files for pseudogenes, rrna, insertion sequences and genes

fa <- FaFile(fna_file)  
# creates a FaFile object for efficiently accessing the FASTA file

is_nonsyn<-matrix("synonymous", 4, len)
rownames(is_nonsyn)<-bases
# matrix for writing mutation type for each base change

for(i in 1:4){

    # create a vcf file where al variants are base[i]

    vcf_file <- file.path(root, "Data/Intermediate", paste0(bases[i], ".vcf"))

    con <- file(vcf_file, open = "w")

    writeLines("##fileformat=VCFv4.2", con)
    writeLines(sprintf("##contig=<ID=%s,length=%d>", chrom, len), con)
    writeLines("#CHROM\tPOS\tID\tREF\tALT\tQUAL\tFILTER\tINFO", con)

    lines <- paste(chrom,pos,".",sequence,bases[i],".","PASS",".",sep = "\t")
    
    writeLines(lines, con)

    close(con)

    vcf <- readVcf(vcf_file)
    # read in the vcf file

    vr  <- granges(vcf)                         # get the coordinates of the variants

    if(i==1){
      # since site invariant only needs to be computed once

      is_cds<-(countOverlaps(vr, cds_gr)>0 & !(countOverlaps(vr, pseudo_gr)>0))
      # logical: is site in coding sequence and not a pseudogene (note insertion sequences are retained)

      hits    <- findOverlaps(vr, gene_gr)
      # finds overlap between variant coordinates and genes
    
      gene<-rep(NA, len)
      gene[queryHits(hits)] <- unlist(gene_gr$locus_tag[subjectHits(hits)])

      long_dat$coding_type<-rep(c("intergenic", "protein_coding")[is_cds+1], each=4)
      long_dat$gene<-rep(gene, each=4)

      heg<-read.csv(file.path(root, "Data/Raw/HEGE.coli.txt"), skip=3, sep="\t")
      # read in data frame of highly expressed genes obtained from https://ppuigbo.me/programs/HEG-DB/

      long_dat$HEG<-long_dat$gene%in%heg$Gene
      # logical: is the site in a gene that is highly expressed 
      # which(!heg$X%in%long_dat$gene) some genes in heg seem to be missing.
    }

    coding <- predictCoding(vcf, txdb, seqSource = fa)
    # gets coding mutation types (synonymous, non-synonymous, )

    is_nonsyn[i,match(names(coding), names(vcf))]<-as.character(coding$CONSEQUENCE)
    # writes mutation type for mutations in coding sequence

    system(paste("rm", vcf_file))
}

long_dat$nonsyn<-as.numeric(c(is_nonsyn)!="synonymous")
long_dat$nonsyn[which(long_dat$coding_type=="intergenic")]<-0
```

### 5 Add mutations

Next we need to add the mutations to the data frame

```
all_mutations<-read.csv(file.path(root, "Data/Raw/all_mutations.csv"))

# read in all mutations (note Foster's mutations are reported for NC_000913.3 co-ordinates)

duplicates<-with(all_mutations, table(paste(POS,ALT) , source))

cs_duplicates<-duplicates[which(rowSums(duplicates>0)>1),]
# cross_study_duplicates

# There are two cross-study duplicates: 

subset(all_mutations, paste(POS, ALT)%in%rownames(cs_duplicates))
```

```
##         POS REF ALT            source
## 21   236978   T   G       Foster_2015
## 308 3946620   A   G GrosseSommer_2026
## 725  236978   T   G          Wei_2022
## 767 3946620   A   G          Wei_2022
```

```
ws_duplicates<-duplicates[which(rowSums(duplicates>1)>0),]

# There are many within-study duplicates, likely due to contamination. Note that some studies may have already discarded duplicates (for example we had one duplicate)

colSums(ws_duplicates>1)
```

```
##       Foster_2015 GrosseSommer_2026         Long_2016         Sane_2018 
##                 2                 0                 0                61 
##      Tincher_2017          Wei_2022 
##                 0                 5
```

```
mutations<-subset(all_mutations, !duplicated(paste(POS, ALT)))
# remove all duplications

# There's also a single site that mutated twice but to a different base

subset(mutations, POS%in%POS[duplicated(POS)])
```

```
##         POS REF ALT   source
## 701 4171941   C   A Wei_2022
## 779 4171941   C   G Wei_2022
```

```
mutations<-subset(mutations, POS!=4171941 | ALT!="A")
# We randomly retained the C to G mutation

long_dat$new_seq<-rep(0, len*4)

long_dat$new_seq<-with(long_dat, as.numeric(base==REF_NUC & !site%in%mutations$POS))
# sites where mutations didn't happen

long_dat$new_seq[(mutations$POS-1)*4+match(mutations$ALT, bases)]<-1
# sites where mutations did happen

write.csv(long_dat, file.path(root, "Data/Intermediate/long_dat.csv"), row.names=FALSE, quote=FALSE)
```

Foster, Patricia L, Heewook Lee, Ellen Popodi, Jesse P Townes, and Haixu Tang. 2015. “Determinants of Spontaneous Mutation in the Bacterium Escherichia Coli as Revealed by Whole-Genome Sequencing.” *Proceedings of the National Academy of Sciences* 112 (44): E5990–99.

Li, Heng. 2018. “Minimap2: Pairwise Alignment for Nucleotide Sequences.” *Bioinformatics* 34 (18): 3094–3100.

Long, Hongan, Samuel F Miller, Chloe Strauss, Chaoxian Zhao, Lei Cheng, Zhiqiang Ye, Katherine Griffin, et al. 2016. “Antibiotic Treatment Enhances the Genome-Wide Mutation Rate of Target Cells.” *Proceedings of the National Academy of Sciences* 113 (18): E2498–2505.

Sane, Mrudula, Joshua John Miranda, and Deepa Agashe. 2018. “Antagonistic Pleiotropy for Carbon Use Is Rare in New Mutations.” *Evolution* 72 (10): 2202–13.

Tincher, Clayton, Hongan Long, Megan Behringer, Noah Walker, and Michael Lynch. 2017. “The Glyphosate-Based Herbicide Roundup Does Not Elevate Genome-Wide Mutagenesis of *Escherichia Coli*.” *G3: Genes, Genomes, Genetics* 7 (10): 3331–35.

Wei, Wen, Wei Chin Ho, Megan G. Behringer, Samuel F. Miller, George Bcharah, and Michael Lynch. 2022. “Rapid evolution of mutation rate and spectrum in response to environmental and population-genetic challenges.” *Nature Communications* 13 (1): 1–10. https://doi.org/10.1038/s41467-022-32353-6.
