## Supplementary material for "Selection bias in microbial mutation accumulation studies and the impact of colony growth": Eden Model SI: Eden_Model_SI.html

Simulating Microbial Mutation-accumulation Experiments


### Simulating Microbial Mutation-accumulation Experiments

- 1 Introduction
  - 1.1 Colony Growth
  - 1.2 Well-mixed Growth
  - 1.3 Spontaneous mutations versus segregating variation
- 2 Density function and random-number generator for the DFE.
- 3 Growth Simulation Functions
  - 3.1 Colony growth
  - 3.2 Well-mixed growth
- 4 Mutation-accumulation Simulation Function
- 5 Functions for assessing the magnitude and impact of selection bias
- 6 Simulations
  - 6.1 Colony growth simulations
    - 6.1.1 Cell-based simulations
    - 6.1.2 Site-based simulations
  - 6.2 Well-mixed growth simulations
    - 6.2.1 Poisson simulations
    - 6.2.2 Doubling simulations
    - 6.2.3 Singe-step simulations
- 7 Weak Selection
- 8 Segregating Variation
- 9 Generation Time
  - 9.1 Scaling of selection efficiency with \(\tau\)
- 10 Meta-analysis of non-synonymous/synonymous ratios
  - 10.1 Standard meta-analysis
  - 10.2 Selection bias informed meta-analysis
- 11 Power
- References

### 1 Introduction

In this workbook we implement code that simulates the Eden model of bacterial colony growth, allowing for mutations that change a cell’s fitness. The aim is to characterise how much selection bias occurs in typical mutation-accumulation lines when inferring the distribution of fitness effects (DFE). Previous theory tackling this problem has assumed that the population of cells is well mixed during the growth phase, but selection dynamics are likely to be different under colony growth. At the end of the workbook we detail a meta-analysis of non-synonymous versus synonymous mutation rates in microbial mutation-accumulation experiments.

#### 1.1 Colony Growth

The Eden (1961) model serves as a basis for modelling colony growth where new cells are added at the colony boundary. Colyer et al. (2024) provide a nice overview of agent-based modelling of colony growth in the context of tumour evolution. A perimeter site is defined as a site that is empty but is adjacent to an occupied site. An occupied site can have up to four perimeter sites, although if cell death is not allowed, the maximum number of perimeter sites is three, since one is occupied by the cell’s mother. We will call any cell/occupied site with a free perimeter site, an edge cell/site. When choosing which cell reproduces and where its daughter is placed we can either sample an edge cell (cell-based rule) or sample a perimeter site (site-based rule). With selection, a simple cell-based rule would be to sample edge cells proportional to their fitnesses minus one (and possibly how many perimeter sites they have - referred to as a bond-based rule in Colyer et al. (2024)) and place the daughter in a random adjacent perimeter site. With selection, a simple site-based rule would be to place a daughter in a random perimeter site, and then sample a mother from the pool of adjacent mothers proportional to their fitnesses minus one. We use fitness minus one (\(w-1\)) since we do not allow cell death. Consequently, the expected number of daughters produced by each mother is \(w-1\), and since the mother itself survives, the total number of descendants is then equal to \(w\) in the next generation, consistent with a definition of fitness.

*Cell-based*: The algorithm works by storing information on all *edge cells* in the matrix, `M`. The fitness of the cell and its co-ordinates are stored in addition to whether (1) or not (0) the four possible adjacent sites are occupied or not (in the order right, below, left, above). An initial cell is placed at (0,0) with a fitness of two and a new cell is added each iteration until the final population size, \(N\), is reached. Each iteration, a mother is sampled from the edge cells with probability proportional to their fitness minus one, and the daughter is placed randomly in an empty adjacent site. The daughter inherits the mothers fitness multiplied by \(1\) or \(1+s\) with probability \(1-\mu\) or \(\mu\), respectively. \(s\) is drawn from the DFE, which we assume to be a mean-shifted reflected gamma distribution. The mother’s fitness is then updated following the same rule. When the daughter’s site is chosen, the occupancy of adjacent sites changes for up to three edge cells surrounding the daughter, and this information is updated in `M`. If the daughter fills the final empty site of an edge cell, the cell is removed from `M` (`M` is resized) and its information stored in `M_final`, a matrix with `N` rows that is progressively filled and contains information on the whole population by the end of the simulation. If the daughter falls in a site with at least one empty adjacent site it is appended to `M`, otherwise it is also added to `M_final`. The costly bits are probably resizing `M` and identifying those cells that lose an empty adjacent site when a daughter is added. The algorithm seems to scale with \(N^{4/3}\) and certainly could be made much more efficient.

*Site-based*: The algorithm works by storing information on all *perimeter sites* in the matrix, `M`. Whether the four possible adjacent sites are occupied (1) or not (0) are stored (in the order right, below, left, above). A site is sampled from the perimeter sites and a new cell placed in it. A cell from adjacent occupied sites is sampled to be its mother, proportional to the mother’s fitness minus one. The new daughter’s fitness is updated as in the *cell-based* algorithm, and the cell’s information written to `M_final`. The mother’s fitness is updated using the same mutation rule. The daughter’s site is removed from `M`. When the daughter’s site is chosen, the occupancy of adjacent sites for perimeter sites changes, and this information is updated in `M`. Some sites adjacent to the daughter may become new perimeter sites and these are appended to `M`.

The colony growth models we use, and the associated mutation model, is likely to result in more efficient selection than would be seen in reality.

1/ *E. coli* colonies tend to have rougher edges than the Eden model predicts, and rougher edges generate greater drift (Hallatschek et al. 2007, Farrell.2017)

2/ We assume mutations occur at constant rate per replication. If mutations occurred at a constant rate per time, many would fall in cells not in the growing edge and are destined never to increase in frequency (Gralka et al. 2016).

3/ *Cell-based*: The chance of being a parent does not depend on how many empty sites you are adjacent to. In reality, cells with more empty adjacent sites are more likely to divide because they have more resources. Since high fitness genotypes should have, on average, fewer empty adjacent sites (because they are filled by their daughters) this will off-set some of their fitness advantage (captured in *Site-based*).

4/ *Cell-based*: The fitness of a cell is not dependent on the fitness of cells that share its free adjacent site. In reality, it is probably best to share free adjacent sites with low fitness genotypes since they will be less competitive. However, high fitness genotypes are more likely to share free adjacent sites with high fitness genotypes therefore reducing the strength of selection on them (captured in *Site-based*).

However, the colony is assumed 2-dimensional. Adding growth in a third dimension (i.e. perpendicular to the plate) is likely to result in less drift per `generation’ because the population of dividing cells is larger (the surface to volume ratio for a sphere is \(3/r\) where as the edge to area ratio for a circle is \(2/r\) - and for a flattened colony the discrepancy will be even greater). Off-setting this, however, is the fact that a given population size can be achieved in fewer generations (Gralka et al. (2016) and Section 9). In addition, it has been shown that beneficial mutations that occur during colony growth are those that have greater chance of moving to the colony edge (Kim et al. 2014) - a phenomenon not captured in our simulations.

#### 1.2 Well-mixed Growth

In addition to the colony growth simulations described above, we also implement three simulations for exploring well mixed growth in which the mutation rule is the same as in colony growth:

*Doubling*: In each generation the population doubles exactly. In generation \(t\), the parents for the \(2^{t}\) offspring are sampled with replacement from the \(2^{t-1}\) individuals present in generation \(t-1\). The probability a parent is chosen is proportional to its fitness minus one. Note that for parents, mutations are only simulated if they reproduce, and even then, it does not depend on how many offspring they have.

*Poisson*: In generation \(t\), the number of daughters produced by each parent is sampled from a Poisson distribution with a mean equal to their fitness minus one. In the absence of mutation and/or a DFE that has a point mass at zero, the population is *expected* to double each generation since the initial ancestor has a fitness of two. However, there will be random fluctuations in the population size, and, if beneficial mutations spread, the population is expected to more than double each generation. As with the Doubling scenario, only single mutation events are simulated for reproducing parents. This is the scenario simulated in Wahl and Agashe (2022).

*Single-step*: The *Doubling* and *Poisson* simulations have discrete generations. However, the colony simulations are in continuous time and cells are picked to be the next ones to divide. In the *Single-step* simulation the population grows by one each time-step and the parents of the single offspring at time \(t\) are sampled from the \(t-1\) parents proportional to their fitnesses minus one. In addition, single mutation events are simulated for each reproductive event and so a cell with two direct descendants has two opportunities for mutation, whereas in the Doubling and Poisson approaches there is only a single opportunity for mutation if a cell has two offspring in the *same* time-step. The *Single-step* approach will therefore result in more similar mutation rates as the colony-growth models, although this is unlikely to affect the distribution of mutations sampled. The *Single-step* approach will also result in faster adaptation compared to the Doubling and Poisson approaches because the number of generations is increased (Section 9).

#### 1.3 Spontaneous mutations versus segregating variation

Conflicting conclusions have been drawn from empirical work contrasting adaptation under under colony growth versus well-mixed growth: Gralka et al. (2016) suggest adaptation is faster under colony growth in *E coli* and *S cerevisiae* whereas Lavrentovich et al. (2016) suggest adaptation is slower in *S cerevisiae*. A number of differences between the studies exist that may explain their opposing conclusions. Gralka et al. (2016) look at how initially segregating beneficial mutations change in frequency per unit of population growth (rather than generations, which are not comparable under colony and well-mixed growth) whereas Lavrentovich et al. (2016) look at fixation probabilities of spontaneous deleterious mutations. In order to assess whether the discrepancy arises because segregating variants are compared against spontaneous mutations, the simulations also allow the scenario where mutations happen instantaneously to all cells present at a specified point in time. By setting the time point relatively early and the mutation rate artificially high we can mimic the scenario where a colony is produced from an initial well-mixed population of cells in which mutations are segregating. By making the DFE a point mass (i.e. making the scale zero) we can simulate a scenario close to that of Gralka et al. (2016) where a population with two variants is used to initiate a colony. This is only implemented for colony growth and single-step well-mixed growth.

### 2 Density function and random-number generator for the DFE.

We assume that the selection coefficients, \(s\), follow a shifted and reflected gamma with a mean of `means` and a `shape` and `scale` parameter. The function `rdfe` generates random numbers form this distribution:

```
rdfe<-function(n=1, means, shape, scale){

  # the mean of the gamma  = shape*scale and so the reflected gamma has mean -shape*scale
  # to translate to the new mean means we have to shift by means+shape*scale

  means+shape*scale-1*rgamma(n, shape=shape, scale=scale)

}
```

The function `ddfe` calculates the probability density of \(s\) (`x`) under this distribution, where the default `bound=TRUE` generates an error if `x` does not have support (i.e. if `x`>`shape*scale+means`) or gives a density of zero or a log-density of \(10^{-16}\) when `FALSE`. The `bound=FALSE` option is primarily implemented for maximum-likelihood estimated of the true DFE from the observed DFE where tested parameter values for the true DFE may generate a log-density of minus infinity.

```
ddfe<-function(x, means, shape, scale, log=FALSE, bound=TRUE){

  # the mean of the gamma  = shape*scale and so the reflected gamma has mean -shape*scale
  # to translate to the new mean means we have to shift by means+shape*scale

  if(log){
    d<-dgamma(-x+shape*scale+means, shape=shape, scale=scale, log=TRUE)
  }else{
    d<-dgamma(-x+shape*scale+means, shape=shape, scale=scale)
  }

  # Note that x cannot be greater than shape*scale+mean

  if(any(x>(shape*scale+means))){
    if(bound){
      stop("x cannot be greater than shape*scale+mean under this distribution")
    }else{
      if(log){
        d[which(x>(shape*scale+means))]<--10^15
      }
    }
  }
  return(d)  
}
```

### 3 Growth Simulation Functions

A suite of functions are implemented that simulate population growth with selection. All functions have the same arguments: `N` (the final population size, or the expected final population size under neutrality if the *Poisson* scenario is simulated), `mu` (the mutation rate) and `means`, `shape` and `scale` (the parameters of the DFE). In addition, for colony growth and single-step well-mixed growth, a population size can be specified in `seg.N`. When non-null, all cells present when the population size reaches `seg.N` are subject to mutation, but not at other times. A potential issue with the simulations is that if the number of mutants per cell is high and/or the chance of sampling strongly deleterious mutations from the DFE is high, then it is possible for \(w-1\) to be negative. This arises since the fitness of a cell with \(n\) mutations is

\[w=2\prod^n\_i(1+s\_i)\]

and so even a single mutation with \(s<-0.5\) would generate a negative value of \(w-1\). In the simulations of Wahl and Agashe (2022) the DFE is very wide with a substantial tail below \(-0.5\) yet this should give a negative expectation for the Poisson, which is invalid. It is not clear how this was dealt with in Wahl and Agashe (2022), but here we set the probability of having a daughter to zero in such cases.

#### 3.1 Colony growth

`EdenSimCell` simulates the process under *cell-based* colony growth:

```
EdenSimCell<-function(N, mu=0, means=0, shape=1, scale=1, seg.N=NULL){

  # Matrix for storing edge cell information.

  M<-matrix(0, 2, 10)
  colnames(M)<-c("w", "x", "y", "t", "d", "nmut", "adj1", "adj2", "adj3", "adj4")

  # w contains the fitness of the cell
  # x and y are the co-ordinates of the cell
  # t is the time-step at which the daughter arose
  # d is the number of cell divisions the cell has been through since the ancestor
  # The adj columns designate whether an adjacent site is occupied (1) or not (0)
  # For a cell at (x,y) the adj columns give the occupancy states at sites: 

  # adj1 = x+1, y 
  # adj2 = x, y-1
  # adj3 = x-1, y 
  # adj4 = x, y+1

  adj.col<-match(paste0("adj", 1:4), colnames(M))
  # position of adj1, adj2, adj3 or adj4 in M.

  # We can specify the four displacements as

  x.move<-c(1, 0, -1, 0)
  y.move<-c(0, -1, 0, 1)

  # If move from site A to B via a displacement adj1, adj2, adj3 or adj4,
  # which displacement would take us from site B back to A:

  back.move<-c(3,4,1,2)

  # For example, if we are at site A and move to site B via the adj2 displacement (0,-1) 
  # we need to make the displacement (0,1) to get from site B back to A.  
  # back.move[2]=4 and indeed adj4 = (0,1). 
 
  # An initial cell is placed at (0,0) with a daughter at (1,0)
  # A daughter is added so that M always has >1 row (otherwise M is coerced into a vector)

  M[1,"t"]<-1       # mother's time
  M[2,"t"]<-2       # daughters's time

  M[1,"d"]<-1       # mother divisions
  M[2,"d"]<-1       # daughters divisions


  if(is.null(seg.N)){
    spontaneous=FALSE
  }else{
    spontaneous=TRUE
  }

  if(runif(1)<mu & !spontaneous){  # 'mother's' fitness after division
    M[1,"w"]<-2*(1+rdfe(1, means=means, shape=shape, scale=scale))
    M[1,"nmut"]<-1  # 'mother's' mutational burden
  }else{
    M[1,"w"]<-2
  }  
 
  if(runif(1)<mu & !spontaneous){  # 'daughter's' fitness after division
    M[2,"w"]<-2*(1+rdfe(1, means=means, shape=shape, scale=scale))
    M[2,"nmut"]<-1  # 'daughter's' mutational burden
  }else{
    M[2,"w"]<-2     
  }

  M[1,"adj1"]<-1  # the mother's adjacent site is occupied by the daughter
  M[2,"x"]<-1     # daughter's x-coordinate is 1 (y-coordinate is 0)
  M[2,"adj3"]<-1  # the daughter's adjacent site is occupied by the mother

  # Matrix for storing cell information.

  M_final<-matrix(0, N, 6)
  colnames(M_final)<-c("w", "x", "y", "t", "d", "nmut")
      
  new_info<-rep(0,ncol(M)) # temporary vector for new daughter cell
  names(new_info)<-colnames(M)

  cnt<-1 # number of cells-1 currently copied to M_final.

  ##############
  ## Simulate ##
  ##############

  for(i in 3:N){

    parent<-sample(1:nrow(M), 1, prob=pmax(M[,"w"]-1,10^-16))
    # sample a parent 

    daughter<-sample(1:4, 1, prob=1-M[parent,adj.col])
    # sample one of the parent's free adjacent sites

    new.x<-M[parent,"x"]+x.move[daughter]
    new.y<-M[parent,"y"]+y.move[daughter]
    # co-ordinates of daughter cell

    M[parent,"d"]<-M[parent,"d"]+1
    # add a division event to the mother

    # put daughter's info for M/M_final in to new_info,

    if(runif(1)<mu & !spontaneous){
      new_info["w"]<-M[parent,"w"]*(1+rdfe(1, means=means, shape=shape, scale=scale))
      new_info["nmut"]<-M[parent,"nmut"]+1
    }else{  
      new_info["w"]<-M[parent,"w"]
      new_info["nmut"]<-M[parent,"nmut"]
    }

    new_info["x"]<-new.x
    new_info["y"]<-new.y
    new_info["t"]<-i
    new_info["d"]<-M[parent,"d"]

    if(runif(1)<mu & !spontaneous){  # add mutation to the `parent` (really another new daughter) 
      M[parent,"w"]<-M[parent,"w"]*(1+rdfe(1, means=means, shape=shape, scale=scale))
      M[parent,"nmut"]<-M[parent,"nmut"]+1
    }  
  

    for(j in 1:4){

       neighbour.x<-which(M[,"x"]==(new.x+x.move[j]))
       # find any cells that are in a column compatible
       # with an adj[j] displacement from the daughter
       neighbour.y<-match(new.y+y.move[j], M[neighbour.x,"y"])
       # find *the* neighbour.x cell (if any) that is in the row
       # of an adj[j] displacement from the daughter
       neighbour<-neighbour.x[neighbour.y]
       # the position of the cell (if any) in M
       # that is an adj[j] displacement from the daughter

       if(!is.na(neighbour)){ # if there is a neighbour at the adj[j] displacement

          new_info[6+j]<-1                # the daughter's adj[j] displacement is occupied
          M[neighbour,6+back.move[j]]<-1  # the neighbour's adj[back.move[j]] displacement
                                          # is occupied (by the daughter)

          if(sum(M[neighbour,adj.col])==4){   # if a neighbour has no free adjacent sites

            M_final[cnt,1:6]<-M[neighbour,1:6]  # write neighbour's info to M_final
            M<-M[-neighbour,]                   # delete neighbour from M
            cnt<-cnt+1
          
          }
       
       }else{

          new_info[6+j]<-0 # the daughter's adj[j] displacement is not occupied

       }   
    }

    if(sum(new_info[adj.col])==4){  # if the daughter has no free adjacent sites

      M_final[cnt,]<-new_info[1:6] # write daughter's info to M_final
      cnt<-cnt+1

    }else{
      M<-rbind(M, new_info)        # append daughter to M
    }

    if(spontaneous && i==seg.N){  # add mutations to all cells
        M[,"nmut"]<-rbinom(nrow(M), prob=mu, size=1)
        M_final[1:(cnt-1),"nmut"]<-rbinom(cnt-1, prob=mu, size=1)
        M[which(M[,"nmut"]==1),"w"]<-2*(1+rdfe(sum(M[,"nmut"]), means=means, shape=shape, scale=scale))
        M_final[which(M_final[1:(cnt-1),"nmut"]==1),"w"]<-2*(1+rdfe(sum(M_final[1:(cnt-1),"nmut"]), means=means, shape=shape, scale=scale))
    }  
  }

  M_final[cnt:N,]<-M[,1:6]  # write the final edges cells to M_final

  return(M_final)
}
```

`EdenSimSite` simulates the process under *site-based* colony growth:

```
EdenSimSite<-function(N, mu=0, means=0, shape=1, scale=1, seg.N=NULL){

  # Matrix for storing perimeter site information.

  M<-matrix(NA, 6, 6)
  colnames(M)<-c("x", "y", "adj1", "adj2", "adj3", "adj4")

  # The adj columns designate which cell is in an adjacent site where 0 indicate unoccupied
  # For a perimeter site at (x,y) the adj columns give the cell at sites 

  # adj1 = x+1, y right
  # adj2 = x, y-1 below
  # adj3 = x-1, y left
  # adj4 = x, y+1 above

  adj.col<-match(paste0("adj", 1:4), colnames(M))
  # position of adj1, adj2, adj3 or adj4 in M.

  # We can specify the four displacements as

  x.move<-c(1, 0, -1, 0)
  y.move<-c(0, -1, 0, 1)

  # If move from site A to B via a displacement adj1, adj2, adj3 or adj4,
  # which displacement would take us from site B back to A:

  back.move<-c(3,4,1,2)

  # For example, if we are at site A and move to site B via the adj2 displacement (0,-1) 
  # we need to make the displacement (0,1) to get from site B back to A.  
  # back.move[2]=4 and indeed adj4 = (0,1). 
 
  # Matrix for storing cell information.

  M_final<-matrix(0, N, 6)
  colnames(M_final)<-c("w", "x", "y", "t", "d", "nmut")

  # An initial cell is placed at (0,0) with a daughter at (1,0)
  # A daughter is added so that M always has >1 row (otherwise M is coerced into a vector)

  M_final[1,"x"]<-0       # mother's coordinates are (0,0)
  M_final[1,"y"]<-0

  M_final[2,"x"]<-1       # daughter's coordinates are (1,0)
  M_final[2,"y"]<-0

  M_final[1,"t"]<-1       # mother's time
  M_final[2,"t"]<-2       # daughter's time

  M_final[1,"d"]<-1       # mother's divisions
  M_final[2,"d"]<-1       # daughter's divisions

  M_final[1,"w"]<-2       # mother's fitness
  M_final[1,"nmut"]<-0    # mother's mutational 'burden'

  if(is.null(seg.N)){
    spontaneous=FALSE
  }else{
    spontaneous=TRUE
  }

  if(runif(1)<mu & !spontaneous){  # 'mother's' fitness after cell division
    M_final[1,"w"]<-2*(1+rdfe(1, means=means, shape=shape, scale=scale))
    M_final[1,"nmut"]<-2  # mother's mutational 'burden'
  }else{
    M_final[1,"w"]<-2   
  }

  if(runif(1)<mu & !spontaneous){ # 'daughter's' fitness after cell division
    M_final[2,"w"]<-2*(1+rdfe(1, means=means, shape=shape, scale=scale)) 
    M_final[2,"nmut"]<-1  # daughter's mutational 'burden'
  }else{
    M_final[2,"w"]<-2  
  }

  # fill in information for 6 initial perimeter sites

  M[,"x"]<-c(-1, 0, 0, 1, 1, 2)
  M[,"y"]<-c(0, 1, -1, 1, -1, 0)
  M[1,"adj1"]<-M[2,"adj2"]<-M[3,"adj4"]<-1
  M[4,"adj2"]<-M[5,"adj4"]<-M[6,"adj3"]<-2

  new_info<-rep(NA,nrow(M)) # temporary vector for new perimeter site
  names(new_info)<-colnames(M)

  ##############
  ## Simulate ##
  ##############

  for(i in 3:N){

    daughter<-sample(1:nrow(M), 1)
    # sample a daughter site

    possible_parents<-M[daughter, adj.col]

    parental_fitness<-pmax(M_final[possible_parents,"w"]-1,10^-16)
    parental_fitness[which(is.na(parental_fitness))]<-10^-16

    parent<-possible_parents[sample(1:4, 1, prob=parental_fitness)]
    # sample a parent from the daughter's occupied adjacent sites

    M_final[parent,"d"]<-M_final[parent,"d"]+1 # add a parental division

    new.x<-M[daughter, "x"]
    new.y<-M[daughter, "y"]

    # put daughter's info for M_final,

    if(runif(1)<mu  & !spontaneous){
      M_final[i,"w"]<-M_final[parent,"w"]*(1+rdfe(1, means=means, shape=shape, scale=scale))
      M_final[i,"nmut"]<-M_final[parent,"nmut"]+1
    }else{  
      M_final[i,"w"]<-M_final[parent,"w"]
      M_final[i,"nmut"]<-M_final[parent,"nmut"]
    }   
    M_final[i,"x"]<-new.x
    M_final[i,"y"]<-new.y
    M_final[i,"t"]<-i
    M_final[i,"d"]<-M_final[parent,"d"]

    if(runif(1)<mu  & !spontaneous){  # add mutation to the `parent` (really another new daughter) 
      M_final[parent,"w"]<-M_final[parent,"w"]*(1+rdfe(1, means=means, shape=shape, scale=scale))
      M_final[parent,"nmut"]<-M_final[parent,"nmut"]+1   
    }  

    M<-M[-daughter,]

    for(j in 1:4){

       neighbour.x<-which(M[,"x"]==(new.x+x.move[j]))
       # find any cells that are in a column compatible
       # with an adj[j] displacement from the daughter
       neighbour.y<-match(new.y+y.move[j], M[neighbour.x,"y"])
       # find *the* neighbour.x cell (if any) that is in the row
       # of an adj[j] displacement from the daughter
       neighbour<-neighbour.x[neighbour.y]
       # the position of the cell (if any) in M
       # that is an adj[j] displacement from the daughter

       if(is.na(neighbour)){  # if there is no neighbour at the adj[j] displacement

         neighbour.x<-which(M_final[,"x"]==(new.x+x.move[j]))
         # find any cells that are in a column compatible
         # with an adj[j] displacement from the daughter
         neighbour.y<-match(new.y+y.move[j], M_final[neighbour.x,"y"])
      
         if(is.na(neighbour.y)){ # and there's no cell at the adj[j] displacement

            new_info[adj.col]<-NA
            new_info["x"]<-new.x+x.move[j] 
            new_info["y"]<-new.y+y.move[j]
            new_info[2+back.move[j]]<-i
            
            M<-rbind(M, new_info)
         }   

       }else{  # if there is a neighbour at the adj[j] displacement update it's adj1 

          M[neighbour,2+back.move[j]]<-i  # the neighbour's adj[back.move[j]] displacement
                                          # is occupied (by cell i, the daughter)
       }   

    }

    if(spontaneous && i==seg.N){  # add mutations to all cells
        M_final[,"nmut"]<-rbinom(nrow(M_final), prob=mu, size=1)
        M_final[which(M_final[,"nmut"]==1),"w"]<-2*(1+rdfe(sum(M_final[,"nmut"]), means=means, shape=shape, scale=scale))
    }  
  }

  return(M_final)
}
```

#### 3.2 Well-mixed growth

`EdenSimDoub` simulates the process under *doubling* well-mixed growth:

```
EdenSimDoub<-function(N, mu=0, means=0, shape=1, scale=1){

  # Matrix for storing cell information.

  M_final<-matrix(0, N, 4)
  colnames(M_final)<-c("w", "t", "d", "nmut")
  M_final[1,]<-c(2,0,0,0)

  tau<-ceiling(log(N)/log(2))

  ##############
  ## Simulate ##
  ##############

  for(i in 1:tau){

    nparents<-2^(i-1)

    if(i==tau){
      noffspring<-(N-2^(tau-1))
    }else{
      noffspring<-2^(i-1)
    }

    parents<-sample(1:nparents, noffspring, prob=pmax(M_final[1:nparents,"w"]-1,10^-16), replace=TRUE)
    # sample a parent 

    mut_event<-rbinom(noffspring, prob=mu, size=1)
    s<-rdfe(noffspring, means=means, shape=shape, scale=scale)

    M_final[unique(parents),"d"]<-M_final[unique(parents),"d"]+1 # note here we are counting a single division even if the `mother` had multiple offspring 

    M_final[nparents+1:noffspring,"w"]<-M_final[parents,"w"]*(1+mut_event*s)
    M_final[nparents+1:noffspring,"nmut"]<-M_final[parents,"nmut"]+mut_event
    M_final[nparents+1:noffspring,"t"]<-i
    M_final[nparents+1:noffspring,"d"]<-M_final[parents,"d"]
    # update 'daughter's' fitness

    # update 'mother's' fitness

    mut_event<-(rbinom(nparents, prob=mu, size=1)==1 & (1:nparents)%in%parents)
    s<-rdfe(nparents, means=means, shape=shape, scale=scale)

    M_final[1:nparents,"w"]<-M_final[1:nparents,"w"]*(1+mut_event*s)
    M_final[1:nparents,"nmut"]<-M_final[1:nparents,"nmut"]+mut_event

  }
  return(M_final)
}
```

`EdenSimPois` simulates the process under *Poisson* well-mixed growth:

```
EdenSimPois<-function(N, mu=0, means=0, shape=1, scale=1){

  tau<-log(N)/log(2)

  if(round(tau,4)%%1!=0){
    stop("N should be such that tau is integer: i.e. N  = exp(tau*log(2)) where tau is integer")
  }else{
    tau<-round(tau)
  }

  # Matrix for storing cell information (twice as large as it needs to be).

  M_final<-matrix(0, 1, 4)
  colnames(M_final)<-c("w", "t", "d", "nmut")
  M_final[1,]<-c(2,0,0, 0)


  ##############
  ## Simulate ##
  ##############

  nparents<-1

  for(i in 1:tau){

    N_off<-rpois(nparents, lambda=pmax(M_final[1:nparents,"w"]-1,0))
    # number of offspring per daughter

    noffspring<-sum(N_off)

    if(noffspring>0){
      # update 'daughter's' fitness

      mut_event<-rbinom(noffspring, prob=mu, size=1)
      s<-rdfe(noffspring, means=means, shape=shape, scale=scale)

      parents<-rep(1:nparents, N_off)

      M_final[unique(parents),"d"]<-M_final[unique(parents),"d"]+1

      M_offspring<-cbind(
        M_final[parents,"w"]*(1+mut_event*s),
        i,
        M_final[parents,"d"]+1,
        M_final[parents,"nmut"]+mut_event
      )

      # update 'mother's' fitness

      mut_event<-(rbinom(nparents, prob=mu, size=1)==1 & (1:nparents)%in%parents)
   
      s<-rdfe(nparents, means=means, shape=shape, scale=scale)

      M_final[,"w"]<-M_final[,"w"]*(1+mut_event*s)
      M_final[,"nmut"]<-M_final[,"nmut"]+mut_event

      M_final<-rbind(M_final, M_offspring)
    }
      
    nparents<-nparents+noffspring
  }
  return(M_final)
}
```

`EdenSimSing` simulates the process under *single-step* well-mixed growth:

```
EdenSimSing<-function(N, mu=0, means=0, shape=1, scale=1, seg.N=NULL){

  # Matrix for storing cell information.

  M_final<-matrix(0, N, 4)
  colnames(M_final)<-c("w", "t", "d","nmut")

  M_final[1,"t"]<-1       # mother's time
  M_final[2,"t"]<-2       # daughter's time

  M_final[1,"d"]<-1       # mother's divisions
  M_final[2,"d"]<-1       # daughter's divisions

  M_final[1,"w"]<-2       # mother's fitness
  M_final[1,"nmut"]<-0    # mother's mutational 'burden'

  if(is.null(seg.N)){
    spontaneous=FALSE
  }else{
    spontaneous=TRUE
  }

  if(runif(1)<mu & !spontaneous){  # 'mother's' fitness after cell division
    M_final[1,"w"]<-2*(1+rdfe(1, means=means, shape=shape, scale=scale))
    M_final[1,"nmut"]<-1  # mother's mutational 'burden'
  }else{
    M_final[1,"w"]<-2   
  }

  if(runif(1)<mu & !spontaneous){ # 'daughter's' fitness after cell division
    M_final[2,"w"]<-2*(1+rdfe(1, means=means, shape=shape, scale=scale))
    M_final[2,"nmut"]<-1  # daughter's mutational 'burden'
  }else{
    M_final[2,"w"]<-2  
  }


  ##############
  ## Simulate ##
  ##############

  for(i in 3:N){

    parent<-sample(1:(i-1), 1, prob=pmax(M_final[1:(i-1),"w"]-1,10^-16))
    # sample a parent 

    M_final[parent,"d"]<-M_final[parent,"d"]+1

    # update 'daughter's' fitness

    if(runif(1)<mu  & !spontaneous){
      M_final[i,"w"]<-M_final[parent,"w"]*(1+rdfe(1, means=means, shape=shape, scale=scale))
      M_final[i,"nmut"]<-M_final[parent,"nmut"]+1
    }else{  
      M_final[i,"w"]<-M_final[parent,"w"]
      M_final[i,"nmut"]<-M_final[parent,"nmut"]
    }
    M_final[i,"t"]<-i
    M_final[i,"d"]<-M_final[parent,"d"]

    # update 'mother's' fitness

    if(runif(1)<mu  & !spontaneous){
      M_final[parent,"w"]<-M_final[parent,"w"]*(1+rdfe(1, means=means, shape=shape, scale=scale))
      M_final[parent,"nmut"]<-M_final[parent,"nmut"]+1
    }

    if(spontaneous && i==seg.N){  # add mutations to all cells
      M_final[,"nmut"]<-rbinom(nrow(M_final), prob=mu, size=1)
      M_final[which(M_final[,"nmut"]==1),"w"]<-2*(1+rdfe(sum(M_final[,"nmut"]), means=means, shape=shape, scale=scale))
    }  
  }
  return(M_final)
}
```

### 4 Mutation-accumulation Simulation Function

In typical microbial MA experiments a single cell is taken from a colony and used to initiate a new colony where the procedure is repeated. In general, this procedure is repeated for many transfers such that cells at the end of the MA experiment have accumulated many mutations. However, the method for correcting for selection bias in Wahl and Agashe (2022) is only applicable to DFE’s estimated from single mutations (see Section 5). The `simulate.MA` function simulates `nlines` MA lines for one round of growth and selects all single mutants cells (`sample_all=TRUE`) or one single mutant cell (`sample_all=FALSE`) if present. Real MA experiments do not have the luxury of selecting single-mutants to be the founder of the next colony, and most transfers will involve non-mutant cells. Real MA experiments, therefore, have to be run for much longer than *in-silico* MA experiments in order to accumulate single-mutants. We refer to the distribution of selection coefficients obtained in this manner as the distribution of sampled fitness effects (DSFE) which will deviate from the DFE due to selection during the MA process. If `sample_all=TRUE` then more single-mutants can be sampled, but the accumulated mutations will not be independent and so `sample_all=FALSE` is preferable when making statistical inferences. As in the simulation functions described above, `simulate.MA` also takes the arguments `N`, `mu`, `means`, `shape` and `scale` together with the argument `method` for specifying which type of population growth is to be simulated (`"cell"`, `"site"`, `"doub"`, `"pois"`, `"sing"`).

```
simulate.MA<-function(nlines=1, N, mu, means=0, shape=1, scale=1, sample_all=FALSE, method="doub", verbose=FALSE){

  if(!method%in%c("cell", "site", "doub", "pois", "sing")){stop("method must be one of 'cell', 'site', 'doub', 'pois' or 'sing'")}

  edfe<-c()
  ngen<-1:nlines

  cnt<-0

  for(i in 1:nlines){

    if(method=="cell"){
      res<-EdenSimCell(N=N, mu=mu, means=means, shape=shape, scale=scale)
    }
    if(method=="site"){
      res<-EdenSimSite(N=N, mu=mu, means=means, shape=shape, scale=scale)
    }
    if(method=="doub"){
      res<-EdenSimDoub(N=N, mu=mu, means=means, shape=shape, scale=scale)
    }
    if(method=="pois"){
      res<-EdenSimPois(N=N, mu=mu, means=means, shape=shape, scale=scale)
    }
    if(method=="sing"){
      res<-EdenSimSing(N=N, mu=mu, means=means, shape=shape, scale=scale)
    }

    ngen[i]<-mean(res[,"d"])

    single_mu<-which(res[,"nmut"]==1)

    nsingle_mu<-length(single_mu)

    if(nsingle_mu!=0){
      if(!sample_all & nsingle_mu>1){
        single_mu<-sample(single_mu, 1)
      }

      edfe<-c(edfe,res[single_mu,"w"]/2-1)
      
      if(verbose){
        print(i)
      }
    }    
  }

  return(list(edfe=edfe, ngen=ngen))
}
```

### 5 Functions for assessing the magnitude and impact of selection bias

Wahl and Agashe (2022) (Equation 2) give an expression for selection bias (the factor by which mutations with selection coefficient \(s\) are more or less likely to be sampled at the end of the growth phase compared to neutral mutations - i.e. an odds ratio):

\[b(s | \tau) = \frac{2^{s\tau}-1}{s\tau ln(2)}\]

where \(\tau\) is the time at the end of the growth phase where time is measured in units of population doublings. Assuming no death, \(N = 2^\tau\), and so \(ln(N)/ln(2) = \tau\) to give

\[b(s | \tau) = \frac{2^{sln(N)/ln(2)}-1}{sln(N)}\]

The function `wahl.bias` calculates this bias given `s` and `tau`:

```
wahl.bias<-function(s, N=NULL, tau=NULL, log=FALSE){

  if(is.null(tau)){
    if(is.null(N)){stop("either N or tau must be given")}
    tau<-log(N)/log(2)
  }else{
    if(!is.null(N)){stop("either N or tau must be given, not both")}
  }

  bs<-(2^(s*tau)-1)/(s*tau*log(2))
  if(any(abs(s)<=.Machine$double.eps)){
    bs[which(abs(s)<=.Machine$double.eps)]<-1
  }  
  if(log){
    log(bs)
  }else{
    bs
  }
}
```

If we multiply the correction by the true DFE and normalise it we get the predicted probability density of the *observed* DFE, which we refer to as the DSFE\(\_{\tau}\) to indicate this is the estimated DSFE given the known \(\tau\). This is implemented in the function `odfe` which requires the parameters of the true DFE (`means`, `shape` and `scale`) and \(\tau\) (`tau`) which determines the amount of selection bias.

```
odfe<-function(x, means, shape, scale, N=NULL, tau=NULL, log=FALSE, normalise=TRUE, bound=TRUE){

    if(is.null(tau)){
      if(is.null(N)){stop("either N or tau must be given")}
    }else{
      if(!is.null(N)){stop("either N or tau must be given, not both")}
    }

    f<-function(x){ddfe(x, means=means, shape=shape, scale=scale, bound=bound)*wahl.bias(x,N=N, tau=tau)}

    if(normalise){
      C<-integrate(f, -1, shape*scale+means)$value
    }else{
      C<-1
    }

    if(log){
       ddfe(x, means=means, shape=shape, scale=scale, log=TRUE, bound=bound)+wahl.bias(x, N=N, tau=tau, log=TRUE)-log(C)
    }else{
       f(x)/C
    }
}
```

The function `fit.wahl` takes samples from the observed DFE (DSFE) and the parameters of the true DFE and estimates the number of population doublings that is consistent with the discrepancy between the observed and actual DFE, given the Wahl and Agashe (2022) model for selection bias. We refer to this estimated \(\tau\) as the effective \(\tau\) or \(\tau\_e\). The log-likelihood and standard error are also given. If an alternative \(\tau\) is given in the argument `true` a likelihood ratio test is conducted assuming \(\tau\)=`true` to be the null hypothesis.

```
fit.wahl<-function(x, means, shape, scale, initial=NULL, true=NULL, tau=FALSE){

  if(tau){
    if(is.null(initial)){initial<-7}
    tau.est<-function(tau, x, means, shape, scale){
       -sum(odfe(x, means, shape, scale, tau=tau, log=TRUE))
    }
    res<-optim(initial, tau.est, x=x, means=means, shape=shape, scale=scale, method="Brent", lower=0, upper=100, hessian=TRUE)

    res$logL<--res$value

    if(!is.null(true)){
      res$p.value<--(res$value-tau.est(true, x, means, shape, scale))
      res$p.value<-1-pchisq(2*res$p.value, 1)
    }
  }else{
    if(is.null(initial)){initial<-100}
    N.est<-function(N, x, means, shape, scale){
    }
    
    res<-optim(initial, N.est, x=x, means=means, shape=shape, scale=scale, method="Brent", lower=0, upper=10^16, hessian=TRUE)

    res$logL<--res$value

    if(!is.null(true)){ # likelihood ratio test against the true value
       res$p.value<--(res$value-N.est(true, x, means, shape, scale))
       res$p.value<-1-pchisq(2*res$p.value, 1)
    }     
  }
    
  if(!is.null(true)){  
    return(list(est=res$par, se=sqrt(1/res$hessian), logL=res$logL, p.value=res$p.value))
  }else{
    return(list(est=res$par, se=sqrt(1/res$hessian), logL=res$logL))
  }  

}
```

Wahl and Agashe (2022) take the DSFE and correct it to get a better estimate of the true DFE. Their procedure starts by discretising the DSFE and then revising the probability that a mutation arises in each bin. However, we can also obtain an estimate of the true DFE conditional on \(\tau\) (DFE\(\_{\tau}\)) using maximum likelihood:

```
est.dfe<-function(x, N=NULL, tau=NULL, initial=c(-0.2,10,0.05), bound=TRUE){

    if(is.null(tau)){
      if(is.null(N)){stop("either N or tau must be given")}
    }else{
      if(!is.null(N)){stop("either N or tau must be given, not both")}
    }

    # Since x cannot be greater than shape*scale+means under the mean-shifted reflected gamma, this places constraints  

    f.log<-function(par,x, N=NULL, tau=NULL){
      -sum(odfe(x, means=par[1], shape=par[2], scale=exp(par[3]), N=N, tau=tau, log=TRUE, bound=bound))
    }

    initial.log<-initial
    initial.log[2]<-initial[2]
    initial.log[3]<-log(initial[3])

    res<-optim(initial, f.log, x=x, N=N, tau=tau, hessian=TRUE, lower=c(-1, 1e-2, -Inf), upper=c(1,100,100), method="L-BFGS-B")

    res<-list(est=cbind(res$par, sqrt(diag(solve(res$hessian)))), logL=-res$value, convergence=res$convergence==0)
    rownames(res$est)<-c("means", "shape", "log.scale")
    colnames(res$est)<-c("est", "se")
    res$est<-rbind(res$est, scale=c(exp(res$est["log.scale",1]), NA))
    return(res)
}
```

However, maximising the likelihood under the original parameterisation proved temperamental and so we estimated the `scale` parameter on the log scale: a maximum-likelihood estimate of the `scale` parameter on the original scale is still available, although standard errors are not. In addition, the mean-shifted reflected gamma distribution only has support for values less than `shape*scale+means` and so the maximum observed \(s\) puts constraints on `shape*scale+means` in addition to the positivity constraints on `shape` and `scale`. However, this additional constraint is ignored during optimisation.

The function `plot.wahl` takes samples from the observed DFE, such as the single-mutant fitness effects saved from the MA simulations, and plots a histogram. The probability density function of the true DFE, given through the parameters `means`, `shape`, `scale`, is overlaid as a solid black line and, if `est_dfe=TRUE`, the predicted observed DFE given `tau` (DSFE\(\_{\tau}\)) is overlaid as a red line. In addition, the probability density function of the DFE predicted using the Wahl and Agashe (2022) correction, given `tau`, is overlaid as a black dotted line (DFE\(\_{\tau}\)). If `est_bias=TRUE` then an estimate of \(\tau\) (\(\widehat{\tau\_e}\)) is made based on comparing the discrepancy between the true and observed DFE and a new predicted observed DFE based on the estimate of \(\tau\_e\) (DFSE\(\_{\tau\_e}\)) is overlaid in blue.

```
plot.wahl<-function(x, means, shape, scale, N=NULL, tau=NULL, est_bias=TRUE, est_dfe=TRUE, ...){

  x.pos<-seq(1.1*min(x), 1.1*max(x), length=100)

  heights<-hist(x, freq=FALSE, breaks=100, xlab="Fitness effects, s", main="", ...)

  act.dfe<-ddfe(x.pos, means=means, shape=shape, scale=scale, bound=FALSE)
  exp.dfe<-odfe(x.pos, means=means, shape=shape, scale=scale, N=N, tau=tau, bound=FALSE) 

  lines(exp.dfe~x.pos, col="red")
  lines(act.dfe~x.pos, col="black")

  if(est_dfe){
    correct.dfe.par<-est.dfe(x, tau=tau, N=N, initial=c(means, shape, scale), bound=FALSE)
    correct.dfe<-ddfe(x.pos, means=correct.dfe.par$est["means","est"], shape=correct.dfe.par$est["shape","est"], scale=correct.dfe.par$est["scale","est"], bound=FALSE)
    lines(correct.dfe~x.pos, col="black", lty=3)
  }

  if(est_bias){
    if(!is.null(tau)){

      est.tau<-fit.wahl(x, means=means, shape=shape, scale=scale, tau=TRUE)$est
      est.dfe<-odfe(x.pos, means=means, shape=shape, scale=scale, tau=est.tau, bound=FALSE)

    }else{

      est.N<-fit.wahl(x, means=means, shape=shape, scale=scale, tau=FALSE)$est
      est.dfe<-odfe(x.pos, means=means, shape=shape, scale=scale, N=est.N, bound=FALSE)

    }
    lines(est.dfe~x.pos, col="blue")
  }else{
    est.tau<-est.N<-0
  }

 plotted<-c(1:4)[which(c(TRUE, est_dfe, TRUE, est_bias))]

  if(!is.null(tau)){
    legend("topleft", legend=c("DFE", substitute(DFE[tau] ~ "     " ~ tau == tau.n, list(tau.n = round(tau, 3))), substitute( DSFE[tau] ~ "   " ~ tau == tau.n, list(tau.n = round(tau, 3))), substitute(DSFE[tau[e]] ~ " " ~ widehat(tau[e]) == tau.n, list(tau.n = round(est.tau, 3))))[plotted], col=c("black", "black", "red", "blue")[plotted], lty=c(1,3,1,1)[plotted], lwd=2, inset=0.05)

  }else{

    legend("topleft", legend=c("DFE", substitute(DFE[N] ~ "     " ~ N == N.n, list(N.n = round(N, 3))), substitute(DSFE[N] ~ "   " ~ N == N.n, list(N.n = round(N, 3))), substitute(DSFE[N[e]] ~ " " ~ widehat(N[e]) == N.n, list(N.n = round(est.N, 3))))[plotted], col=c("black", "black", "red", "blue")[plotted], lty=c(1,3,1,1)[plotted], lwd=2, inset=0.05)
  }  

}
```

Note that all functions have arguments `tau` and `N` and so either parameterisation (number of doublings or final population size) can be used by specifying one argument and leaving the other as `NULL`.

### 6 Simulations

In the following simulations we use the DFE used in Wahl and Agashe (2022) with a mean \(s\) of \(-0.2\), a shape of \(10\) and scale of \(0.05\). This DFE is much wider than expected for spontaneous mutations and we refer to it as the wide-DFE:

```
means<--0.2
shape<-10
scale<-0.05

x.pos<-seq(-1, 0.2, length=100)
plot(ddfe(x.pos,means=means, shape=shape, scale=scale)~x.pos, xlab="Distribution of Fitness Effects", ylab="Probability Density", type="l")
```

For example, in the *E.coli* experiments published in Couce et al. (2024), the bulk of the DFE typically lies between -0.05 and 0.02, despite the mutations being due to transposon insertions, which are likely to have larger fitness effects than spontaneous mutations. We set the mutation rate to \(10^{-3}\) as this is approximately the genomic mutation rate in *E. coli* (Foster et al. 2015). Since the simulations are rather slow, we only grow the colonies for 16 doublings (\(N=\) 65536 cells) - less than the 28 doublings (\(N=2.78\times10^{8}\) cells) typically seen in an *E. coli* colony after 24 hours of growth (Foster et al. 2015). The simulations are performed for 2,000 lines which generally result in close to 2,000 independent single-mutations (nearly all colonies have at least one single-mutant cell).

```
mu<-10^-3
nlines<-2000
tau<-16

N<-round(exp(tau*log(2)))
```

#### 6.1 Colony growth simulations

##### 6.1.1 Cell-based simulations

We simulate an MA experiment under *Cell-based* colony growth

```
edfe_cell<-simulate.MA(nlines=nlines, N=N, mu=mu, means=means, shape=shape, scale=scale, method="cell")$edfe
```

and we compare the simulated observed DFE with that predicted using the correction in Wahl and Agashe (2022):

```
plot.wahl(edfe_cell, means=means, shape=shape, scale=scale, tau=tau, est_bias=FALSE, est_dfe=FALSE)
```

We can see that selection bias under *cell-based* colony growth is actually much more pronounced than predicted using approximations under well-mixed growth (Wahl and Agashe 2022). The histogram of sampled selection coefficients is shifted toward more positive values compared to what we expect (DFSE\(\_{\tau}\) in red). Despite the increased drift variance expected under colony growth (Hallatschek et al. 2007) the increased number of generations required to achieve the same population size seems to result in greater levels of adaptation (Gralka et al. 2016). Consequently, if we try to estimate the true DFE assuming homogeneous growth selection bias (DFE\(\_{\tau}\) - black dotted line) we can see it is biased towards less deleterious or beneficial effects:

```
plot.wahl(edfe_cell, means=means, shape=shape, scale=scale, tau=tau, est_bias=FALSE)
```

If we try to estimate \(\tau\) the estimate is significantly greater than the true value (\(\tau\)=16)

```
fit.wahl(edfe_cell, means=means, shape=shape, scale=scale, true=tau, tau=TRUE)
```

```
## $est
## [1] 36.87703
## 
## $se
##           [,1]
## [1,] 0.6299973
## 
## $logL
## [1] 1604.863
## 
## $p.value
## [1] 0
```

This implies that selection bias in colonies is equivalent to that in a homogenous culture ran for almost twice as long (36.9 doublings as opposed to 16 doublings, assuming that the bias predicted by Wahl and Agashe (2022) is accurate under homogeneous growth (see below)). The observed DFE predicted using the effective \(\tau\) (DSFE\(\_{\tau\_e}\) in blue) seems to fit samples from the DFSE (the histogram) much better.

```
plot.wahl(edfe_cell, means=means, shape=shape, scale=scale, tau=tau)
```

##### 6.1.2 Site-based simulations

We can also use a site-based algorithm:

```
edfe_site<-simulate.MA(nlines=nlines, N=N, mu=mu, means=means, shape=shape, scale=scale, method="site")$edfe
```

and perform the same comparisons as was done for the cell-based simulations.

```
plot.wahl(edfe_site, means=means, shape=shape, scale=scale, tau=tau)
```

Here we see a dramatic difference - the amount of selection bias is considerably less than predicted in homogenous culture, equivalent to approximately 5.3 doublings (as opposed to 16) and correcting for the predicted selection bias results in an estimated DFE (dotted black line) that is considerably more deleterious than the true DFE (solid black line). This is presumably due to the high levels of kin-competition seen in a site-based scenario.

#### 6.2 Well-mixed growth simulations

We also ran a series of well-mixed growth simulations, including an implementation of the original simulation in Wahl and Agashe (2022) (*Poisson*). The two additional simulations differ from the *Poisson* simulation in that the final population size is constant (*Doubling* and *Singe-step*) rather than stochastic (*Poisson*) and growth is continuous (*Singe-step*) rather than in discrete generations (*Poisson* and *Doubling*). The colony growth models have fixed population sizes and are in continuous time and so the motivation for these additional well-mixed growth simulations is to make sure the differences between the colony growth simulations and the *Poisson* simulation are not due to these differences.

##### 6.2.1 Poisson simulations

We run the *Poisson* simulation of Wahl and Agashe (2022):

```
edfe_pois<-simulate.MA(nlines=nlines, N=N, mu=mu, means=means, shape=shape, scale=scale, method="pois")$edfe
```

and compare it to expectation

```
plot.wahl(edfe_pois, means=means, shape=shape, scale=scale, tau=tau)
```

We can see that the amount of selection bias is actually less than is predicted by Wahl and Agashe (2022), and is consistent with 9.7 doublings rather than (and significantly different from) 16. However, the estimated true DFE is not too dissimilar from the true DFE, suggesting that the over-estimate of the amount of selection bias in Wahl and Agashe (2022) has minimal substantive effect.

##### 6.2.2 Doubling simulations

Running the *Doubling* simulations

```
edfe_doub<-simulate.MA(nlines=nlines, N=N, mu=mu, means=means, shape=shape, scale=scale, method="doub")$edfe
```

and comparing

```
plot.wahl(edfe_doub, means=means, shape=shape, scale=scale, tau=tau)
```

we get very similar results to that seen under *Doubling*.

##### 6.2.3 Singe-step simulations

Running the *Singe-step* simulations

```
edfe_sing<-simulate.MA(nlines=nlines, N=N, mu=mu, means=means, shape=shape, scale=scale, method="sing")$edfe
```

and comparing

```
plot.wahl(edfe_sing, means=means, shape=shape, scale=scale, tau=tau)
```

we can see that the amount of selection bias is greater than that seen in the discrete generation simulations. This is assumed to happen because in the continuous time single-step simulations the number of generations exceeds \(\tau\) (see Section 9). The predicted selection bias is closer to that predicted in Wahl and Agashe (2022) but there does remain a small, but significant, over-estimation, presumably because the theory is deterministic and therefore does not account for the added drift. Nevertheless, the difference between the true and estimated DFE is very small.

### 7 Weak Selection

The DFE used in Wahl and Agashe (2022) generates mutations with unusually large fitness effects. A more reasonable DFE would be one with a mean selection coefficient of \(-0.02\), a shape of \(5\) and a scale of \(0.01\) (e.g. Couce et al. (2024)).

```
means_weak<--0.02
shape_weak<-5
scale_weak<-0.01
```

Even with a typical colony size achieved after 28 doublings (\(N=2.68\times 10^8\) cells) the amount of selection bias predicted under homogeneous growth is quite negligible since most mutations fall into the nearly neutral category.

```
x<-seq(-0.1,0.03,length=100)
act.dfe<-ddfe(x, means=means_weak, shape=shape_weak, scale=scale_weak)
exp.dfe<-odfe(x, means=means_weak, shape=shape_weak, scale=scale_weak, tau=28)

plot(act.dfe~x, type="l", ylim=range(c(act.dfe, exp.dfe)), ylab="Probability Density", xlab="Fitness effects, s")
lines(exp.dfe~x, col="red")
legend("topleft", legend=c("DFE", substitute( DSFE[tau] ~ "   " ~ tau == tau.n, list(tau.n = 28))), col=c("black", "red"), lty=c(1,1), lwd=2, inset=0.05)
```

With 16 doublings, as used in the simulations, selection bias would have been hard to detect:

```
act.dfe<-ddfe(x, means=means_weak, shape=shape_weak, scale=scale_weak)
exp.dfe<-odfe(x, means=means_weak, shape=shape_weak, scale=scale_weak, tau=tau)
plot(act.dfe~x, type="l", ylim=range(c(act.dfe, exp.dfe)), ylab="Probability Density", xlab="Fitness effects, s")
lines(exp.dfe~x, col="red")
legend("topleft", legend=c("DFE", substitute( DSFE[tau] ~ "   " ~ tau == tau.n, list(tau.n = 16))), col=c("black", "red"), lty=c(1,1), lwd=2, inset=0.05)
```

We can rerun the simulations with this more reasonable DFE:

```
edfe_cell_weak<-simulate.MA(nlines=nlines, N=N, mu=mu, means=means_weak, shape=shape_weak, scale=scale_weak, method="cell")$edfe
edfe_site_weak<-simulate.MA(nlines=nlines, N=N, mu=mu, means=means_weak, shape=shape_weak, scale=scale_weak, method="site")$edfe
edfe_doub_weak<-simulate.MA(nlines=nlines, N=N, mu=mu, means=means_weak, shape=shape_weak, scale=scale_weak, method="doub")$edfe
edfe_pois_weak<-simulate.MA(nlines=nlines, N=N, mu=mu, means=means_weak, shape=shape_weak, scale=scale_weak, method="pois")$edfe
edfe_sing_weak<-simulate.MA(nlines=nlines, N=N, mu=mu, means=means_weak, shape=shape_weak, scale=scale_weak, method="sing")$edfe
```

In each case the results largely recapitulate that seen under the wide-DFE although as expected the divergence between the distributions is more limited.

```
plot.wahl(edfe_cell_weak, means=means_weak, shape=shape_weak, scale=scale_weak, tau=tau)
```

```
plot.wahl(edfe_site_weak, means=means_weak, shape=shape_weak, scale=scale_weak, tau=tau)
```

```
plot.wahl(edfe_doub_weak, means=means_weak, shape=shape_weak, scale=scale_weak, tau=tau)
```

```
plot.wahl(edfe_pois_weak, means=means_weak, shape=shape_weak, scale=scale_weak, tau=tau)
```

```
plot.wahl(edfe_sing_weak, means=means_weak, shape=shape_weak, scale=scale_weak, tau=tau)
```

### 8 Segregating Variation

Gralka et al. (2016) is a key empirical paper demonstrating increased selection efficiency under colony growth. However, in that paper it is unclear whether the result depends on initiating experimental populations with a certain level of standing variation. In sexually reproducing organisms, beneficial spontaneous mutations are more likely to go extinct with higher drift, however, those that are destined to fixation actually end up fixing faster. While extinction is impossible without cell death, as modelled here, a similar process is possible where a rare beneficial mutation can get stochastically blocked out of the colony edge. However, the initial frequency of the beneficial allele may have been sufficiently large in Gralka et al. (2016) that at least one of the segregating beneficial alleles escapes being trapped and is destined to fixation. It is possible it would do so more quickly than in a well-mixed population, resulting in faster adaptation. In Gralka et al. (2016) the colony is formed from an initial well-mixed population of \(5\times10^4\) cells and left to grow until it reaches \(2\times10^8\) cells. In the initial population a beneficial mutation with selection coefficient \(0.08\) (experiments were also ran with \(s=-0.01\) and \(s=0.015\)) is present at a frequency of \(0.02\). In terms of generations, the initial population size is consistent with 15.6 doublings and the final population size is consistent with 27.6 doublings. Given that in our simulations only 16 generations of selection occur, we rescale the initial population size so that it occurs at 9.1 generations (\(N=\) 533). We then calculate the frequency of the beneficial mutations after 16 doublings for cell-based and site-based colony growth and single-step well-mixed growth.

```
edfe_cell_seg<-edfe_site_seg<-edfe_sing_seg<-rep(NA, nlines)

seg.N<-round(2^(16*log(5*10^4)/log(2*10^8)))

for(i in 1:nlines){
  edfe_cell_seg[i]<-sum(EdenSimCell(N=N, mu=0.02, means=0.08, shape=0, scale=0, seg.N=seg.N)[,"nmut"])/N
  edfe_site_seg[i]<-sum(EdenSimSite(N=N, mu=0.02, means=0.08, shape=0, scale=0, seg.N=seg.N)[,"nmut"])/N
  edfe_sing_seg[i]<-sum(EdenSimSing(N=N, mu=0.02, means=0.08, shape=0, scale=0, seg.N=seg.N)[,"nmut"])/N
  print(i)
}
```

Looking at the frequency distribution of the beneficial mutation in each of the three scenarios

```
par(mfrow=c(3,1))
breaks<-hist(edfe_cell_seg, xlab="Frequency", main="Cell-based Colony", breaks=100)$breaks  
hist(edfe_site_seg, breaks=breaks, xlab="Frequency", main="Site-based Colony")  
hist(edfe_sing_seg, breaks=breaks, xlab="Frequency", main="Single-step Well-mixed")
```

we can see that in colony-based growth the beneficial mutations often get ‘trapped’ in the centre of the colony early on and are then present at low frequency. However, if it escapes this ‘extinction’ it then reaches frequencies that are greater than observed in well-mixed growth, particularly under the cell-based scenario. However, the mean frequencies of the beneficial mutation under each scenario have the same rank order as the selection bias with spontaneous mutations (cell-based: 0.113, site-based: 0.022 and single-step: 0.042). This suggests that the faster adaptation seen under colony growth by Gralka et al. (2016) cannot be attributed solely to the focus on segregating versus spontaneous variation. However, in our down-scaled simulations, there are only 533 \(\times\) 0.02 = 11 beneficial mutations initially, on average. It is possible that with a higher number of initial cells the chance that all beneficial mutations get trapped is considerably smaller in which case the site-based model may also generate higher frequencies of the benefical mutations compared to well mixed growth. With an initial frequency of 20% there will be 533 \(\times\) 0.2 = 107 beneficial mutations initially, and the chance that several are on the periphery of the colony, and could fix, is much greater:

```
edfe_cell_seg2<-edfe_site_seg2<-edfe_sing_seg2<-rep(NA, nlines)

for(i in 1:nlines){
  edfe_cell_seg2[i]<-sum(EdenSimCell(N=N, mu=0.2, means=0.08, shape=0, scale=0, seg.N=seg.N)[,"nmut"])/N
  edfe_site_seg2[i]<-sum(EdenSimSite(N=N, mu=0.2, means=0.08, shape=0, scale=0, seg.N=seg.N)[,"nmut"])/N
  edfe_sing_seg2[i]<-sum(EdenSimSing(N=N, mu=0.2, means=0.08, shape=0, scale=0, seg.N=seg.N)[,"nmut"])/N
}
```

Histograms of the final frequencies show that the chance of all beneficial mutations getting trapped in the colony centre is much reduced. However, the rank order of the mean frequencies remain unchanged
(cell-based: 0.654, site-based: 0.212 and single-step: 0.343):

```
par(mfrow=c(3,1))
breaks<-hist(edfe_cell_seg2, xlab="Frequency", main="Cell-based Colony", breaks=100)$breaks  
hist(edfe_site_seg2, breaks=breaks, xlab="Frequency", main="Site-based Colony")  
hist(edfe_sing_seg2, breaks=breaks, xlab="Frequency", main="Single-step Well-mixed")
```

### 9 Generation Time

Mutation rate estimates from microbial MA have taken the number of generations to be \(log\_2(N)\). However, this is a lower limit on the number of generations, and is only achieved when all cells divide synchronously and every cell at the end has been through exactly the same number of cell divisions since the common ancestor . If there is variation in the number of cell divisions cells have been though, then the average number of generations will be greater than this minimum. Sackin’s index of tree imbalance (Shao and Sokal 1990), normalised by the final population size, is equal to the average number of generations. The index is known to increase with the amount of tree imbalance. Even in well-mixed growth, the tree will exhibit some imbalance just by chance, but when there is correlated variation in cell division rates down lineages, as is expected under spatially constrained growth, the amount of imbalance may be considerable. Indeed, under the most constrained growth where a population consists of all cells in a single line of descent (for example as produced by the mother machine - Robert et al. (2018)) the tree is ladder-like and the average number of generations is close to half the final number of final cells. To explore how the number of generations deviates from that assumed under synchronous growth we ran cell-based and site-based colony-growth simulations, with the mutation rate to zero so that we could explore generation number under neutrality. We also reran the single-step homogeneous growth model in which generation is defined consistently with the colony growth-models. In the absence of mutation, the single-step homogeneous growth model results in a neutral coalescent tree with as many tips as there are final cells in the population. The expectation of Sackin’s index for a neutral coalescent tree has been derived (Kirkpatrick and Slatkin 1993) leading to an expectation for the number of generations under single-step homogeneous growth:

\[
E[g] = 2\sum^{N}\_{i=2}\frac{1}{i}
\]

For \(\tau=16\), the expected number of generations is therefore 21.34. From results for the harmonic series, the expected number of generations is approximately \(2ln(N)-1+\gamma\) where \(\gamma\) is Euler’s constant. Since \(ln(N)=log\_2(N)ln(2)=\tau ln(2)\), \(E[g]\) is approximately linear in \(\tau\) (\(E[g]\approx2ln(2)\tau-1+\gamma\)) and as \(\tau\) becomes large the expected number of generations is close to \(2ln(2)\tau = 1.386\tau\), which is 22.18 for \(\tau=16\) and 38.82 for \(\tau=28\). Note that this is slightly smaller than that obtained by Armitage (1952) under asynchronous growth where the number of generations was obtained as \(\tau/log(2)\approx 1.443\tau\). We can simulate 100 colonies for \(\tau=\) 16 generations under both colony-based growth models and the single-step homogeneous growth model:

```
ngen_cell<-simulate.MA(nlines=100, tau=tau, mu=0, method="cell")$ngen
ngen_site<-simulate.MA(nlines=100, tau=tau, mu=0, method="site")$ngen
ngen_sing<-simulate.MA(nlines=100, tau=tau, mu=0, method="sing")$ngen
```

Plotting the distribution of the average number of generations shows that under colony growth the number of generations is almost two-orders of magnitude higher than the commonly assumed theoretical minimum (\(\tau=\) 16) and does not seem to strongly depend on whether a cell-based or site-based model is used.

```
par(mfrow=c(1,2))
hist(ngen_cell, main="Cell-based Colony", xlab="Average number of generations", breaks=15)
hist(ngen_site, main="Site-based Colony", xlab="Average number of generations", breaks=15)
```

Even under well-mixed growth the number of generations exceeds \(\tau\) by roughly the expected amount 33% (red line) which is close to the asymptotic (with respect to \(\tau\)) expectation (blue line).

```
hist(ngen_sing, main="Single-step Well-mixed", xlab="Average number of generations", breaks=15)
abline(v=2*sum(1/(2:N)), col="red")
abline(v=2*log(2)*tau, col="blue")
```

When \(\tau=1\) the number of generations must be exactly one, and so the discrepancy between \(\tau\) and the number of generations must grow with \(\tau\). Under well-mixed growth however, the results of Kirkpatrick and Slatkin (1993) show that the relationship between average generation number and \(\tau\) becomes roughly linear as \(\tau\) becomes large such that the discrepancy (as a proportion) stabilises. In order to assess how the average number of generations scales with \(\tau\) under colony growth we simulate 10 colonies for \(\tau\) ranging from 2 to 16:

```
ngen_sing_tau<-ngen_cell_tau<-ngen_site_tau<-1:(15*10)

for(i in 2:16){
  ngen_sing_tau[(i-2)*10+1:10]<-simulate.MA(nlines=10, N=2^i, mu=0, method="sing")$ngen
  ngen_cell_tau[(i-2)*10+1:10]<-simulate.MA(nlines=10, N=2^i, mu=0, method="cell")$ngen
  ngen_site_tau[(i-2)*10+1:10]<-simulate.MA(nlines=10, N=2^i, mu=0, method="site")$ngen
}
```

We can take the average number of generations per \(\tau\) and plot them against \(\tau\). As expected, under the single-step homogeneous model the results for Sackin’s index predict the number of generations well (red line) even if the asymptotic approximation is used (blue line).

```
ngen_sing_tau_av<-tapply(ngen_sing_tau, rep(2:16, each=10), mean)
plot(ngen_sing_tau_av~I(2:16), main="Single-step Well-mixed", xlab=expression(tau), ylab="Average number of generations")
lines(sapply(2^(2:16), function(x){2*sum(1/(2:x))})~I(2:16), col="red")
abline(0,1)
abline(-digamma(1)-1,2*log(2), col="blue")
```

As predicted by Gralka et al. (2016), the number of generations in an Eden model increases exponentially with \(\tau\) and this does not seem to depend on whether a cell-base or site-based model is used:

```
ngen_cell_tau_av<-tapply(ngen_cell_tau, rep(2:16, each=10), mean)
ngen_site_tau_av<-tapply(ngen_site_tau, rep(2:16, each=10), mean)
plot(ngen_cell_tau_av~I(2:16), ylim=range(c(ngen_cell_tau_av, ngen_site_tau_av)), xlab=expression(tau), ylab="Average number of generations", cex=1.5)
points(ngen_site_tau_av~I(2:16), col="red", cex=1.5)

legend("topleft", legend=c("Cell-based Colony", "Site-based Colony"), fill=c("black", "red"), inset=0.05)
```

Gralka et al. (2016) state that the number of generations is approximately proportional to \(\sqrt{N}\) which implies a log-linear relationship with \(\tau\): \(E[g]\propto \sqrt{N}\) such that \(log(E[g]) = c + \tau ln(2)/2\) where \(c\) is some constant. Indeed, this relationship seems to predict the number of generations well (green line) and is less than the expected asymptotic relationship under single-step homogenous growth (blue line), as expected:

```
plot(log(ngen_cell_tau_av)~I(2:16), ylim=range(log(c(ngen_cell_tau_av, ngen_site_tau_av))), xlab=expression(tau), ylab="log(Average number of generations)", cex=1.5)
points(log(ngen_site_tau_av), col="red", cex=1.5)
abline(-log(2),log(2), col="blue")
abline(0, log(2)/2, col="green")

legend("topleft", legend=c("Cell-based Colony", "Site-based Colony"), fill=c("black", "red", "blue"), inset=0.05)
```

#### 9.1 Scaling of selection efficiency with \(\tau\)

Under well-mixed growth, the amount of selection bias will increase almost linearly with \(\tau\) across the range of \(\tau\)’s used in microbial MA. However, for colony growth it seems likely that the selection bias will accelerate with \(\tau\) as the number of generations over which selection operates also accelerates. To assess this we can simulate both site-based and cell-based growth under a range of \(\tau\)’s (8 to 17) and estimate \(\tau\_e\).

```
edfe_cell_tau<-8:17
edfe_site_tau<-8:17
for(i in 8:17){
edfe_cell_tmp<-simulate.MA(nlines=1000, N=2^i, mu=mu, means=means, shape=shape, scale=scale, method="cell")$edfe
edfe_cell_tau[i-7]<-fit.wahl(edfe_cell_tmp, means=means, shape=shape, scale=scale, tau=TRUE)$est
edfe_site_tmp<-simulate.MA(nlines=1000, N=2^i, mu=mu, means=means, shape=shape, scale=scale, method="site")$edfe
edfe_site_tau[i-7]<-fit.wahl(edfe_site_tmp, means=means, shape=shape, scale=scale, tau=TRUE)$est
}
```

We can plot \(\tau\_e\) against \(\tau\) for both cell-based (black) and site-based (red) models of colony growth with a dashed black 1:1 line:

```
plot(edfe_cell_tau~I(8:17), ylim=range(c(8:17, edfe_cell_tau, edfe_site_tau)), type="l", lwd=2, ylab=expression(paste("Effective population doublings: ", tau[e])), xlab=expression(paste("Population doublings: ", tau)))
lines(edfe_site_tau~I(8:17), col="red", type="l", lwd=2)
abline(0, 1, lty=2)
legend("topleft", legend=c("Cell-based Colony", "Site-based Colony"), fill=c("black", "red"), inset=0.05)
```

It is apparent that the difference between cell-based colony growth and well-mixed colony growth increases with increasing \(\tau\), likely due to the fact that the number of generations is scaling exponentially with \(\tau\) under colony growth. For small \(\tau\) (<12), the effect of drift seems to outweigh the effect of increasing the number of generations but for the range of \(\tau\)’s typically used in microbial MA, the selection bias is stronger (\(\tau\_e>\tau\)) under cell-based colony growth. On the other hand, site-based colony growth seems much less sensitive to \(\tau\) and selection bias will most likely always be less than in well-mixed growth across a reasonable range of \(\tau\)’s.

For cell-based colony growth, the effect of the exponential increase in the number of generations with \(\tau\) on selection bias raises an important issue. Even when \(\tau=16\), which for *E. coli* would take around 9 hours of colony growth under optimal conditions, the Eden model is predicting 181 generations. This is not realistic as it would require the average generation time to be about 3 minutes, far shorter than what is possible. Since the number of generations appears unrealistic in the simulations, we could imagine that in real systems the threshold \(\tau\) under which the selection bias is greater under cell-based colony growth may be larger.

### 10 Meta-analysis of non-synonymous/synonymous ratios

In the main manuscript we present a meta-analysis of non-synonymous/synonymous ratios. We collected data on observed (\(r\_{obs}\)) and expected ratios (\(r\_{exp}\)) of non-synonymous versus synonymous mutations from published MA experiments together with the total number of coding mutations. Since non-synonymous mutations are expected to be more deleterious, on average, than synonymous mutations we expect the observed ratio to be less than expected if selection bias exists. To test this, we fitted binomial generalised linear mixed models with logit link to the number of non-synonymous and synonymous mutations per study. Observation-level random effects were fitted to soak up any overdispersion.

#### 10.1 Standard meta-analysis

In the first analysis we fitted \(ln(r\_{exp})\) as an offset such that the intercept is the log of the expected deviation of \(r\_{obs}\) from \(r\_{exp}\).

```
tau_dat<-read.xlsx(file.path(root, "Data/Raw/colonyMA.xlsx"))

tau_dat$nsyn<-round(tau_dat$coding_bps*tau_dat$nsyn_ratio/(tau_dat$nsyn_ratio+1))
# number of non-synonymous mutations

tau_dat$syn<-tau_dat$coding_bps-tau_dat$nsyn
# number of synonymous mutations

tau_dat$obs<-as.factor(1:nrow(tau_dat))
# observation-level factor

m_tau_dat<-glmer(cbind(nsyn, syn)~offset(log(nsyn_exp))+(1|obs), family="binomial", data=tau_dat)
# fit model with constant deficit (intercept only)

summary(m_tau_dat)
```

```
## Generalized linear mixed model fit by maximum likelihood (Laplace
##   Approximation) [glmerMod]
##  Family: binomial  ( logit )
## Formula: cbind(nsyn, syn) ~ offset(log(nsyn_exp)) + (1 | obs)
##    Data: tau_dat
## 
##       AIC       BIC    logLik -2*log(L)  df.resid 
##     200.4     203.1     -98.2     196.4        27 
## 
## Scaled residuals: 
##     Min      1Q  Median      3Q     Max 
## -2.0446 -0.9607 -0.1255  0.6769  1.2226 
## 
## Random effects:
##  Groups Name        Variance Std.Dev.
##  obs    (Intercept) 0.001917 0.04378 
## Number of obs: 29, groups:  obs, 29
## 
## Fixed effects:
##             Estimate Std. Error z value Pr(>|z|)   
## (Intercept) -0.08041    0.02588  -3.107  0.00189 **
## ---
## Signif. codes:  0 '***' 0.001 '**' 0.01 '*' 0.05 '.' 0.1 ' ' 1
```

The intercept is significantly negative indicating a significant deficit of non-synonymous mutations: exponentiating the intercept tell us that the observed ratio is only 92.27% its expected value.

In the main manuscript we developed a multinomial model than can fully control for the mutational spectrum when assessing whether non-synonymous mutations are under-represented compared to synonymous mutations (see Multinomial Model SI). When applying this model to *E. coli* data we found that a model that does not control for differences in transition/transversion rate predicts a greater deficit of non-synonymous mutations. Since only a subset of studies in the meta-analysis controlled for differences in transition/transversion rates, we may worry that the deficit estimated in the meta-analysis reflects a failure to control for the mutational spectrum, rather than selection against non-synonymous mutations. If this were the case, we may expect to see a reduced deficit in studies that control for differences in transition/transversion rates.

```
tau_dat$TrTv<-grepl("2", tau_dat$nsyn_con)
# indicator of whether a study controlled for differences in transition/transversion rates (TRUE) or not (FALSE)

m_tau_dat2<-glmer(cbind(nsyn, syn)~offset(log(nsyn_exp))+TrTv+(1|obs), family="binomial", data=tau_dat)

summary(m_tau_dat2)
```

```
## Generalized linear mixed model fit by maximum likelihood (Laplace
##   Approximation) [glmerMod]
##  Family: binomial  ( logit )
## Formula: cbind(nsyn, syn) ~ offset(log(nsyn_exp)) + TrTv + (1 | obs)
##    Data: tau_dat
## 
##       AIC       BIC    logLik -2*log(L)  df.resid 
##     198.8     202.9     -96.4     192.8        26 
## 
## Scaled residuals: 
##     Min      1Q  Median      3Q     Max 
## -2.4792 -0.7961 -0.1107  0.6978  1.4353 
## 
## Random effects:
##  Groups Name        Variance Std.Dev.
##  obs    (Intercept) 0        0       
## Number of obs: 29, groups:  obs, 29
## 
## Fixed effects:
##             Estimate Std. Error z value Pr(>|z|)  
## (Intercept) -0.03921    0.02807  -1.397   0.1624  
## TrTvTRUE    -0.09204    0.04539  -2.028   0.0426 *
## ---
## Signif. codes:  0 '***' 0.001 '**' 0.01 '*' 0.05 '.' 0.1 ' ' 1
## 
## Correlation of Fixed Effects:
##          (Intr)
## TrTvTRUE -0.618
## optimizer (Nelder_Mead) convergence code: 0 (OK)
## boundary (singular) fit: see help('isSingular')
```

Perhaps surprisingly, the deficit in studies that control for differences in transition/transversion rates is larger, not smaller, perhaps because studies that control for differences in transition/transversion rates are also more likely to control for codon usage and GC content. Either way, it seems unlikely that the deficit of non-synonymous mutations is due to a failure to control for differences in transition/transversion rates and indeed more careful controls may well suggest the deficit due to selection is larger than we estimate.

#### 10.2 Selection bias informed meta-analysis

To put the deficit estimated from the standard meta-analysis into context we fitted an additional analysis where the log of the observed ratio is assumed to be equal to \(ln(b(s | \tau)r\_{exp})\) with \(s\) estimated. This gives an estimate of \(s\) assuming that all synonymous mutations are neutral (\(s=0\)) and all non-synonymous mutations have the same \(s\) and are subject to the selection bias predicted under homogeneous growth. This is a non-linear model and so we provide the function `meta.wahl` to fit this model which relies on a call to `optim` to estimate \(s\):

```
meta.wahl<-function(data){

  if(any(!c("nsyn", "syn", "nsyn_exp", "tau")%in%colnames(data))){
    stop("nsyn, syn, nsyn_exp and tau must appear in data")
  }

  data$obs<-1:nrow(data)

  fn<-function(s, data=data){

    x<-wahl.bias(s, tau=data$tau, log=TRUE)
    LL<-logLik(glmer(cbind(nsyn, syn)~offset(log(nsyn_exp)+x)-1+(1|obs), family="binomial", data=data))
     return(-LL)
  }

  out<-optim(-0.01, fn, data=data, hessian=TRUE)

  LL.int<-logLik(glmer(cbind(nsyn, syn)~offset(log(nsyn_exp))+(1|obs), family="binomial", data=data))

  res<-list(s=out$par,               # estimated selection coefficient
            se=1/sqrt(out$hessian),  # standard error
            logLik=-out$value,       # log-likelihood
            dAIC=as.numeric(-2*(-out$value-LL.int)),  # AIC difference with intercept only model
            p=2*pnorm(-abs(out$par*sqrt(out$hessian)))) # two-tailed p-value


  return(res)

}
```

We can fit this model to the published data (note one study is discarded because \(\tau\) was not reported):

```
m_tau_dat3<-meta.wahl(data=subset(tau_dat, !is.na(tau)))
m_tau_dat3
```

```
## $s
## [1] -0.01095312
## 
## $se
##             [,1]
## [1,] 0.003340783
## 
## $logLik
## [1] -98.07196
## 
## $dAIC
## [1] -0.2568752
## 
## $p
##             [,1]
## [1,] 0.001043197
```

The best estimate is -0.011 \(\pm\) 0.003 and the model is a very slight improvement over the intercept only model (\(\Delta\)AIC=-0.257). The estimated \(s\) should be seen as an upper bound on the mean \(s\) of non-synonymous mutations *if* the homogeneous growth model is accurate. \(s\) would have been estimated as more negative had we been able to factor in selection against synonymous mutations, accounted for variation in \(s\) around the mean (since \(b(s|\tau)\) is convex), and accommodated the drift component of homogeneous growth.

### 11 Power

Despite the meta-analysis showing a clear deficit of non-synonymous to synonymous mutations, relative to their expectation, few, if any studies, have found a significant deficit. However, what sort of sample sizes would be required to detect it? We can look at the power of a binomial test where the null odds that a mutation is non-synonymous is \(r\_{exp}=P\_{ns}/(1-P\_{ns})\) and the odds under the alternate hypothesis is specified. Given a total of `nmu` synonymous and non-synonymous mutations of which a proportion `pns` are non-synonymous, the function `power.bias` returns the power of such a test (given a significance threshold of 0.05). If `s` and `tau` are given, the alternate odds is calculated to be \(b(s, \tau)r\_{exp}\). If \(s\) is not given the alternate odds need to specified through `alt.pns` (the frequency of non-synonymous mutations under the alternate hypothesis). Since, a deficit of non-synonymous mutations is expected the default is a one-tailed test (`one.tailed=TRUE`).

```
power.bias<-function(nmu, pns, s=NULL, tau=NULL, alt.pns=NULL, one.tailed=TRUE){

  odds<-pns/(1-pns)

  if(is.null(s)){
    if(is.null(alt.pns)){stop("if s is NULL, alt.pns needs to be given")}
  }
  if(!is.null(s)){
    if(!is.null(alt.pns)){stop("if s is given, alt.pns should be NULL")}
    if(is.null(tau)){stop("if s is given, tau should also be given")}
  }

  if(!is.null(s)){
    adj.odds<-odds*wahl.bias(s=s, tau=tau)

    alt.pns<-adj.odds/(1+adj.odds)
  }

  if(one.tailed){

    critical<-qbinom(0.05, size=nmu, prob=pns)

    power<-pbinom(critical-1, size=nmu, prob=alt.pns)
  }else{
    critical<-qbinom(0.025, size=nmu, prob=pns)

    power<-pbinom(critical-1, size=nmu, prob=alt.pns)

    critical<-qbinom(0.975, size=nmu, prob=pns)

    power<-power+1-pbinom(critical-1, size=nmu, prob=alt.pns)


  }

  return(power)
}
```

With \(\tau=28\) and a 0.76 chance of a mutation being non-synonymous in the absence of selection (Foster et al. 2015), the chance of detecting any bias when non-synonymous mutations confer a 1% disadvantage is less than the desired 80% unless the total number of mutations exceeds 3,000. If \(\tau=10\) then more than \(23,000\) mutations would be required:

```
nmu<-seq(10, 2000, 100)
power<-nmu
for(i in 1:length(nmu)){
power[i]<-power.bias(nmu[i], pns=0.76, s=-0.01, tau=28)
}
plot(power~nmu, ylab="Power", xlab="Number of mutations", type="l", ylim=c(0,1))
text(expression(paste(tau, "=28")), y=power[i]*1.06, x=1800)
for(i in 1:length(nmu)){
power[i]<-power.bias(nmu[i], pns=0.76, s=-0.01, tau=10)
}
lines(power~nmu, lty=2)
text(expression(paste(tau, "=10")), y=power[i]*1.12, x=1800)

abline(h=0.8, col="red")
```

In Table 1 of the main manuscript we summarise previous studies that have looked for selection bias in microbial MA. Below, we calculate the power of these studies given a selection coefficient of \(-0.01\) and compare it to the power of our own study (0.268: \(P\_{NS}\)=0.727 (given an expected ratio of 2.67: Table S1 in Foster et al. (2015)) 641 coding mutations and \(\tau=28\)):

```
tau_dat$power<-power.bias(nmu=tau_dat$coding_bps, pns=tau_dat$nsyn_exp/(1+tau_dat$nsyn_exp), tau=tau_dat$tau, s=-0.01)
hist(tau_dat$power, main="", xlab="Power", xlim=c(0,0.3), breaks=30)
abline(v=power.bias(641, pns=0.727, tau=28, s=-0.01), col="red")
```

Our study has the greatest power to detect selection bias, although note that many studies do not give all of the relevant information to calculate power.

### References

Armitage, P. 1952. “The Statistical Theory of Bacterial Populations Subject to Mutation.” *Journal of the Royal Statistical Society: Series B (Methodological)* 14 (1): 1–33.

Colyer, Blair, Maciej Bak, David Basanta, and Robert Noble. 2024. “A Seven-Step Guide to Spatial, Agent-Based Modelling of Tumour Evolution.” *Evolutionary Applications* 17 (5): e13687.

Couce, Alejandro, Anurag Limdi, Melanie Magnan, Siân V Owen, Cristina M Herren, Richard E Lenski, Olivier Tenaillon, and Michael Baym. 2024. “Changing Fitness Effects of Mutations Through Long-Term Bacterial Evolution.” *Science* 383 (6681): eadd1417.

Eden, Murray. 1961. “A Two-Dimensional Growth Process.” *Dynamics of Fractal Surfaces* 4 (223-239): 598.

Foster, Patricia L, Heewook Lee, Ellen Popodi, Jesse P Townes, and Haixu Tang. 2015. “Determinants of Spontaneous Mutation in the Bacterium Escherichia Coli as Revealed by Whole-Genome Sequencing.” *Proceedings of the National Academy of Sciences* 112 (44): E5990–99.

Gralka, Matti, Fabian Stiewe, Fred Farrell, Wolfram Möbius, Bartlomiej Waclaw, and Oskar Hallatschek. 2016. “Allele Surfing Promotes Microbial Adaptation from Standing Variation.” *Ecology Letters* 19 (8): 889–98.

Hallatschek, Oskar, Pascal Hersen, Sharad Ramanathan, and David R Nelson. 2007. “Genetic Drift at Expanding Frontiers Promotes Gene Segregation.” *Proceedings of the National Academy of Sciences* 104 (50): 19926–30.

Kim, Wook, Fernando Racimo, Jonas Schluter, Stuart B Levy, and Kevin R Foster. 2014. “Importance of Positioning for Microbial Evolution.” *Proceedings of the National Academy of Sciences* 111 (16): E1639–47.

Kirkpatrick, Mark, and Montgomery Slatkin. 1993. “Searching for Evolutionary Patterns in the Shape of a Phylogenetic Tree.” *Evolution* 47 (4): 1171–81.

Lavrentovich, Maxim O, Mary E Wahl, David R Nelson, and Andrew W Murray. 2016. “Spatially Constrained Growth Enhances Conversional Meltdown.” *Biophysical Journal* 110 (12): 2800–2808.

Robert, Lydia, Jean Ollion, Jerome Robert, Xiaohu Song, Ivan Matic, and Marina Elez. 2018. “Mutation Dynamics and Fitness Effects Followed in Single Cells.” *Science* 359 (6381): 1283–86.

Shao, Kwang-Tsao, and Robert R Sokal. 1990. “Tree Balance.” *Systematic Zoology* 39 (3): 266–76.

Wahl, Lindi M, and Deepa Agashe. 2022. “Selection Bias in Mutation Accumulation.” *Evolution* 76 (3): 528–40.
