## Supplementary material for "Selection bias in microbial mutation accumulation studies and the impact of colony growth": Multinomial SI: Multinomial_Model_SI.html

Using mlogit to Model Mutational Spectra


### Using mlogit to Model Mutational Spectra

In this workbook we implement and explain a set of multinomial models for analysing mutational data from mutational accumulation lines. The models are fit using the mlogit R package (Croissant 2020).

```
library(mlogit)
library(dfidx)
```

We have sequence data from \(471\) *Escherichia coli* mutation accumulation (MA) lines where there are four outcomes (A, C, G, T) at each of 4,641,652 sites. In the language of mlogit, sites are referred to as *individuals* or *choice situations*, and the bases as *alternatives*. Ideally, at each site we would record the \(471\) outcomes. Since most sites have no mutations, the majority of sites would have all 471 outcomes as a single base - the base of the original sequence. For a minority of sites, 470 outcomes will be the original base and one outcome will be an alternative base caused by a mutation in a single line. In a very small number of cases more than one mutation may be observed at a site. Unfortunately, mlogit can only deal with multinomial outcomes of size one (i.e for each choice situation (site) only one alternative (base) is observed). However, when the chance of observing a mutation is small (because the mutation rate is small and/or the total number of MA generations is small) - as in our case - then multiple mutations at a site are extremely rare and can be ignored. Then, we can make the outcome the base of the original sequence if no mutations have been observed, and if a mutation has been observed, we can make the outcome the base of the new mutation. If the total number of MA generation is \(g\) then the probability of no observed mutations at a site is \((1-\mu)^g\) and the probability of one mutation is \(g\mu(1-\mu)^{g-1}\). By ignoring multiple mutations we approximate the latter probability as \(1-(1-\mu)^{g}\), with the approximation being exact as \(g\mu\) becomes small. For our *E. coli* MA data set, \(g\) is of the order \(10^6\) and previous estimates of the mutation rate have been of the order \(2\times 10^{-10}\) (Lee et al. 2012) giving \(g\mu\) of the order \(2\times 10^{-4}\). The relative error in the approximation is therefore vanishingly small \(g\mu(1-\mu)^{g-1}\) = \(1.9996\times 10^{-4}\) and \(1-(1-\mu)^{g}\) = \(1.9998\times 10^{-4}\).

### 1 Data-frame structure

mlogit requires the data to be in long-format where each row corresponds to a specific combination of choice situation (site) and alternative (base). With a genome of 4,641,652 sites and four bases this results in data-frame with 18,566,608 rows (See Ecoli\_Data\_SI for details - the data-frame is large (1.5Gb) and is not hosted on GitHub):

```
  long_dat<-read.csv(file.path(root, "Data/Intermediate/long_dat.csv"))
```

The data frame needs to be an object of class dfidx where the two indicies (choice situation and alternative) are defined through the argument `idx`:

```
  long_dat <- dfidx(long_dat, shape = "long", idx=c("site", "base"))
```

The data frame contains the response variable `new_seq` which for each site contains a single one (for the base of the original sequence, or the mutated base) and three zeros. For example, at site 570 the original sequence has a C and no mutations were seen in any line:

```
long_dat[which(long_dat$idx$site==570), c("new_seq", "mutate")]
```

```
##   new_seq mutate   idx
## 1       0      1 570:A
## 2       1      0 570:C
## 3       0      1 570:G
## 4       0      1 570:T
## 
## ~~~ indexes ~~~~
##   site base
## 1  570    A
## 2  570    C
## 3  570    G
## 4  570    T
## indexes:  1, 2
```

`new_seq` is one for base C at site 570 and `mutate` is zero. At site 571 the original sequence has a G and a mutation was seen in a line:

```
long_dat[which(long_dat$idx$site==571), c("new_seq", "mutate")]
```

```
##   new_seq mutate   idx
## 1       0      1 571:A
## 2       1      1 571:C
## 3       0      0 571:G
## 4       0      1 571:T
## 
## ~~~ indexes ~~~~
##   site base
## 1  571    A
## 2  571    C
## 3  571    G
## 4  571    T
## indexes:  1, 2
```

`mutate` is zero for G indicating it was the original sequence but `new_seq` is one for C indicating the mutation \(G\rightarrow C\) ocurred in a line. Note the original sequence and any mutations are always given for the leading strand.

### 2 Model

The underlying logit model has the form

\[P\_{ij} = exp(\eta\_{ij})/\sum\_{k=1}^{4}exp(\eta\_{ik})\]

where \(P\_{ij}\) is the probability that site \(i\) has base \(j\) as the outcome, and \(\eta\_{ij}\) is the linear predictor for that outcome. The denominator sums the four exponentiated linear predictors for the site (one for each base).

The model formula has `new_seq` on the left-hand side and the right-hand size has three parts separated by pipes that define the form for the linear predictor (see here)

- Part A) a covariate \(x\) that varies over sites and/or bases for which you want a single regression parameter: a main effect of \(x\) where \(x\) may vary across bases within a site.
- Part B) a covariate \(x\) that varies over sites only for which each base has a different regression parameter: an interaction between \(x\) and base where \(x\) does not vary across bases within a site.
- Part C) a covariate \(x\) that varies over sites and bases for which each base has a different regression parameter: an interaction between \(x\) and base where \(x\) can vary across bases within a site.

#### 2.1 Basic Models

In the simplest case we could just estimate the odds of seeing a mutation:

```
  m.basic <- mlogit(new_seq ~ mutate | 0 | 0, data=long_dat)
```

```
summary(m.basic)
```

```
## 
## Call:
## mlogit(formula = new_seq ~ mutate | 0 | 0, data = long_dat, method = "nr")
## 
## Frequencies of alternatives:choice
##       A       C       G       T 
## 0.24620 0.25424 0.25366 0.24591 
## 
## nr method
## 11 iterations, 0h:2m:26s 
## g'(-H)^-1g = 1.05E-10 
## gradient close to zero 
## 
## Coefficients :
##         Estimate Std. Error z-value  Pr(>|z|)    
## mutate -9.681663   0.033926 -285.38 < 2.2e-16 ***
## ---
## Signif. codes:  0 '***' 0.001 '**' 0.01 '*' 0.05 '.' 0.1 ' ' 1
## 
## Log-Likelihood: -9282.4
```

This model has the form

\[P\_{ij} = exp(\textrm{mutate}\_{ij}\beta^{(1)})/\sum\_{k=1}^{4}exp(\textrm{mutate}\_{ij}\beta^{(1)})\]

where \(\textrm{mutate}\_{ij}=1\) if base \(j\) is not the original base (i.e would constitute a mutation). For every site, \(\textrm{mutate}\_{ij}=0\) for one base (the original base) and is one otherwise. Consequently, the denominator is \(3exp(\beta^{(1)})+1\) and the numerator is either 1 (if the base is the original base) or \(exp(\beta^{(1)})\) if the base constitute a mutation. For a base \(j\) that constitutes a mutation at site \(i\) the probability is therefore:

\(P\_{ij} = exp(\beta^{(1)})/(3exp(\beta^{(1)})+1)\)

Since there are three such bases, the probability that a mutation happens at site \(i\) is the sum of the three probabilities. Since the probabilities are assumed constant, the probability of observing a mutation is estimated to be three times the above quantity.

```
b<-coef(m.basic)["mutate"]
3*exp(b)/(3*exp(b)+1)
```

```
##       mutate 
## 0.0001872178
```

This can be compared to the observed frequency of mutations

```
sum(long_dat$new_seq & long_dat$mutate)/4641652
```

```
## [1] 0.0001872178
```

and is identical, as would be expected in such a simple model.

This is the frequency of observing at least one mutation across 266 lines which in aggregate involved 939,774 MA generations. This frequency, \(p\), is equal \(1-(1-\mu)^{939774}\) giving an estimate of \(\mu\) as \(1-exp(log(1-p)/939774)=\) \(1.992\times 10^{-10}\).

A more complicated model allows the bases to have different mutation rates:

```
  m.basic2 <- mlogit(new_seq ~ mutate | 0 | mutate, data=long_dat)
```

```
summary(m.basic2)
```

```
## 
## Call:
## mlogit(formula = new_seq ~ mutate | 0 | mutate, data = long_dat, 
##     method = "nr")
## 
## Frequencies of alternatives:choice
##       A       C       G       T 
## 0.24620 0.25424 0.25366 0.24591 
## 
## nr method
## 11 iterations, 0h:2m:47s 
## g'(-H)^-1g = 4.98E-10 
## gradient close to zero 
## 
## Coefficients :
##           Estimate Std. Error   z-value Pr(>|z|)    
## mutate   -9.587134   0.064552 -148.5181  < 2e-16 ***
## mutate:C -0.238803   0.097542   -2.4482  0.01436 *  
## mutate:G -0.123569   0.094492   -1.3077  0.19097    
## mutate:T -0.034291   0.092072   -0.3724  0.70957    
## ---
## Signif. codes:  0 '***' 0.001 '**' 0.01 '*' 0.05 '.' 0.1 ' ' 1
## 
## Log-Likelihood: -9278.9
```

Here the main effect of \(\textrm{mutate}\) is the coefficient for the base-line category (in this case A) and the remaining coefficients are deviations from this. C and G have lower mutation rates than A (C significantly so) but T is very similar. The coefficients are perhaps more easily understand in terms of the frequencies of mutated bases had they been at equal frequency in the original sequence:

```
p.base<-exp(coef(m.basic2)[1])
p.base<-c(p.base, exp(coef(m.basic2)[1]+coef(m.basic2)[-1]))
p.base<-p.base/sum(p.base) 
p.base
```

```
##    mutate  mutate:C  mutate:G  mutate:T 
## 0.2749049 0.2165068 0.2429503 0.2656380
```

#### 2.2 Advanced Models

The model we used in the main manuscript is considerably more complicated than these basic models, and, with appropriate predictors, can be formulated using only Part A) in the model formula. One set of two-level predictors have fairly straightforward interpretations: factors that distinguish coding from non-coding sites (`coding_type`) sites in highly expressed genes or not (`HEG`) and whether the outcome base would constitute a transition or transversion (`trans`) or a non-synonymous change or not (`nonsyn`). The remaining predictors allow the chance of mutation to depend on the original base and the surrounding sequence. These predictors are set up in a way where the sequence properties are strand independent (lagging or leading) in the baseline model, and any effect of the leading sequence properties are modelled as deviations from this.

The predictor `comb_REF` specifies whether the original base on the leading strand is \(\{\)A, T\(\}\) or \(\{\)C, G\(\}\). In the absence of strand-specificity the two bases in a pair should be equivalent. For example, imagine the sequence

\(5'\) ————— A —————– \(3'\) leading strand

\(3'\) ————— T —————– \(5'\) lagging strand

Since the original sequence is encoded for the leading strand the base is A. However, the sequence

\(5'\) ————— T —————– \(3'\) leading strand

\(3'\) ————— A —————– \(5'\) lagging strand

also has an A, albeit on the lagging strand. If A affects the chance of mutation in a strand-independent manner then these two sequences (with leading A or T) should be identical. Therefore, we can combine information on the leading nucleotide into two main categories encoded by `comb_REF`:

| comb\_REF | Leading Nucleotide | Lagging Nucleotide |
| --- | --- | --- |
| AT | A | T |
| AT | T | A |
| CG | C | G |
| CG | G | C |

The predictors `leading_T` and `leading_G` are indicator variables specifying whether the original base on the leading strand is T or G respectively. These can be used to allow mutation rate/type to depend on properties of the base in the leading strand.

The predictors `AT_nb`, `TA_nb`, `CG_nb` and `GC_nb` are numerical with values 0, 1 or 2 and represent information about neighbouring bases. As with `comb_REF` they encode information about the effect of neighbouring bases when strand identity is not important.

Imagine the sequence

\(5'\) ————– T A C —————- \(3'\) leading strand

\(3'\) ————– A T G —————- \(5'\) lagging strand

If we consider a mutation at the central site, on the leading strand the 5’ neighbour is T and on the lagging strand the 5’ neighbour is G. However, since the 5’ neighbour of one strand is the 3’ neighbour on the other strand, we can record this information as the 5’ and 3’ neighbour on the leading strand. When strand identity does not matter a 5’ T is equivalent to a 3’ A (since this implies the lagging strand has a 5’ T). We can write this effect as \(T|A\) to indicate it will be present if there is a 5’ T *or* a 3’ A. A similar logic can applied to other combinations of neighbours:

| Effect | Leading 5’ Neighbour | Leading 3’ Neighbour |
| --- | --- | --- |
| A\|T | A | T |
| T\|A | T | A |
| C\|G | C | G |
| G\|C | G | C |

Each site is assigned two of these effects depending on its two neighbours. For example, consider these sequences:

Example 1.1

\(5'\) ————– T A G —————- \(3'\) leading strand

\(3'\) ————– A T C —————- \(5'\) lagging strand

Example 2.1

\(5'\) ————– A A G —————- \(3'\) leading strand

\(3'\) ————– T T C —————- \(5'\) lagging strand

Example 3.1

\(5'\) ————– G A C —————- \(3'\) leading strand

\(3'\) ————– C T G —————- \(5'\) lagging strand

These are assigned the following values where the columns are the predictor variables `AT_nb`, `TA_nb`, `CG_nb` and `GC_nb`:

| Example | A\|T | T\|A | C\|G | G\|C |
| --- | --- | --- | --- | --- |
| 1.1 | 0 | 1 | 1 | 0 |
| 2.1 | 1 | 0 | 1 | 0 |
| 3.1 | 0 | 0 | 0 | 2 |

If we reverse the strands, we get the same answer:

Example 1.2

\(5'\) ————– C T A —————- \(3'\) leading strand

\(3'\) ————– G A T —————- \(5'\) lagging strand

Example 2.2

\(5'\) ————– C T T —————- \(3'\) leading strand

\(3'\) ————– G A A —————- \(5'\) lagging strand

Example 3.2

\(5'\) ————– G T C —————- \(3'\) leading strand

\(3'\) ————– C A G —————- \(5'\) lagging strand

| Example | A\|T | T\|A | C\|G | G\|C |
| --- | --- | --- | --- | --- |
| 1.2 | 0 | 1 | 1 | 0 |
| 2.2 | 1 | 0 | 1 | 0 |
| 3.2 | 0 | 0 | 0 | 2 |

Note that all four predictors are not identifiable and we drop `AT_nb` which generates the reference category AAT or ATT if `leading_T` is not fitted, or AAT if not. The predictor `leading_minus_neighbor` is a four level factor giving the original base immediately 5’ of the focal site on the leading strand. It can be used to allow mutation rate/type to depend on properties of the neighbours on the leading strand. The category A is dropped giving rise to the same reference category as above.

We can start with a model where there is no strand specificity. We can fit `mutate` as a main effect which models the log odds of a mutation in the reference category. We can then interact `mutate` with our predictors which then increment the odds of mutation. The reference category is a transition mutation in a non-coding A or T with a 5’ A and 3’ T:

```
  m1 <- mlogit(new_seq ~ mutate+mutate:(coding_type + HEG + CG_nb + GC_nb + TA_nb + trans + nonsyn + comb_REF) | 0 | 0, data=long_dat)
```

```
summary(m1)
```

```
## 
## Call:
## mlogit(formula = new_seq ~ mutate + mutate:(coding_type + HEG + 
##     CG_nb + GC_nb + TA_nb + trans + nonsyn + comb_REF) | 0 | 
##     0, data = long_dat, method = "nr")
## 
## Frequencies of alternatives:choice
##       A       C       G       T 
## 0.24620 0.25424 0.25366 0.24591 
## 
## nr method
## 11 iterations, 0h:3m:27s 
## g'(-H)^-1g = 4.06E-07 
## gradient close to zero 
## 
## Coefficients :
##                                   Estimate Std. Error  z-value  Pr(>|z|)    
## mutate                           -9.085055   0.120938 -75.1217 < 2.2e-16 ***
## mutate:coding_typeprotein_coding -0.716177   0.102222  -7.0061  2.45e-12 ***
## mutate:HEGTRUE                    0.060220   0.152825   0.3940   0.69355    
## mutate:CG_nb                      0.151271   0.072683   2.0812   0.03741 *  
## mutate:GC_nb                      0.592476   0.068932   8.5951 < 2.2e-16 ***
## mutate:TA_nb                      0.095109   0.074720   1.2729   0.20306    
## mutate:transtransversion         -0.845282   0.068766 -12.2921 < 2.2e-16 ***
## mutate:nonsyn                    -0.065645   0.090596  -0.7246   0.46870    
## mutate:comb_REFCG                 0.021747   0.068712   0.3165   0.75163    
## ---
## Signif. codes:  0 '***' 0.001 '**' 0.01 '*' 0.05 '.' 0.1 ' ' 1
## 
## Log-Likelihood: -9117.3
```

The model provides good evidence that mutations are less likely to happen in coding regions and transversions are less likely than transitions. Perhaps surprisingly, although \(\{\)C, G\(\}\) sites have higher mutation rate than \(\{\)A, T\(\}\) sites it is not significant. On the other hand, neighbouring bases seem to have a strong effect: having a leading 5’ G or 3’ C increases the mutation rate relative to having a leading 5’ A or 3’ T.

We can also add strand specificity to this model:

```
  m2 <- mlogit(new_seq ~ mutate+mutate:(coding_type + HEG + CG_nb + GC_nb + TA_nb + trans + nonsyn + comb_REF + leading_minus_neighbor+leading_T+leading_G) | 0 | 0, data=long_dat)
```

```
summary(m2)
```

```
## 
## Call:
## mlogit(formula = new_seq ~ mutate + mutate:(coding_type + HEG + 
##     CG_nb + GC_nb + TA_nb + trans + nonsyn + comb_REF + leading_minus_neighbor + 
##     leading_T + leading_G) | 0 | 0, data = long_dat, method = "nr")
## 
## Frequencies of alternatives:choice
##       A       C       G       T 
## 0.24620 0.25424 0.25366 0.24591 
## 
## nr method
## 11 iterations, 0h:4m:14s 
## g'(-H)^-1g = 6.83E-07 
## gradient close to zero 
## 
## Coefficients :
##                                   Estimate Std. Error  z-value  Pr(>|z|)    
## mutate                           -9.033299   0.129738 -69.6275 < 2.2e-16 ***
## mutate:coding_typeprotein_coding -0.716251   0.102235  -7.0059 2.454e-12 ***
## mutate:HEGTRUE                    0.060583   0.152832   0.3964   0.69181    
## mutate:CG_nb                      0.073227   0.102651   0.7134   0.47562    
## mutate:GC_nb                      0.560472   0.094709   5.9178 3.263e-09 ***
## mutate:TA_nb                      0.014694   0.104465   0.1407   0.88814    
## mutate:transtransversion         -0.844921   0.068774 -12.2855 < 2.2e-16 ***
## mutate:nonsyn                    -0.068704   0.090597  -0.7584   0.44824    
## mutate:comb_REFCG                 0.222616   0.091240   2.4399   0.01469 *  
## mutate:leading_minus_neighborC    0.162878   0.150907   1.0793   0.28044    
## mutate:leading_minus_neighborG    0.070650   0.132346   0.5338   0.59346    
## mutate:leading_minus_neighborT    0.166523   0.149278   1.1155   0.26462    
## mutate:leading_T                 -0.113063   0.099141  -1.1404   0.25411    
## mutate:leading_G                 -0.572744   0.097617  -5.8673 4.430e-09 ***
## ---
## Signif. codes:  0 '***' 0.001 '**' 0.01 '*' 0.05 '.' 0.1 ' ' 1
## 
## Log-Likelihood: -9098.4
```

While most of the results are broadly similar it is interesting to note that the effect of `comb_REFCG` has now increased (and is now significant) and the effect of `leading_G` is strongly negative. When we add `leading_G` the effect of `comb_REFCG` is now that for a leading C, and the effect of a leading G is the sum of `comb_REFCG` and `leading_G`. It seems that leading C’s increase the mutation rate but leading G’s decrease it. In the non-strand specific model leading C’s and G’s are equivalent and so the effect is missed because they largely cancel each other out. This has been noted in previous work (Foster et al. 2015).

We can also ask whether we would have detected differences between coding and non-coding regions and between synonymous and non-synonymous changes, had we not controlled for other properties that affect mutation (note codon usage bias is automatically controlled for in this analysis).

```
  m3 <- mlogit(new_seq ~ mutate+mutate:(coding_type+nonsyn)| 0 | 0, data=long_dat)
```

```
summary(m3)
```

```
## 
## Call:
## mlogit(formula = new_seq ~ mutate + mutate:(coding_type + nonsyn) | 
##     0 | 0, data = long_dat, method = "nr")
## 
## Frequencies of alternatives:choice
##       A       C       G       T 
## 0.24620 0.25424 0.25366 0.24591 
## 
## nr method
## 11 iterations, 0h:2m:46s 
## g'(-H)^-1g = 2.78E-09 
## gradient close to zero 
## 
## Coefficients :
##                                   Estimate Std. Error   z-value  Pr(>|z|)    
## mutate                           -9.118733   0.065949 -138.2697 < 2.2e-16 ***
## mutate:coding_typeprotein_coding -0.564088   0.100485   -5.6136 1.981e-08 ***
## mutate:nonsyn                    -0.191512   0.088875   -2.1549   0.03117 *  
## ---
## Signif. codes:  0 '***' 0.001 '**' 0.01 '*' 0.05 '.' 0.1 ' ' 1
## 
## Log-Likelihood: -9242.8
```

While the difference between coding and non-coding regions is very similar, the chance of seeing non-synonymous mutations has increased and is now significant. However, once the difference in transition/transversion rate is controlled for the deficit of non-synonymous mutations is reduced, and the estimated deficit is actually smaller than in the model in which full mutational spectrum is controlled for (model \(\textrm{m2}\)).

```
  m4 <- mlogit(new_seq ~ mutate+mutate:(coding_type+nonsyn+trans)| 0 | 0, data=long_dat)
```

```
summary(m4)
```

```
## 
## Call:
## mlogit(formula = new_seq ~ mutate + mutate:(coding_type + nonsyn + 
##     trans) | 0 | 0, data = long_dat, method = "nr")
## 
## Frequencies of alternatives:choice
##       A       C       G       T 
## 0.24620 0.25424 0.25366 0.24591 
## 
## nr method
## 11 iterations, 0h:2m:55s 
## g'(-H)^-1g = 1.18E-07 
## gradient close to zero 
## 
## Coefficients :
##                                   Estimate Std. Error   z-value  Pr(>|z|)    
## mutate                           -8.637458   0.073198 -118.0020 < 2.2e-16 ***
## mutate:coding_typeprotein_coding -0.699242   0.101033   -6.9209 4.486e-12 ***
## mutate:nonsyn                    -0.010615   0.090034   -0.1179    0.9061    
## mutate:transtransversion         -0.850988   0.068948  -12.3425 < 2.2e-16 ***
## ---
## Signif. codes:  0 '***' 0.001 '**' 0.01 '*' 0.05 '.' 0.1 ' ' 1
## 
## Log-Likelihood: -9167.2
```

Croissant, Yves. 2020. “Estimation of Random Utility Models in R: The mlogit Package.” *Journal of Statistical Software* 95 (11): 1–41.

Foster, Patricia L, Heewook Lee, Ellen Popodi, Jesse P Townes, and Haixu Tang. 2015. “Determinants of Spontaneous Mutation in the Bacterium Escherichia Coli as Revealed by Whole-Genome Sequencing.” *Proceedings of the National Academy of Sciences* 112 (44): E5990–99.

Lee, Heewook, Ellen Popodi, Haixu Tang, and Patricia L Foster. 2012. “Rate and Molecular Spectrum of Spontaneous Mutations in the Bacterium Escherichia Coli as Determined by Whole-Genome Sequencing.” *Proceedings of the National Academy of Sciences* 109 (41): E2774–83.
